# The cerebellar components of the human language network

**DOI:** 10.1101/2025.04.14.645351

**Authors:** Colton Casto, Moshe Poliak, Greta Tuckute, Hannah Small, Patrick Sherlock, Agata Wolna, Benjamin Lipkin, Anila M. D’Mello, Evelina Fedorenko

## Abstract

The cerebellum’s capacity for neural computation is arguably unmatched. Yet despite now ample evidence of cerebellar contributions to cognition, including language, its precise role in language processing remains debated. Here, we systematically characterize cerebellar language-responsive regions using precision fMRI. We identify four cerebellar regions that respond to language across modalities (Experiments 1a-b, n=754). One region—spanning Crus I/II/lobule VIIb—is selective for language relative to diverse non-linguistic perceptual, cognitive, and motor tasks (Experiments 2a-f, n=732), and the rest exhibit mixed-selective profiles, responding strongly to language but also to one or more of the non-linguistic conditions. Similar to the neocortical language system, the language-selective region is engaged by sentence-level meanings during comprehension and production (Experiments 3a-b, n=100) and shows fine-grained sensitivity to linguistic processing difficulty (Experiment 3c, n=5). Further, this region’s response to language is not due to the frequent presence of social content in language, as it is strongly engaged by both social and nonsocial sentences (Experiment 3d, n=10). Finally, all four regions, but especially Crus I/II/VIIb, are functionally connected to the neocortical language system (Experiment 4, n=85). We propose that these cerebellar regions constitute components of the extended language network, with one region supporting linguistic semantic processing and closely mirroring the selectivity of the neocortical language network, and the other three plausibly integrating information from diverse neocortical regions.

## Introduction

The cerebellum contains an impressive 80% of all neurons in the human brain (despite being only 10% of the size, Azevedo et al., 2009) and almost 80% of the surface area of the cerebral cortex (Sereno et al., 2020), making it one of the most computationally formidable neural structures. Historically, the cerebellum has been associated with motor control (Shadmehr et al., 2010; Manto et al., 2012), but decades of neuroimaging and neuropsychological research have now established that the cerebellum also contributes to—and may have even evolutionarily enabled (Leiner et al., 1986; 1993; Balsters et al., 2010; Kochiyama et al., 2018; Magielse et al., 2023)—several cognitive functions, including the uniquely human capacity for language (Peterson et al., 1989; Stoodley & Schmahmann, 2009a; Murdoch, 2010; Stoodley et al., 2012; Mariën et al., 2014; Gatti et al. 2021a; 2023; LeBel & D’Mello, 2023).

The cerebellum contains a mosaic of functionally distinct subregions (Stoodley & Schmahmann, 2009a; Buckner et al., 2011; Stoodley et al., 2012; King et al., 2019; Nettekoven et al., 2024; Saadon-Grossman et al., 2024), which plausibly result from reciprocal closed-loop connections with distinct contralateral neocortical regions (Middleton & Strick, 2001; Kelly & Strick, 2003). This rich organization suggests that different cerebellar subregions may contribute to cognition in distinct ways. However, the precise contributions of these different regions to cognition remain debated. For example, many studies have implicated the right posterior cerebellum in language processing (e.g., Crus I/Crus II; Peterson et al., 1989; Stoodley & Schmahmann, 2009a; 2010; Stoodley et al., 2012; Mariën et al., 2014; Guell et al., 2018a; 2018b; King et al., 2019; Gatti et al., 2020a; 2020b; Li et al., 2022; Nakatani et al., 2023, Turker et al., 2023; Nettekoven et al 2024), but no consensus has emerged about what linguistic computations this region supports. In neuroimaging research, activations in the right posterior cerebellum have been reported for diverse language paradigms, including verb generation (Frings et al., 2006; Stoodley et al., 2012), sentence completion (Moberget et al., 2014; D’Mello et al., 2017), and passive comprehension of naturalistic passages (King et al., 2019; Guell et al., 2018; LeBel et al., 2021). Similarly, damage to the right posterior cerebellum can lead to diverse linguistic deficits (Mariën et al., 2001; de Smet et al., 2007; Satoer et al., 2024), including grammatical difficulties (Silveri et al., 1994; Justus, 2004), dysprosodia (Schmahmann & Sherman, 1998), anomia (Schmahmann & Sherman, 1998), alexia (Moretti et al., 2002), reduced verbal fluency (phonological and semantic; Leggio et al., 2000), and ‘metalinguistic’ deficits (Guell et al., 2015). Tying all these diverse findings into a single coherent story of cerebellar contributions to language has proven challenging.

Progress in cerebellar language research has been slowed by several recurring issues—some that are prevalent in language neuroscience generally, and others that are specific to cerebellar studies. First, similar to early work on neocortical language regions, many tasks that have been commonly used to study language in the cerebellum do not isolate language from non-linguistic perceptual, motor, and cognitive processes (all of which draw on brain regions distinct from the language system in the neocortex; see Fedorenko et al., 2024 for a review). For example, paradigms such as verb generation (e.g., Frings et al., 2006) or verbal fluency (e.g., Schlosser et al., 1998) conflate linguistic processing with general task demands, and many language paradigms lack controls for low-level perceptual and/or motor processes, which makes it challenging to unambiguously interpret the observed neural responses. Second, many past cerebellar brain imaging and lesion studies have relied on anatomical localization and averaging across subjects. These approaches have obscured the fine-grained, individually-varying functional organization that can be revealed through precision fMRI (Gordon et al., 2017; Braga & Buckner, 2017; Fedorenko & Blank, 2020; DiNicola & Buckner, 2020). The limited use of within-participant localization approaches makes it difficult to assess whether different studies observe effects in the ‘same’ region (see Brett et al., 2002 and Saxe et al., 2006 for early discussions). For example, prior studies have reported activation in the right posterior cerebellum for language processing but also for other aspects of cognition (e.g., theory of mind and executive processing: Desmond & Fiez, 1998; Bellebaum & Daum, 2007; Stoodley & Schmahmann, 2009a; Stoodley et al., 2012; E et al., 2014; Marek et al., 2018; Guell et al., 2018a; D’Mello et al., 2020a; Van Overwalle, 2024). Without performing within-individual comparisons, it is difficult to determine whether the cerebellar language regions overlap with these regions supporting non-linguistic functions, or whether they are distinct but nearby (for similar issues in the neocortex, see: Scholz et al., 2009; Braga & Buckner, 2017; Fedorenko & Blank, 2020; DiNicola & Buckner, 2020).

Research on language processing in the cerebellum, in particular, has typically only examined responses in a single or a small number of paradigms, leaving open many questions about the underlying computations. For example, some studies have evaluated specific hypotheses about the cerebellum’s contribution to language (e.g., linguistic prediction: Moberget et al., 2014; Lesage et al., 2017, D’Mello et al., 2017)—often grounded in circuit-level models of the cerebellum (Marr, 1969; Albus, 1971; Ito & Kano, 1982)—but have not tested the relevant regions’ responses to other aspects of language or to non-linguistic tasks. As a result, these studies do not tell us whether the region in question implements linguistic prediction vs. language processing more generally, nor whether its computations are language-specific vs. domain general. Other studies have attempted to understand the overall organization of the cerebellum by using multi-domain task batteries (e.g., King et al., 2019; Nettekoven et al., 2024) or large amounts of naturalistic-cognition data (e.g., resting state data; Habas et al., 2009; Xue et al., 2021; Saadon-Grosman et al., 2024; see Buckner et al., 2011 for earlier work based on group-level data). Such studies identify cerebellar regions that are distinct from one another but can inform function within a given domain only coarsely.

Finally, neocortical and cerebellar language regions have predominantly been studied separately, with the vast majority of previous studies focusing on the neocortex, leaving the role of the linguistic cerebellum *relative* to the neocortical circuits underspecified (see e.g., Shashahani et al., 2024 for a recent proposal of how to systematically test for functional differences between the neocortex and the cerebellum). For example, linguistic computations hypothesized to be supported by the cerebellum (most notably, linguistic prediction, as mentioned above; Sokolov et al., 2017; Moberget & Ivry, 2016; Argyropoulos, 2016; Gatti et al., 2021b) have also been shown to be performed by the neocortical language regions (Lopopolo et al. 2017; Shain, Blank et al. 2020; Heilbron et al., 2022). The fragmentation of language neuroscience into cerebral and cerebellar research has minimized collective theorizing, which is crucial for developing a comprehensive account of how the brain processes language (LeBel & D’Mello, 2023; Wang et al., 2025).

Here, we adopted an approach that has proven productive in illuminating the function of the neocortical language system (see Fedorenko et al., 2024 for a review) to shed light on the cerebellar contributions to language processing. In particular, using a large-scale fMRI dataset (n=846 unique participants, n=1,033 scanning sessions, n=26 experiments), we i) used a well-established functional localizer for language (Fedorenko et al., 2010; Lipkin et al., 2022)—which effectively isolates language processing from non-linguistic perceptual, motor, and cognitive processes—to search for language-responsive regions in the cerebellum. Then, we characterized these language-responsive regions—functionally defined in individual participants in each experiment—in terms of ii) their selectivity for language relative to other functions and, for language-selective regions, iii) the nature of their linguistic contributions. Finally, we iv) compared the similarity and functional connectivity of the cerebellar language regions to the neocortical language network. To foreshadow our critical results, we identified four regions in the right posterior cerebellum that respond to language both during reading and listening. One of these regions shows strong selectivity for language relative to diverse non-linguistic tasks and appears to process sentence-level meanings during comprehension and production. Relative to other cerebellar language regions, this region is functionally most similar to and shows the strongest functional connectivity with the neocortical language network. The remaining three cerebellar language-responsive regions all show mixed selectivity, suggesting they may integrate linguistic and non-linguistic information from multiple neocortical regions.

## Results

### 1) Four distinct regions in the right posterior cerebellum respond robustly to language across modalities

We first investigated the topography of language responses in the cerebellum across a large cohort of participants (Expt. 1a, n=754, data from Lipkin et al., 2022). All participants completed a language ‘localizer’ paradigm that involved reading sentences and lists of nonwords (which were perceptually similar but did not contain linguistic meaning or structure (**Fig. 1a**, Fedorenko et al., 2010; Lipkin et al., 2022). The Language (sentences) > Control (nonwords) contrast targets brain areas that support computations related to retrieving words from memory, syntactic structure building, and semantic composition, and generalizes robustly across modalities, tasks, and languages (e.g., Fedorenko et al., 2024 for a review).

**Figure 1:**
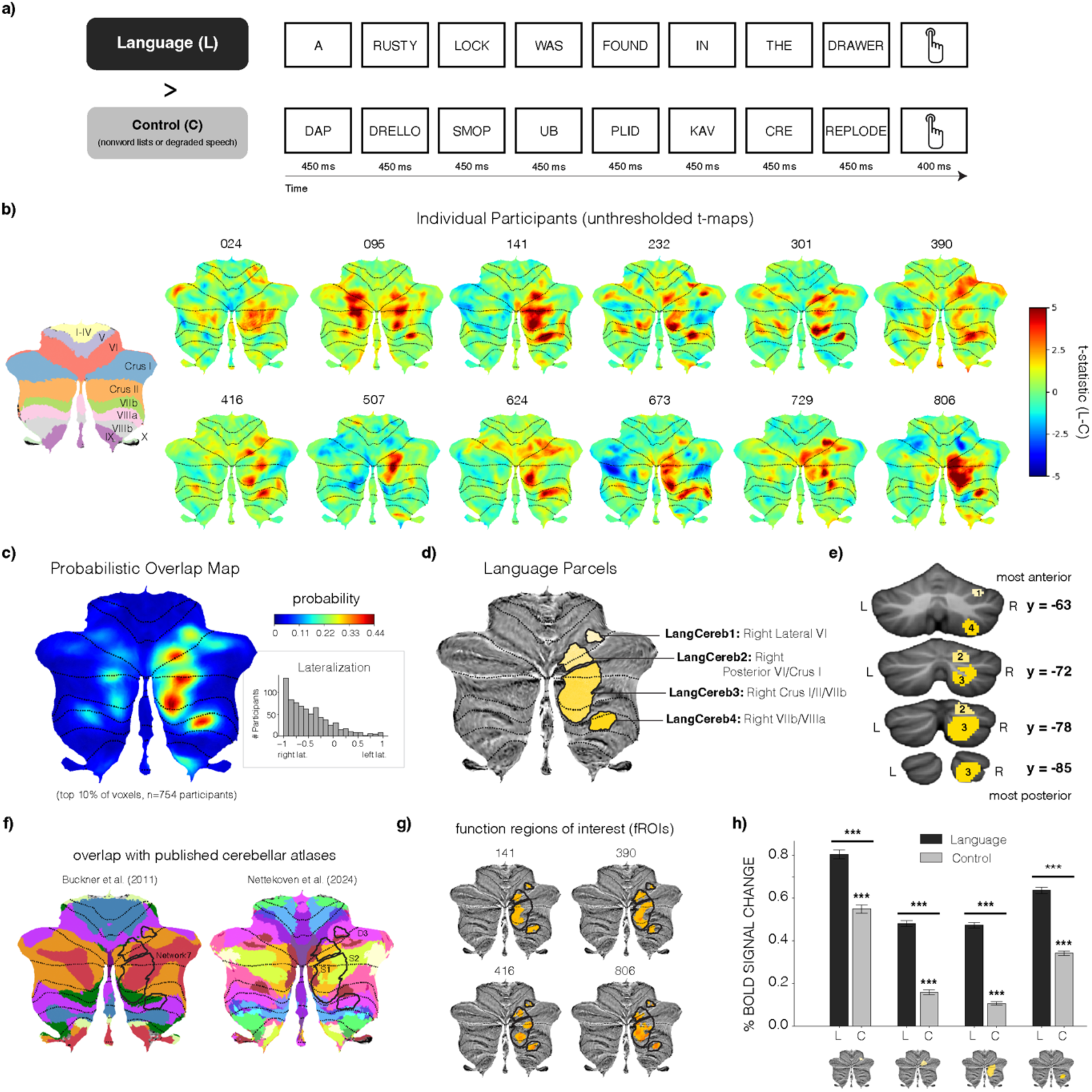
Language-responsive regions of the cerebellum. **a)** Language localizer paradigm. Participants read sentences (language, L) or non-word lists (a perceptually-matched control condition, C) presented one word/nonword at a time. **b)** Left: A cerebellar atlas showing cerebellar lobules visualized on a cerebellar flatmap (used for most visualizations in the paper). Right: Cerebellar responses to language in 12 individual participants. The maps are unthresholded t-statistic maps for the Language>Control contrast; cerebellar lobule boundaries are marked in black. **c)** Probabilistic overlap map across n=754 participants constructed from individual binarized Language>Control maps. Individual maps have a value of 1 for the top 10% of language-responsive voxels across the whole brain and 0 otherwise. Color shading of the overlap map reflects the proportion of participants for whom that voxel was in the top 10% of their most language-responsive voxels. *Inset:* Lateralization of cerebellar language responses within individual participants, computed as the difference between the number of significant language-responsive voxels (uncorrected t > 2.576, p<0.01, two-sided t-test) in the left cerebellum minus the number of significant voxels in the right cerebellum, over the sum of the total number of significant voxels in the cerebellum (negative values indicate a right-hemispheric bias). **d)** Parcellation of the probabilistic overlap map via the group-constrained subject-specific (GSS) analysis which identifies spatially consistent areas of activation across participants (<u>Methods</u>). Four parcels were identified, all falling in the right posterior cerebellum. **e)** The parcels visualized in volume space shown on four coronal slices. **f)** Overlap of our language parcels with two published cerebellar atlases: Buckner et al. (2011) and Nettekoven et al. (2024) (see **Suppl. Table 2** for a quantitative comparison of the overlap). LangCereb3 (Crus I/II/VIIb) and LangCereb2 (posterior VI/Crus I) overlap substantially with the S1R and S2R (i.e., “sociolinguistic”) regions reported in Nettekoven et al. (2024) and the DMN region (i.e., Network7) in the Buckner et al. (2011) parcellations. The other cerebellar language regions (LangCereb1 and 4) primarily overlap with the D3R (i.e., “demand”) region from the Nettekoven et al. (2024) parcellation and the Network 4 and 6 regions from the Buckner et al. (2011) parcellations. We would like to emphasize that although the locations of our parcels are largely consistent with where others have reported cerebellar language activations, our parcels are designed to be used in conjunction with functional localization within individuals. The parcels are, by design, relatively large, and as a result, using the full set of voxels in each parcel would drastically underestimate language selectivity and make it more difficult to decipher the linguistic areas’ computations. **g)** Illustration of the procedure used to define fROIs within individual participants (shown here for four sample participants). Parcel outlines are marked in black, and the most language-responsive voxels (i.e., the top 10%) within a given parcel are marked in orange. Responses to the conditions in all experiments are reported for these individually-defined subsets of voxels. **h)** Responses to the Language and Control conditions in each of the four fROIs (estimated using different runs to localize the regions and measure their responses to language; <u>Methods</u>).

To search for language-responsive regions within the cerebellum, we used a group-constrained subject-specific approach (‘GSS’, see Fedorenko et al., 2010 and <u>Methods</u>). This approach is similar to traditional random-effects group analyses in that it searches for areas of spatially consistent activation across participants, but critically, it allows inter-individual differences in the precise locations of functional regions. This is especially important in the cerebellum, where language responses vary substantially in their topography across participants (**Fig. 1b**; see OSF https://osf.io/y5t46/ for activation maps from all participants; see **Suppl. Fig. 1a** for evidence that language responses in the cerebellum are more topographically variable than in the neocortex). Nonetheless, there is sufficient consistency in the location of cerebellar language responses (**Fig. 1c**), such that a whole-brain GSS analysis successfully recovered *four language regions,* all in the right cerebellum (**Fig. 1d-e**). The right-hemisphere bias was also apparent when looking at language-responsive voxels across the cerebellum (**Fig. 1c inset**; see **Suppl. Fig. 1b** for evidence of a strong negative correlation between cerebellar and neocortical lateralization).

All four language regions are located in the posterior portion of the right cerebellum (ordered from most superior to most inferior): i) the smallest region located in lateral lobule VI (LangCereb1); ii) a region spanning posterior lobule VI and Crus I (LangCereb2); iii) the largest region spanning Crus I, Crus II and lobule VIIb (LangCereb3); and iv) a region spanning lobules VIIb and VIIIa (LangCereb4; **Fig. 1d-e**; the GSS analyses applied exclusively to the cerebellum yielded similar parcels, but additionally yielded a region in right lobule IX—LangCereb5—and four regions in the left cerebellum, **Suppl. Fig. 2a**). The locations of these regions overlap partially with where language regions have been reported in prior studies (e.g., Buckner et al., 2011; Nettekoven et al., 2024; **Fig. 1f**, **see Suppl. Table 2** for a quantitative comparison of this overlap). We defined individual functional regions of interest (fROIs) in each participant (**Fig. 1g**) and tested responses to language (**Fig. 1h**) using an independent portion of the data (e.g., using half of the runs to identify the fROIs and the other half to measure the responses in those regions). All four regions exhibited stronger responses to language than to the control condition (**Fig. 1h**; ps<0.001 evaluated with a linear mixed-effects model, <u>Methods</u>; see **Suppl. Table 1** for complete statistical results). The responses to the Language condition were more than two times higher than to the Control condition in LangCereb2 and LangCereb3with more moderate effect sizes in the other regions (**Fig. 1h**).

For spoken (cf. signed) languages, meaning can be extracted through reading (when linguistic input is written) or listening (when linguistic input is spoken). In the neocortex, the same network supports language comprehension in both modalities (e.g., Vagharchakian et al., 2012; Regev et al., 2013; Scott et al., 2017; Deniz et al., 2019; Malik-Moraleda, Ayyash et al., 2022). Therefore, we next tested whether cerebellar language responses also generalize across modalities. A subset of participants in Expt. 1a also completed a listening-based localizer (Expt. 1b, n=85, originally published in Malik-Moraleda, Ayyash et al., 2022) in which responses to short passages from *Alice in Wonderland* were contrasted with acoustically degraded versions of those passages (where the content was no longer intelligible, <u>Methods</u>). The topography of cerebellar responses during reading and listening was qualitatively similar both within individual participants (Fig. 2a, see OSF https://osf.io/y5t46/ for activation maps from all participants) and at the group level (Fig. 2b; Pearson correlation between reading and listening probabilistic overlap maps = 0.78; though we did find a statistical difference between two modalities in their fine-grained patterns of activation in all regions, Fig. 2c**-g**, ps<0.05, one sample Wilcoxon signed-rank test). In line with the coarse topographic similarity of language responses across modalities—and similar to the neocortical language network (Fig. 2c)—all four cerebellar language regions showed robust responses both during reading and listening relative to the respective control condition (Fig. 2d**-g**, **Suppl. Table 3**; see Suppl. Fig. 2b for responses during reading vs. listening in right lobule IX and Suppl. Fig. 3 for responses in the left cerebellar homotopic regions). In addition, LangCereb1 (Lateral VI) showed an overall preference for written stimuli (Fig. 2d, **Suppl. Table 3**), and LangCereb3 (Crus I/II/VIIb)—for spoken stimuli (Fig. 2f, **Suppl. Table 3**, <u>Methods</u>). Together, these analyses suggest that cerebellar language regions are engaged in the abstract, modality-independent processing of linguistic input, similar to their neocortical counterparts (see also Expt. 3d below).

**Figure 2:**
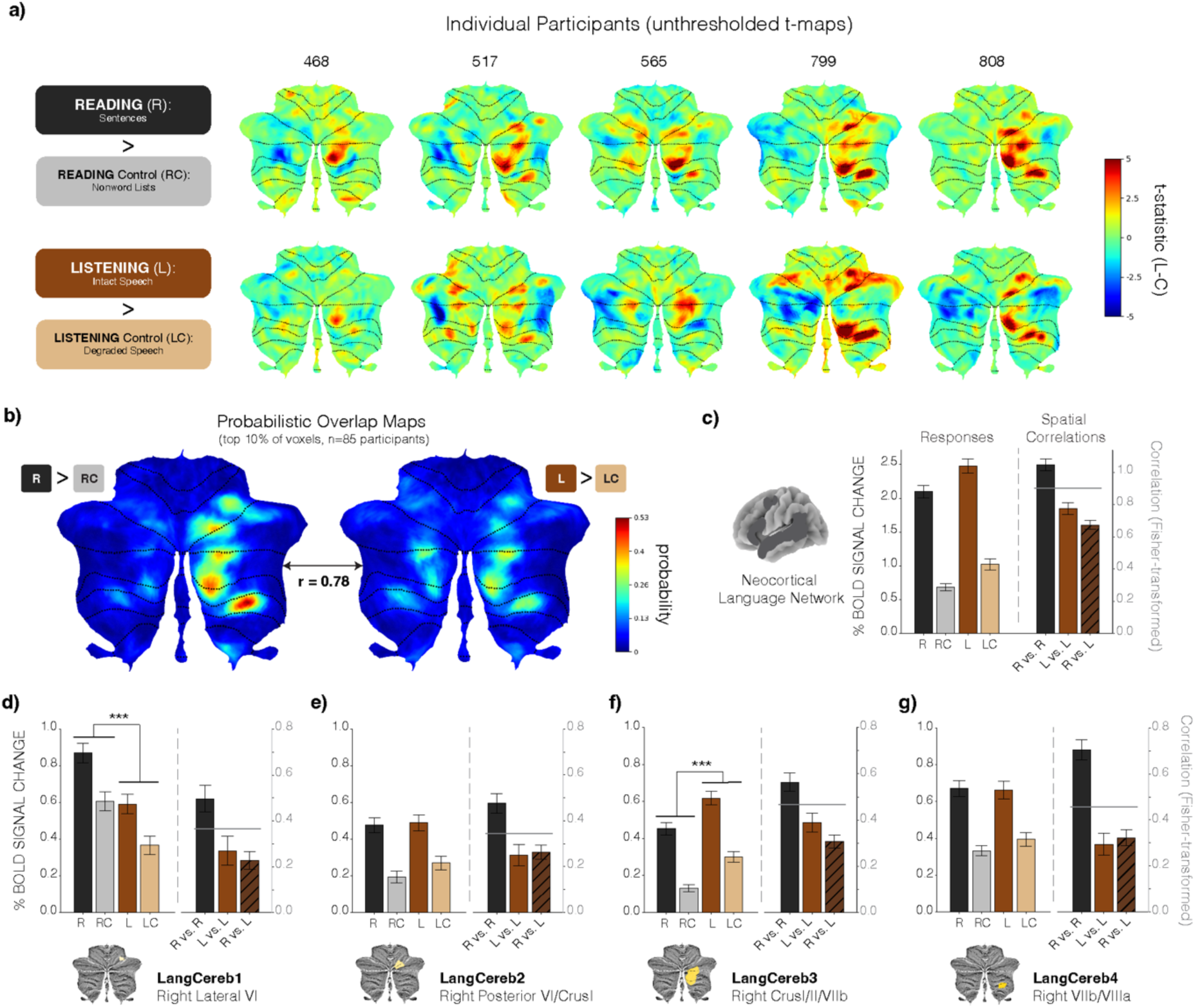
Engagement of cerebellar language regions during reading and listening. **a)** Cerebellar responses during reading (R) versus a visual control condition (RC) (top) and during listening (L) versus an auditory control condition (LC) (bottom) in five sample participants. The maps are un-thresholded t-statistic maps for the Language>Control contrast in each modality. **b)** Probabilistic overlap maps across n=85 participants constructed from individual binarized maps for the reading contrast (top) vs. the listening contrast (bottom). **c-g)** Responses during reading and listening (left) and spatial correlations between reading and listening (right) in the neocortical language network (**c**) and the four cerebellar language regions (localized using the reading-derived parcels; **d-g**). Horizontal grey lines over the spatial correlations reflect the geometric mean of the reading-reading and listening-listening correlations for a given region—the value that we would expect the reading-listening correlations of a region to be if the activation patterns were exactly the same. The reading-listening correlations were significant lower than this value in all regions (ps>0.05, one sample Wilcoxon signed-rank test). *Left:* All of the cerebellar language regions respond robustly during both reading and listening (over and above their respective perceptually-matched control conditions), with LangCereb1 showing a preference for visual stimuli and LangCereb3 showing a preference for auditory stimuli (evaluated with a linear mixed-effects model, <u>Methods</u>; **Suppl. Table 3**). *Right:* All cerebellar language regions exhibit robust spatial correlations between patterns of activation to reading and listening, however, the correlations are less than would be expected if the patterns were identical.

### 2) One cerebellar language region (LangCereb3) shows selectivity for language, and the remaining three show mixed-selectivity profiles

Having identified the regions of the cerebellum that are recruited during language processing, we next tested whether these regions respond selectively to language, which can help inform their function(s). In particular, if a region only responds to language, this would suggest that it supports specifically linguistic computations; if, on the other hand, a region responds to both language and some non-linguistic condition(s), this would suggest that the region supports multiple functions, or implements a computation that is shared across the conditions that it responds to (for hypotheses about language sharing computational demands with domains such as music, non-verbal semantics, and social perception see e.g., Lerdahl & Jackendoff, 1977; Krumhansl & Keil, 1982; Talmy, 2000; Binder et al., 2009; Kendon, 1967; McNeill, 1992; Novack & Goldin-Meadow, 2017). We therefore examined the cerebellar language regions’ responses to a suite of non-linguistic conditions (Expts. 2a-e).

We organized the battery of non-linguistic conditions into five domains. First, given the well-documented importance of the cerebellum in motor control (Shadmehr et al., 2010; Manto et al., 2012) and speech production in particular (Ackermann et al., 2007; Ackermann, 2008), we examined the engagement of the cerebellar language regions during ***speech articulation***. We instructed participants (n=33, Expt. 2a, Wolna et al., in prep.) to overtly recite a syllable sequence “ba-ga-ra-da” or tap a sequence using their fingers (see Suppl. Fig. 4a for example stimuli for all tasks; see **Suppl. Table 43** for links to all experimental stimuli; see <u>Methods</u> for design details for all tasks). Neocortical language regions respond only weakly to these motor conditions (not significantly above the response to the control condition from the language localizer, evaluated here and elsewhere with a linear mixed-effects model, <u>Methods</u>; see **Suppl. Tables 4-7** for complete statistical results for the neocortical language regions; Fig. 3a). Similarly, in cerebellar language regions, we found that LangCereb2 (posterior VI/Crus I) and LangCereb3 (Crus I/II/VIIb) also responded only weakly to these motor conditions (ps>0.05 against the fixation baseline; see **Suppl. Tables 12-15** and **16-19** for complete statistical results for LangCereb2 and LangCereb3, respectively; Fig. 3c, d; see Suppl. Fig. 5 for results from homotopic versions of the right cerebellar language regions, and see Suppl. Fig. 2c for results from the language region in right lobule IX). In contrast, the language regions in lateral VI (LangCereb1) and VIIb/VIIIa (LangCereb4) responded at least as strongly to the Articulation and Finger Tapping conditions (and not differentially so, ps>0.05; **Suppl. Tables 7, 11, 15, 19, 23**) as during sentence comprehension (ps>0.05; see **Suppl. Tables 8-11** and **20-23** for complete statistical results for LangCereb1 and LangCereb4, respectively; Fig. 3b, e). These results show that certain cerebellar language regions (LangCereb2 and LangCereb3) are selective for language relative to articulatory processes, but other regions (LangCereb1 and LangCereb4) are recruited during both language and speech/motor tasks.

**Figure 3:**
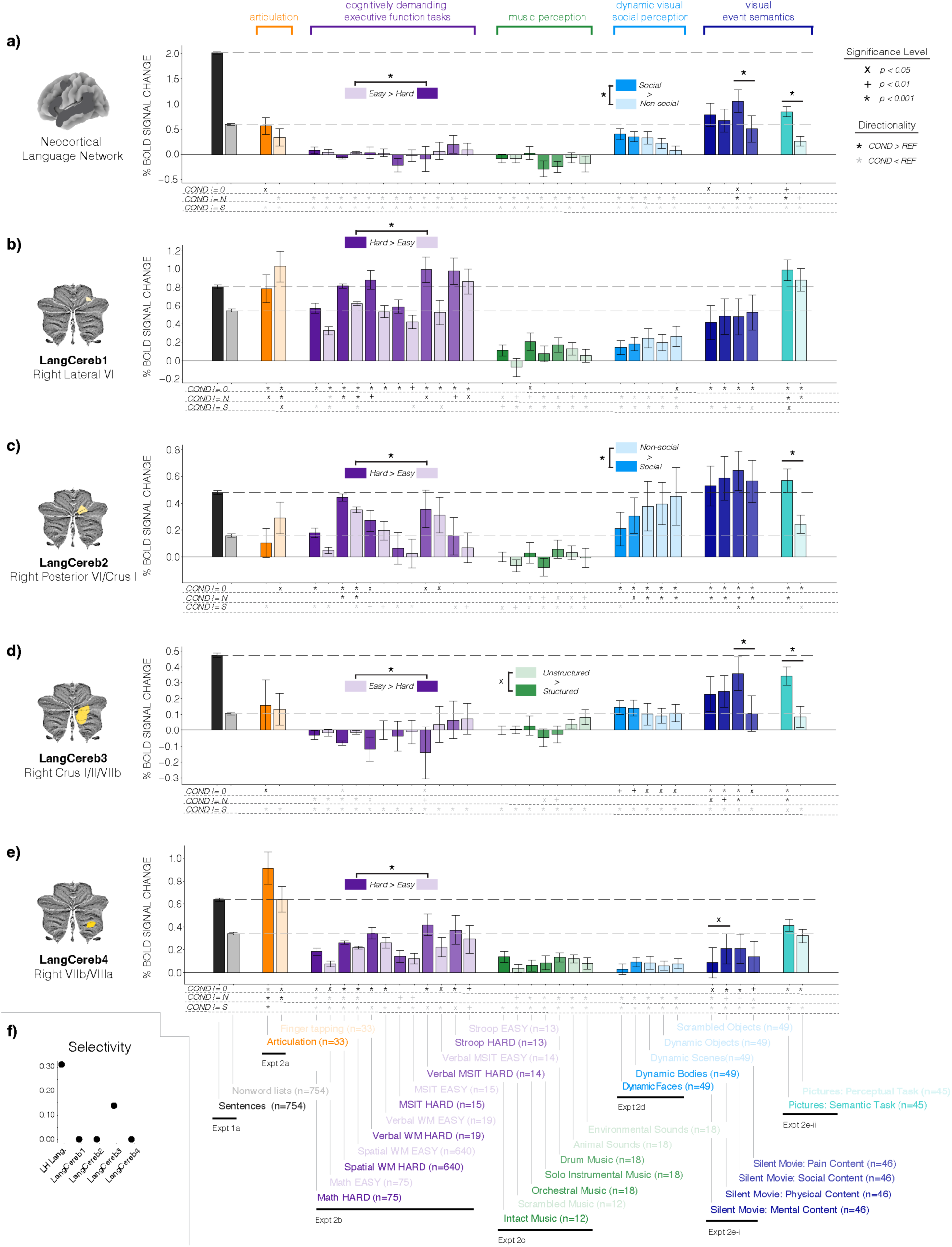
Selectivity of cerebellar language regions for language over non-linguistic tasks. Neocortical and cerebellar responses to a suite of non-linguistic tasks organized into five domains. **a-e)** Responses to the language localizer are shown in black and gray (Expt. 1a, n=754, data from Lipkin et al., 2022). Conditions that evaluate the cerebellum’s role in speech articulation are shown in orange (Expt. 2a, n=33, data from Wolna et al., in prep). Conditions that evaluate the cerebellum’s role in cognitively demanding executive function tasks are shown in purple (Expt. 2b-i, n=75, data from Malik-Moraleda, Ayyash et al., 2022; Expt. 2b-ii, n=640, data from Fedorenko et al., 2011; Lipkin et al., 2023; Expt. 2b-iii, n=19, data from Fedorenko et al., 2011 for this and all subsequent executive function tasks; Expt. 2b-iv, n=15; Expt. 2b-v, n=14; Expt. 2b-vi, n=13). Conditions that evaluate the cerebellum’s role in music perception are shown in green (Expt. 2c-i, n=12, data from Fedorenko et al., 2011; Expt. 2c-ii, n=18, data from Chen et al., 2023). Conditions that evaluate the cerebellum’s role in dynamic visual social perception are shown in light blue (Expt. 2d, n=49, data from Pritchett et al., 2018). Conditions that evaluate the cerebellum’s role in visual event semantics are shown in dark blue (Expt. 2e-i, n=46, data from Shain, Paunov, Chen et al., 2023) and teal (Expt. 2e-ii, n=45, data from Ivanova et al., 2021; Ivanova, 2022). Across all experiments, control conditions are denoted by lightly shaded bars. Responses reflect the percent BOLD signal change in the individually-defined language fROIs (<u>Methods</u>). All error bars reflect the standard error of the mean over participants for the given condition. The dashed black line indicates the response of a given region to language; the dashed gray line indicates the response of a given region to the control condition from the language localizer. **a)** Neocortical responses averaged over the five ‘core’ left-hemisphere language regions (orbital inferior frontal gyrus, inferior frontal gyrus, middle frontal gyrus, anterior temporal lobe, and posterior temporal lobe; see <u>Methods</u>). **b-e)** Responses in the four cerebellar language regions defined in Fig. 1d. **f)** Selectivity for language in the neocortical language regions and the cerebellar language regions. ‘Classic’ selectivity refers to the difference between a region’s response to language and the next highest non-linguistic condition. The entropy-based measure of selectivity is computed over the entire response profile (<u>Methods</u>).

Second, given the previous reports of cerebellar engagement in executive function tasks (Desmond & Fiez, 1998; Bellebaum & Daum, 2007; Stoodley & Schmahmann, 2009; Stoodley et al., 2012; Guell et al., 2018; D’Mello et al., 2020a), and executive function deficits in cerebellar patients (Schmahmann & Sherman, 1998), we examined the engagement of the cerebellar language regions during ***cognitively-demanding*** tasks. We presented participants with several commonly used tasks, each with a hard and an easy condition, including an arithmetic addition task (Expt. 2b-i, n=75, Suppl. Fig. 4b, data from Malik-Moraleda, Ayyash et al., 2022), a spatial working memory task (Expt. 2b-ii, n=640, Suppl. Fig. 4c, data from Fedorenko et al., 2011; Lipkin et al., 2023), a verbal working memory (digit span) task (Expt. 2b-iii, n=19, Suppl. Fig. 4d, for this and the rest of experiments in this set, the data come from Fedorenko et al., 2011), a multi-source interference task (MSIT, Bush & Shinn, 2006; Expt. 2b-iv, n=15, Suppl. Fig. 4e), a verbal multi-source interference task (vMSIT, Expt. 2b-v, n=14, Suppl. Fig. 4f), and a Stroop task (Expt. 2b-vi, n=13, Suppl. Fig. 4g).

As shown in prior work (Fedorenko et al., 2011, 2012; Malik-Moraleda, Ayyash et al., 2022), the neocortical language regions do not respond above baseline to any of these tasks (ps>0.05, **Suppl. Table 4**, Fig. 3a). Within the cerebellum, LangCereb3 (Crus I/II/VIIb) showed a similar profile (ps>0.05, **Suppl. Tables 16-19**, Fig. 3d). However, the other cerebellar language regions were engaged to some degree during many of the cognitively demanding tasks (ps<0.05; **Suppl. Tables 4**, **8**, **16**; Fig. 3b, c, e), and generally responded more to the Hard than the Easy condition (**Suppl. Tables 11, 15, 23**; Fig. 3b, c, e). This profile is reminiscent of the domain-general Multiple Demand (MD) neocortical network (Duncan, 2010; Fedorenko et al., 2013; Assem et al., 2020; Duncan et al., 2020). However, in contrast to the neocortex, where the language and MD networks do not overlap (see Fedorenko & Blank, 2020 for review), three of the cerebellar language regions showed mixed selectivity, with high responses during both language comprehension and cognitively-demanding tasks.

Third, given a) the long-standing parallels that have been drawn between language and music, especially concerning their ***hierarchical structure*** (e.g., Lerdahl & Jackendoff, 1977), b) previous reports of cerebellar contributions to music processing (Evers & Tölgyesi, 2022), and c) the critical role of prediction—a hypothesized core function of the cerebellum (e.g., Sokolov et al., 2017)—for music processing (Huron, 2006), we examined the responses in cerebellar language regions to music. Across two experiments, participants listened to instrumental music clips along with various auditory control conditions that lack hierarchical structure (Expt. 2c-i, n=12, data from Fedorenko et al., 2011; and Expt. 2c-ii, n=18, data from Chen et al., 2023). Neocortical language regions do not respond above baseline to music (ps>0.05, **Suppl. Table 4**, Fig. 3a), nor do they show sensitivity to music structure (ps>0.05, **Suppl. Table 7**, Fig. 3a; Fedorenko et al., 2011; Chen et al., 2023). The cerebellar language regions all mirrored this pattern of response (ps>0.05, **Suppl. Tables 8-23**, Fig. 3b**-e**).

Fourth, because language is a ***communicative*** signal, typically present in social contexts, we examined responses of cerebellar language regions to non-linguistic social stimuli that can be used to communicate information (Expt. 2d, n=49, data from Pritchett et al., 2018). Participants watched videos of faces and bodies, as well as non-social stimuli (e.g., inanimate objects; Suppl. Fig. 3h). Neocortical language regions respond only weakly to these conditions (not significantly above the response to the control condition from the language localizer, ps>0.05, **Suppl. Table 5**, Fig. 3a; Pritchett et al., 2018). Similarly, all cerebellar language regions except LangCereb2 (posterior VI/Crus I) responded weakly to these conditions (ps>0.05; **Suppl. Tables 9**, **17**, **21**; Fig. 3b, d, e), and none of these three showed a stronger response to the social conditions than the non-social conditions (ps>0.05; **Suppl. Tables 11**, **19**, **23**; Fig. 3b, d, e). LangCereb2 (posterior VI/Crus I) responded quite strongly to these visual conditions (**Suppl. Tables 12-14**, Fig. 3c) with a stronger response to the non-social conditions than the social conditions (p<0.001, **Suppl. Table 15**, Fig. 3c).

Lastly, because language carries ***meaningful information***, we examined the responses of cerebellar language regions to non-verbal meaningful stimuli. In one experiment, participants watched a silent movie (Pixar’s *Partly Cloudy*, Expt. 2e-i, n=46, Suppl. Fig 4i, data from Shain, Paunov, Chen et al., 2023). Segments of the movie were annotated for different types of events (e.g., events that involved social interactions, events that involved physical pain, etc.). In another experiment, participants watched static images of events (e.g., a jester entertaining a king, a shark biting a man; Expt. 2e-ii, n=45, Suppl. Fig 4i, data from Ivanova et al., 2021 and Ivanova, 2022) and either performed a semantic judgment (e.g., deciding whether the event is plausible) or a difficulty-matched perceptual judgment (e.g., deciding whether the image is moving to the left or to the right; this experiment also included a linguistic version of the events, Suppl. Fig. 6, <u>Methods</u>).

Neocortical language regions respond a moderate amount to visual semantic stimuli (significantly above the response to the Control condition from the language localizer, ps<0.01 for 3 of the 5 semantic conditions, **Suppl. Table 5**), but significantly below the response to language (ps<0.001 for all semantic conditions, **Suppl. Table 6**, Fig. 3a; Ivanova et al., 2021; Ivanova, 2022; Shain, Paunov, Chen et al., 2023; Sueoka, Paunov et al., 2024). In the cerebellum, LangCereb1 (lateral VI) and LangCereb4 (VIIb/VIIIa) mirrored this pattern, with substantially stronger responses during language processing than the visual semantic conditions (p<0.05; **Suppl. Tables 9-10, 21-22**; Fig. 3b, e). LangCereb3 (Crus I/II/VIIb) also responded more strongly to language than to visual semantics (ps<0.05, **Suppl. Table 18**, Fig. 3d), although it did show a stronger response to the semantic task than the perceptual task in Expt 2e-ii, ruling out explanations in terms of low-level visual processing or task demands (p<0.001; **Suppl. Table 19**, Fig. 3d). LangCereb3 is also the only cerebellar language region that mirrored the neocortical language network in showing a significantly greater effect to the social-interaction condition than the physical-pain condition in Expt. 2e-i (p<0.001, **Suppl. Table 19**). The remaining region— LangCereb2 (posterior VI/Crus I)—responded similarly strongly to language and visual semantics (ps>0.05, **Suppl. Table 14**, Fig. 3c), and, similar to LangCereb3, showed a stronger response to the semantic task than the perceptual judgement task (ps<0.001; **Suppl. Table 15**; Fig. 3c). These results suggest that LangCereb2 and 3 may process semantic information across both verbal and non-verbal formats, although LangCereb3, similar to the neocortical language network, still exhibits a preference for verbal semantic information.

Having examined the responses of the cerebellar language regions to diverse non-linguistic conditions, we pursued two additional analyses to quantify i) the overall selectivity of these regions for language and ii) the degree of overlap with nearby cognitive and motor networks. First, we evaluated the overall selectivity of the cerebellar language regions (and the neocortical language regions, for comparison) across the entire suite of non-linguistic conditions. For a region to be ‘selective’ for language, it would need to be engaged much more strongly by language than by non-linguistic tasks—if a region is engaged as strongly by, for example, the spatial working memory task (Expt. 2b-ii) as it is for language (regardless of whether it shows an effect of Hard > Easy), then it would be challenging to argue that this region is engaged in a specifically-linguistic computation. In particular, we defined selectivity as the difference between a region’s response to language and its response to the next highest non-linguistic condition (scaled by sum of the two responses) (**Suppl. Table 24**, Fig. 3f; <u>Methods</u>). The neocortical language network was more selective for language than any of the cerebellar language regions, responding two times stronger to the preferred condition (language) than to any of the non-preferred (non-linguistic) conditions (S=0.311, **Suppl. Table 24**, Fig. 3f)—similar to the degree of selectivity reported in category-selective visual areas (e.g., Kanwisher et al., 1997). CerebLang3 (Crus I/II/VIIb) was the most selective of the cerebellar language regions, though it was slightly less selective than the neocortical language network (S=0.14, **Suppl. Table 24**, Fig. 3f). This finding aligns with the visually observable similarity of its response profile to the neocortical language network’s profile. Selectivity for the remaining cerebellar language regions was at zero (i.e., at least one non-linguistic condition elicited a response at or above the level of its response to language) (**Suppl. Table 24**, Fig. 3f). Thus, although these regions responded strongly and consistently across modalities to language, they were not selective for language. LangCereb1 (lateral VI) and LangCereb4 (VIIb/VIIIa) were recruited for language and for non-linguistic cognitively demanding tasks (Fig. 3b, e), with LangCereb4 also showing engagement during speech articulation and hand motor tasks (Fig. 3e), and LangCereb2—for processing meaningful visual stimuli (Fig. 3c). These findings suggest that these three regions do not carry out exclusively linguistic computations and merit additional experimentation to further characterize their function(s) (see <u>Discussion</u>).

Second, in light of observations that cognitive networks within the cerebellum are highly interdigitated (e.g., Saadon-Grossman et al., 2024; Nettekoven et al., 2024), and given the high degree of functional heterogeneity in some of our response profiles (Fig. 3), we evaluated the degree to which the cerebellar language regions are dissociable from three nearby cognitive and motor-control networks: the Theory of Mind (ToM) network (Saxe & Kanwisher, 2003; shown in prior work to span Crus I/II/VIIb, potentially overlapping with LangCereb3; Guell et al., 2018a; King et al., 2019; Nettekoven et al., 2024; Saadon-Grossman et al., 2024; Van Overwalle, 2024), the ‘Multiple Demand’ (MD) network already mentioned above (Duncan et al., 2010; previously reported in lateral lobule VI/Crus I, potentially overlapping with LangCereb1 and LangCereb4; Desmond & Fiez, 1998; Stoodley et al., 2012; Marek et al., 2018; Guell et al., 2018a; Nettekoven et al., 2024; Saadon-Grossman et al., 2024), and the brain network that supports speech articulation (Wolna et al., 2023), previously reported in the anterior lobe, lobule VI, and lobule VIII, potentially overlapping with LangCereb4; Frings et al., 2006; Stoodley et al., 2012; Basilakos et al., 2018; Guell et al., 2018a; Correia et al., 2020). In these analyses we are testing whether the ‘peaks’ in the activation landscape for language vs. e.g., ToM are dissociable, as would be indicated by a) greater within-contrast than between-contrast topographic similarity, and b) statistically distinguishable response profiles. Not all brain regions may have sharp boundaries, but dissociations even in the presence of some overlap are still informative with respect to the functional architecture. Indeed, in LangCereb1-4, we find that the peak of language activity in all regions can be dissociated from the ToM, MD, and articulation peaks (**Suppl. Tables 25-27, Suppl. Figs. 7-9**; see <u>Discussion)</u>. Importantly, this dissociability is in no way guaranteed. Lobule IX provides a nice counterexample—the functional profile is indistinguishable between the language-defined and the ToM-defined voxel (Suppl. Fig. 7f), suggesting that the most language-responsive voxels in this area are also the most ToM-responsive.

### 3) The language-selective cerebellar region (LangCereb3) processes sentence-level meanings, shows sensitivity to language processing difficulty, and responds to both social and non-social language

Having established that one of the cerebellar language regions (LangCereb3, which spans Crus I/II/VIIb) shows selectivity for language processing, albeit less strongly than the neocortical language network, we asked: what linguistic computations does this region support? We chose to focus on this language-selective region here because its function plausibly relates to specifically linguistic processes. We still report the responses in the non-selective regions (Suppl. Fig. 11; **Suppl. Tables 29-30**, **32**; see Suppl. Fig. 2d for responses in right lobule IX), but leave characterization of their contributions to language and cognition as an exciting avenue of future work (see <u>Discussion</u>).

One paradigm that has been widely employed to characterize the linguistic contributions of the neocortical language regions uses sentences and various linguistically degraded conditions (Expt. 3a, n=71, Fig. 4a, data from Fedorenko et al., 2010; Shain, Kean et al., 2024; and Kauf et al., 2024). This paradigm tests for sensitivity to core components of language processing: lexical access (recognizing individual words), syntactic-structure building (determining how words in a sentence are structurally related to, or dependent on, one another), and semantic composition (constructing a sentence-level representation given the individual words and the syntactic dependency structure; Fedorenko et al., 2010; Pallier et al., 2011; Giglio et al., 2022; Shain, Kean et al., 2024). The paradigm included meaningful grammatical sentences (S; which require all three components), lists of unconnected words (W; which only require accessing stored lexical items), grammatical but meaningless Jabberwocky sentences (J; which only require constructing a syntactic dependency structure), and nonword lists (N; which do not require lexical access, nor syntactic-structure building, nor semantic composition; Fig. 4a; see Suppl. Fig. 10 for responses to each of Expt. 3a’s sub-experiments, which shared the same design but used different materials and participants).

**Figure 4:**
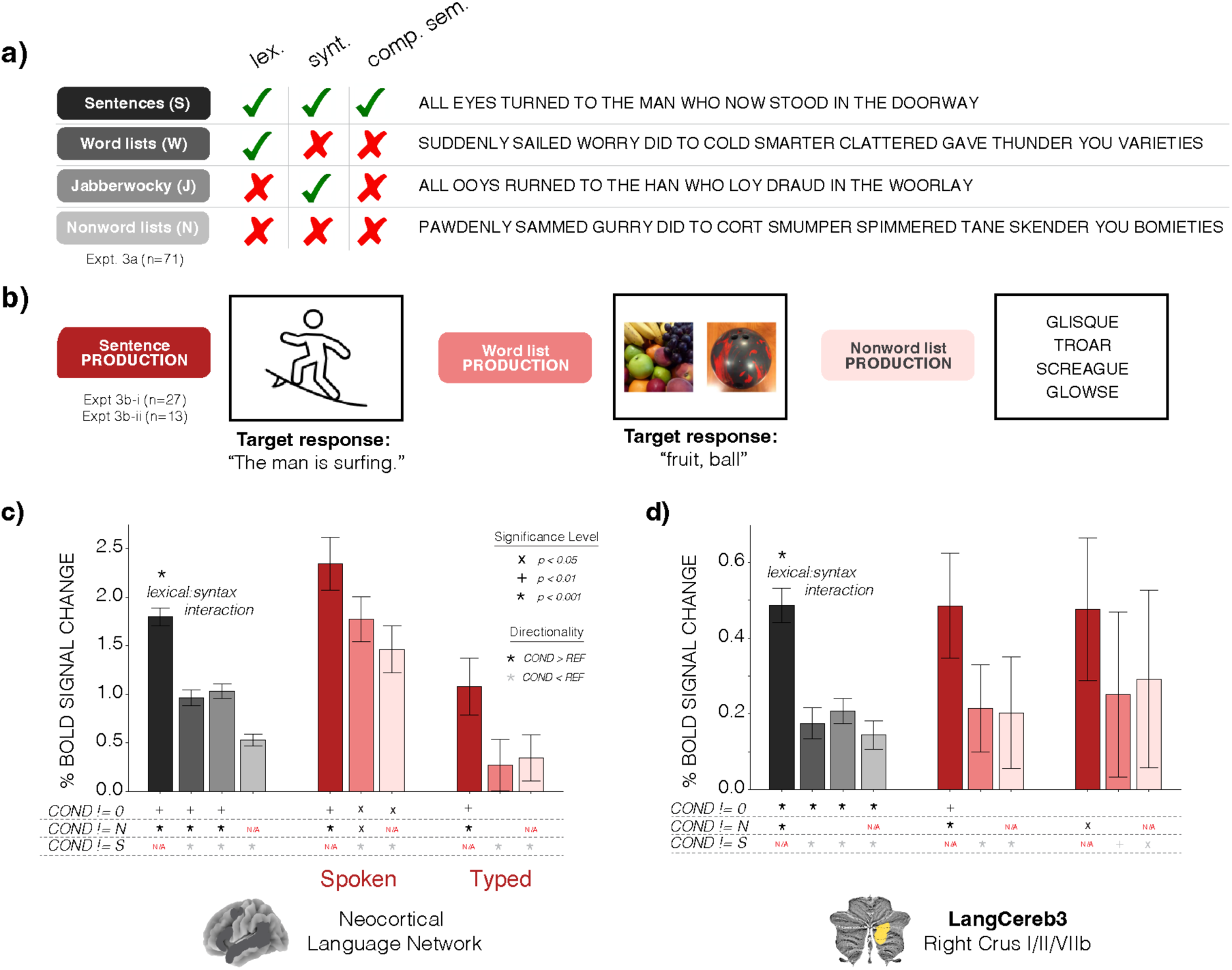
Linguistic contributions of LangCereb3 (Crus I/II/VIIb) during comprehension and production. a) Language comprehension experiment (Expt. 3a, n=71, data from Fedorenko et al., 2011; Shain, Kean et al., 2024; and Kauf et al., 2024). Participants read sentences and various linguistically degraded conditions that were designed to isolate aspects of language processing. Sentences and word lists both required accessing real lexical items (Jabberwocky sentences and nonword lists did not), whereas sentences and Jabberwocky both required building a syntactic structure (word and nonword lists did not); finally, only sentences allowed for semantic composition (see <u>Methods</u> for additional experimental details). b) Language production experiment (Expt. 3b, n=40, data from Hu, Small et al., 2023). The design mirrored the language comprehension experiment—participants were asked to produce a sentence given an image of an event, a list of words given multiple images of objects, or recite a list of nonwords (<u>Methods</u>). **c-d)** Responses to language comprehension and production experiments in the neocortical language network (averaged over the five ‘core’ left-hemisphere language regions (<u>Methods)</u>, (**c**) and LangCereb3 (Crus I/II/VIIb) (**d**). For the comprehension experiment, fROIs were identified and responses were measured using Independent runs of the data. Responses reflect the percent BOLD signal change in individually-defined language fROIs (<u>Methods</u>). All error bars reflect the standard error of the mean over participants for the given condition. Condition responses were evaluated relative to i) a fixation baseline, ii) the response to nonword lists (only S, W, and J responses, in the relevant experiment), and iii) the response to sentences (only W, J and N responses, in the relevant experiment) using linear mixed-effects models (see **Suppl. Tables 28**, **31**, <u>Methods</u>).

All neocortical language regions show the strongest response to sentences and the weakest response to nonword lists, with the response to word lists and Jabberwocky sentences falling in between (significantly below the response to sentences, and significantly above the nonword response; **Suppl. Table 28**; Fig. 4c). This pattern suggests that the neocortical language network supports computations related to lexical access and syntactic-structure building. Furthermore, lexical access and syntactic-structure building interact such that the response to sentences is greater than the sum of the response to word lists and Jabberwocky sentences (p<0.001, **Suppl. Table 33**). This result indicates that the neocortical language network additionally supports computations related to semantic composition (Fedorenko et al., 2010, see Shain, Kean et al., 2024 for more discussion; Fig. 4a, c, **Suppl. Table 28**). In contrast, LangCereb3 (Crus I/II/VIIb) exhibited a strong response to sentences, but a low response to all three linguistically degraded conditions (i.e., responses to word lists and Jabberwocky sentences were similar in magnitude to the level of response to the nonword lists; ps>0.05; **Suppl. Table 31**, Fig. 4d). This pattern suggests that LangCereb3 primarily supports compositional semantic processing, and, in fact, LangCereb3 was slightly more sensitive to compositional semantic processing than the neocortical language network, as evidenced by a three-way interaction (p<0.05, **Suppl. Table 34**, <u>Methods</u>, see <u>Discussion</u>).

We next investigated whether LangCereb3 also supports compositional semantic processing during language production. Participants had to produce, by speaking out loud, sentences (by describing pictures of events; the ‘Sentence’ condition), words (by naming pictures of objects; the ‘Word list’ condition), and nonwords (by reciting lists of monosyllabic nonwords; e.g., “sloal”; the ‘Nonword list’ condition; Expt. 3b-i, n=27, data from Hu, Small et al., 2023; Fig. 4b). A separate cohort of participants completed a typed version of this paradigm (Expt. 3b-ii, n=13, data from Hu, Small et al., 2023). Mirroring the pattern in comprehension, the neocortical language regions exhibit the strongest response during sentence production, an intermediate response to word-list production, and the lowest response to nonword-list production (**Suppl. Table 28**, Fig. 4c; Hu, Small et al., 2023). In contrast, LangCereb3 (Crus I/II/VIIb) responded strongly during sentence production (**Suppl. Table 31**) and showed a similarly low response to word-list and nonword-list production, both during speaking and typing (ps>0.05; **Suppl. Table 31**, Fig. 4d), in line with its comprehension profile.

The three non-selective cerebellar language regions exhibited diverse functional profiles during both comprehension and production, with none showing a profile that perfectly mirrored the neocortical language network (Suppl. Fig. 11; **Suppl. Tables 29-30**, **32**; see Suppl. Fig. 2d for responses in right lobule IX).

To perform a more fine-grained evaluation of how responses in LangCereb3 (Crus I/II/VIIb) are modulated by linguistic features, we examined its responses to 1,000 diverse sentences (Expt. 3c, n=5, data from Tuckute et al., 2024; Fig. 5a). First, we examined the overall similarity in the response profile between LangCereb3 and the neocortical language network. The responses (averaged across the 5 participants) were strongly correlated, such that if a sentence elicited a strong response in the neocortical network, it also elicited a strong response in LangCereb3 (r=0.638, Fig. 5b). In fact, the responses *between* LangCereb3 and the neocortical language network were as similar across participants as the responses *within* either of the regions (p>0.05, one-sample Wilcoxon signed-rank test, Fig. 5c, <u>Methods</u>).

**Figure 5:**
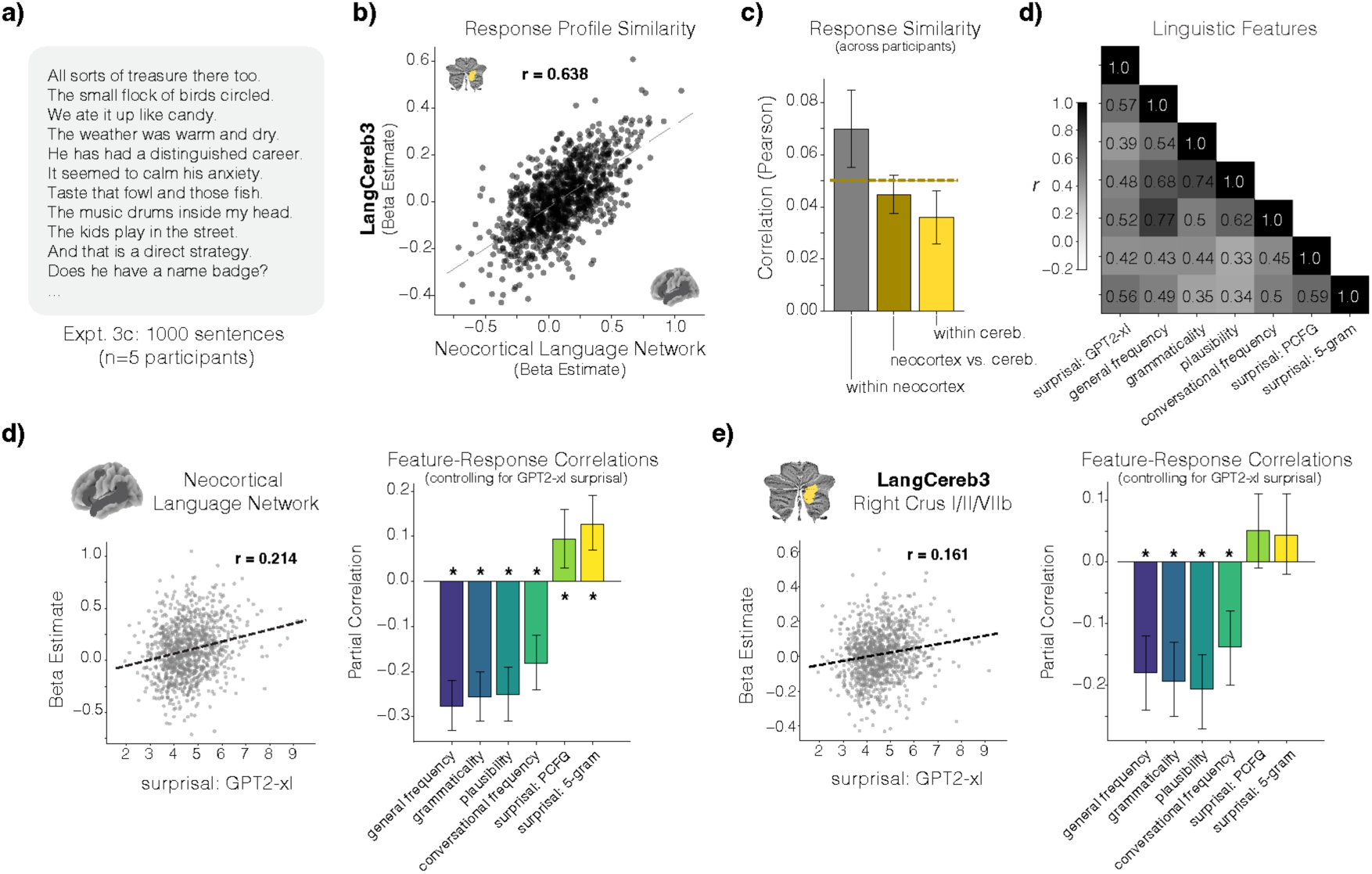
Responses in LangCereb3 (Crus I/II/VIIb) and the neocortical language network are modulated by processing difficulty. **a)** A deeply-sampled dataset of 5 participants who read 1,000 sentences each (Expt. 3c, n=5, data from Tuckute et al., 2024). **b)** Similarity of responses to the sentences in the neocortical language network and LangCereb3 (averaged across the 5 participants). **c)** Response similarity across participants within vs. between regions (neocortical language network and LangCereb3). Similarity between regions was assessed with the same amount of data as the within-region comparison. Error bars reflect standard error of the mean over all possible participant pairs. The dashed line reflects the geometric mean of the within-region correlations. If the responses in the two regions differed, we would expect the between region correlations to be significantly lower than the geometric mean of the within region correlations. We did not find a significant difference (p>0.05, one-sample Wilcoxon signed-rank test). **d)** The sentences were annotated using a variety of linguistic features that captured how difficult a sentence is to process (e.g., surprisal, grammaticality, frequency; <u>Methods</u>). The correlation of these features over the entire set of 1,000 sentences is shown. **e-f)** Modulation of brain responses in the neocortical language network (**e**) and LangCereb3 (Crus I/II/VIIb, (**f**) by individual features. Feature-response correlations reflect the partial correlation of responses in a region with a given feature, controlling for GPT2-xl surprisal. The order of the features was defined based on the magnitude of the partial correlation in the neocortical language network. In (**f**), the individual features are presented in the same order as in (**e**) to highlight the similarity of the features that modulate responses in the neocortical language network and LangCereb3. Error bars reflect the 95% parametric confidence interval (**Suppl. Tables 35-39**).

We then considered how responses in LangCereb3 were modulated by sentence-level features that relate to processing difficulty—a factor known to modulate responses in the neocortical network (Just et al., 1996; Ben-Shachar et al., 2003; Blank et al., 2016; Shain, Blank et al., 2020; Shain et al., 2022; Heilbron et al., 2022). Some of these features were automatically extracted from language models and others estimated from behavioral ratings in independent participants (Fig. 5d, <u>Methods</u>; see Tuckute et al., 2024 for details). As expected—given the strong correlation in the response profile of LangCereb3 and the neocortical language network—the responses in LangCereb3 were modulated by all these features. This includes an overall measure of how predictable a sentence is (surprisal: Levy, 2008; Shannon & Weaver, 1949), estimated using GPT2-xl (Radford et al., 2019; this measure also encompasses aspects of words, grammatical structure, and the overall sentence meaning), with less predictable sentences eliciting a stronger response (Fig. 5e**-f**, <u>Methods</u>). More interestingly, after controlling for surprisal, the pattern of sensitivity to other features that affect processing difficulty was remarkably similar between LangCereb3 and the neocortical language network (Fig. 5e**-f**, **Suppl. Table 35**, **38**). Similar features also modulated responses in the three non-selective cerebellar language regions (**Suppl. Tables 36-37**, **39**), although their patterns of response were qualitatively less similar to the neocortical language network. These results indicate that LangCereb3 is not only coarsely responsive to language stimuli, but shows fine-grained sensitivity to linguistic features known to modulate neocortical language responses.

Finally, we tested whether selective linguistic responses in LangCereb3 could be explained by the frequent presence of social content in language stimuli (e.g., in the items in our language localizer, Expt. 1a, Fig. 6a). Language is a function deeply embedded in social interaction (Seyfarth & Cheney, 2018): infants acquire language through social exchanges, and much of language use is socially-grounded. In particular, in face-to-face interaction, we have to integrate diverse non-verbal social-communicative signals, such as facial expressions, intonation, and gestures, with linguistic content (Vigliocco et al., 2014; Hadley et al., 2022; Hagoort et al., 2024), and these non-verbal cues are often critical to interpretation. In many cases, interpreting others’ utterances requires social reasoning (consideration of their beliefs, desires, and emotions; Grice, 1975; Levison, 1983; Clark, 1996; Van Overwalle, 2024). Finally, although we can use language to talk about diverse topics, including highly abstract ideas that have nothing to do with the social world (e.g., math or physics), we often use language to share social information about others, and some have even hypothesized that the need to do so gave rise to the emergence of language in the first place (Dunbar, 1998). All this said, past work has suggested that in the neocortex, language processing is supported by distinct mechanisms than those supporting perception of social signals, such as facial expressions or gestures (Deen et al., 2015; Pritchett et al., 2018; Jouravlev et al., 2019), as well as the social reasoning system (Paunov et al., 2019; Shain, Chen, Paunov et al., 2022). Does this separation hold in the cerebellum?

**Figure 6:**
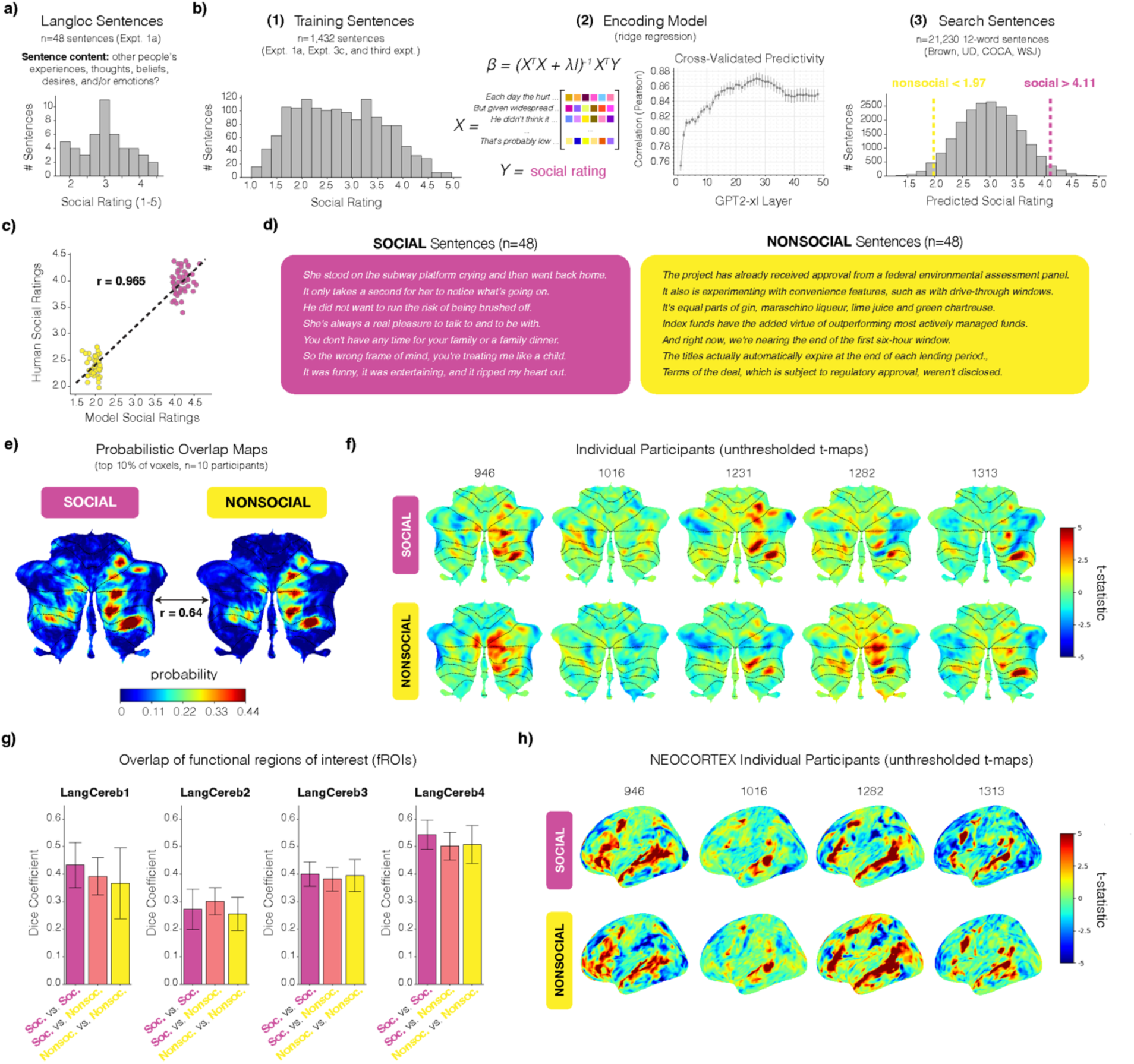
Cerebellar regions recruited by social vs. nonsocial sentences. **a)** Behavioral ratings from online participants for the n=48 sentences from the language localizer (Expt. 1a). Participants were asked to rate each sentence for how much it made them think of other people’s experiences, thoughts, beliefs, desires, and/or emotions on a scale from 1 (not at all) to 5 (very much). **b)** The procedure for sentence selection consisted of three steps. First, behavioral ratings were collected from online participants for 1,432 sentences. Second, an encoding model (ridge regression) was trained to predict the average per-sentence ratings from sentence-level embeddings from GPT2-xl. The encoding model was evaluated by computing—for each layer separately—the Pearson correlation between predicted and actual ratings on the left-out test set (and then averaged over the 5 cross-validation folds; <u>Methods</u>). The best-performing layer (#27) achieved the predicted-to-actual correlation of r=0.87. Third, the encoding model (trained using embeddings from the best-performing model layer) was used to predict the socialness ratings for 21,230 sentences extracted from publicly-available corpora. These predicted ratings were used to identify sentences that were highly social or nonsocial. **c)** The predicted socialness ratings were validated against human ratings collected for these 96 sentences. A correlation of 0.965 was observed, and the two sets of sentences were fully non-overlapping in the human ratings. **d)** Example social and nonsocial sentences from the final set of 96 sentences (<u>Methods</u>). **e)** Probabilistic overlap maps across n=10 participants constructed from individual binarized Language>Control maps. Individual maps have a value of 1 for the top 10% of language-responsive voxels across the whole brain and 0 otherwise. Color shading of the overlap map reflects the proportion of participants for whom that voxel was in the top 10% of their most language-responsive voxels. **f)** Cerebellar activations in 5 individual participants for the Social>Control vs. Nonsocial>Control contrasts. The maps are unthresholded t-statistic maps; cerebellar lobule boundaries are marked in black. **g)** Overlap of the functional regions of interest (fROIs) that are identified depending on the sentence type (social vs. nonsocial) that is used in the language localizer. (The between-condition Dice coefficients were computed with the same amount of data as the within-condition coefficients.) **h)** Neocortical activations in 4 individual participants for the Social>Control vs. Nonsocial>Control contrasts. The maps are unthresholded t-statistic maps.

In Section 2 we already showed that responses in LangCereb3 to dynamic faces and during mentalizing were much lower than during language processing, and here we examine the effect of the social content conveyed through language on responses in LangCereb3. The sensitivity to linguistic processing difficulty, just discussed, suggests that the responses are modulated by content-independent features related to the structure and overall plausibility of the sentence. Nevertheless, many studies have reported responses to social stimuli in the Crus area, close to LangCereb3 (e.g., Guell et al., 2018a; King et al., 2019; Nettekoven et al., 2024; Saadon-Grossman et al., 2024; Van Overwalle, 2024), and it is possible that processing difficulty features are correlated with particular kinds of semantic content. To this end, we designed two new versions of the language localizer: one with highly social sentences (e.g., *She is always a real pleasure to talk to and to be with.*) and another with highly nonsocial sentences (e.g., *The titles actually automatically expire at the end of each lending period.*), matched for surprisal (average sentence-level surprisal 3.86 vs. 3.84; p>0.05, Mann-Whitney U test). The sentences were selected from language corpora using an encoding model trained to predict human socialness ratings from language model embeddings (Fig. 6b**-c**, <u>Methods</u>). Nonword sequences were used as the control condition, similar to Experiment 1. We then tested whether these two localizer versions engage the same regions in the cerebellum, including critically within LangCereb3. We observed strong similarity in the pattern of activation for the two contrasts (Social sentences > Control; Nonsocial sentences > Control) at the group level (r=0.64, Fig. 6e), and in individual participants (Fig. 6f). The overlap between these contrasts within LangCereb3, as quantified by the Dice coefficient, was similar to the Dice coefficient based on the comparison across runs within the same type of stimulus-content (Fig. 6g). These results suggest that the same cerebellar regions are recruited by social *and* nonsocial sentences, similar to the neocortical language network (Fig. 6h).

### 4) All cerebellar language regions—but especially LangCereb3—bear functional similarity to the neocortical language network

In Sections 2 and 3 we found that, relative to the other cerebellar language regions, the response profile of LangCereb3 (Crus I/II/VIIb) was more qualitatively similar to the neocortical language network. Here, we quantify the relationship between the right cerebellar language regions and the left-hemisphere neocortical language network, considering both the experimental response profiles and patterns of functional connectivity during naturalistic cognition.

In line with the analyses reported in Sections 2 and 3, LangCereb3 (Crus I/II/VIIb) showed the greatest similarity in its experimental response profile (i.e., the pattern of univariate response magnitudes across Expts. 2a-e and 3a-b) to the neocortical language network (r=0.81, p<0.001, Fig. 7c). The experimental response profiles of the other cerebellar language regions were positively but less strongly correlated with the neocortical network (LangCereb1: r=0.56, Fig. 7a; LangCereb2: r=0.49, Fig. 7b; and LangCereb4: r=0.71, Fig. 7d; ps<0.001; see Suppl. Fig. 2e**-f** for response profile correlations from right lobule IX and Suppl. Fig. 12a**-i** for response profile correlations from the left cerebellar homotopes). In line with the results in Section 2, each of these regions exhibited clear ‘off-diagonal’ conditions that elicited a stronger response than in the neocortical language regions (e.g., LangCereb2 responded relatively more strongly to the visual semantically-rich conditions: Expts. 2e-i and 2e-ii, blue data points, Fig. 7b).

**Figure 7:**
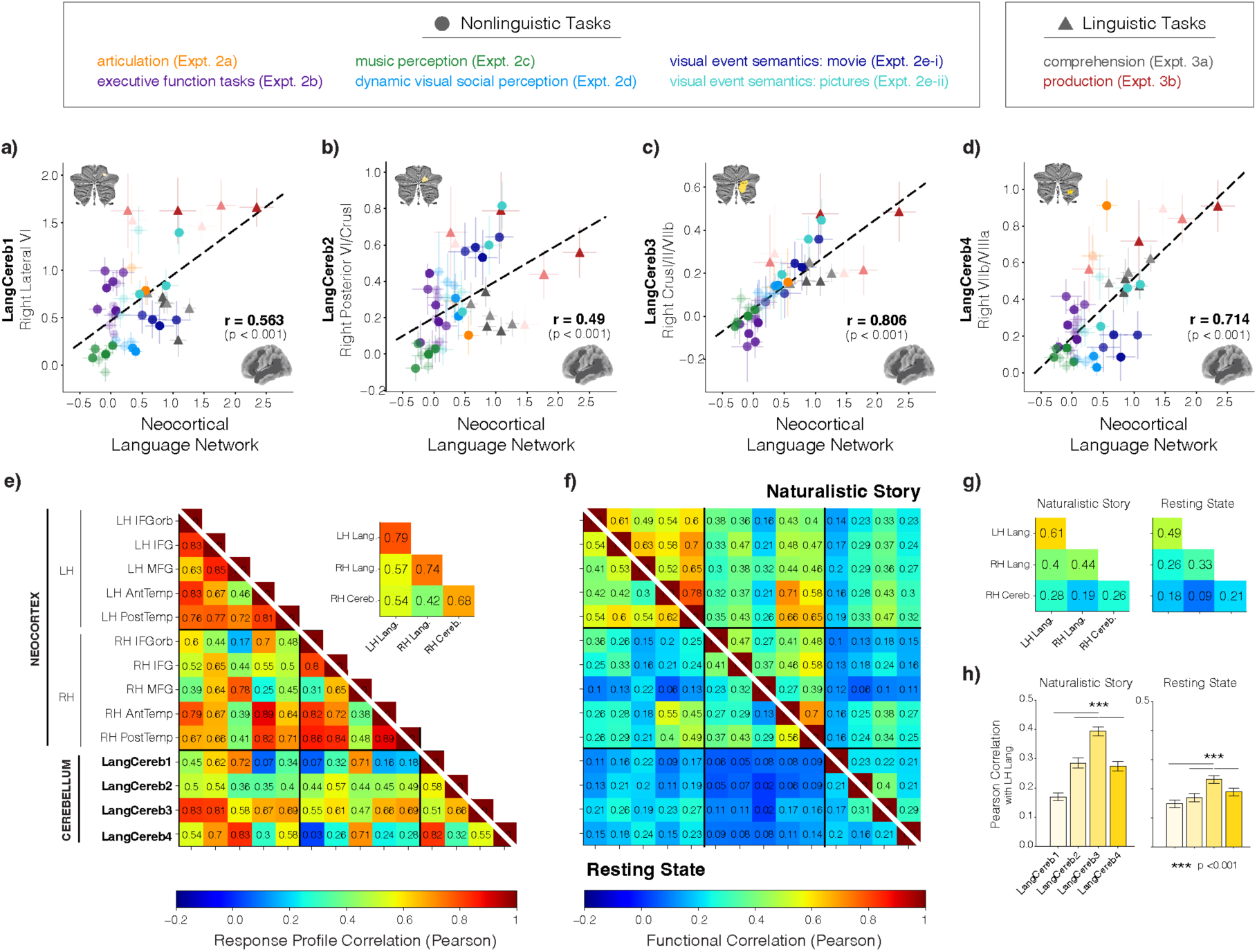
Response profile similarity and functional connectivity of the cerebellar language regions with the neocortical language network. **a-d)** Correlation in the response profiles (i.e., the magnitudes of response from Expts. 2a-e and 3a-b, Fig. 3 and 4) of the cerebellar language regions and the neocortical language network (averaged over the five ‘core’ regions, <u>Methods</u>). Responses to individual conditions are shown in the same colors as in Fig. 3 and 4. Error bars around individual points reflect the standard error of the mean over participants for that condition in the neocortical language regions (x-error) and a given cerebellar language region (y-error). Responses during sentence and nonword-list comprehension (e.g., Expt. 3a) are not included in the scatter plots because the voxels from which responses were measured were selected to have a large difference between these conditions and could, therefore, inflate correlation values. Black dashed lines correspond to the line of best fit (<u>Methods</u>). **e)** Correlation in the response profiles between the cerebellar language regions and individual neocortical regions, including right-hemisphere neocortical language regions. *Inset:* Summary of response profile correlations by gross anatomical region (left-hemisphere neocortex, right-hemisphere neocortex, and right cerebellum). **f)** Functional correlations between the cerebellar and neocortical language regions during resting state (lower triangle) or naturalistic story listening (upper triangle; Expt. 4a-b, n=85, data from Malik-Moraleda, Ayyash et al., 2022; <u>Methods</u>). **g)** Summary of functional correlations by gross anatomical region. **h)** Functional correlations between individual cerebellar language regions and the left-hemisphere neocortical language network.

It is also worth noting that the response profiles of the right cerebellar language regions were all relatively more correlated with the contralateral (left), compared to the ipsilateral (right), neocortical language regions (Fig. 7e, **Suppl. Fig 13a-d**, except LangCereb2). Additionally, within the cerebellum, LangCereb3 showed the least similar response profile to its left-hemisphere cerebellar homotopic region (r=0.51, Suppl. Fig. 13g), in contrast to the other cerebellar regions whose profiles were more symmetrical (LangCereb1: r=0.88, Suppl. Fig. 13e; LangCereb2: r=0.73, Suppl. Fig. 13f; LangCereb4: r=0.82, Suppl. Fig. 13h).

Although the experiments in Sections 2 and 3 span quite diverse conditions, they of course do not comprehensively cover the space of mental computations required in our daily lives. Therefore, in a complementary approach, we examined the degree to which the cerebellar language regions co-fluctuate with the neocortical language network during two naturalistic-cognition paradigms: resting state (Fig. 7f lower triangle) and naturalistic language comprehension (Fig. 7f upper triangle; Expt. 4a, n=85 participants listening to an excerpt from *Alice in Wonderland*; data for both conditions come from Malik-Moraleda, Ayyash et al., 2022). Across cerebellar regions, functional correlations were higher during story listening than during rest (p<0.001, **Suppl. Table 40**, Fig. 7g), in line with what has been reported for the neocortical language regions (Blank et al., 2014), and higher between the cerebellar language regions and the left-hemisphere, compared to right-hemisphere, neocortical language regions (p<0.01, **Suppl. Table 41**, Fig. 7g; see Suppl. Fig. 2g for functional correlations from right lobule IX and Suppl. Fig. 11j**-k** for functional correlations from the left cerebellar homotopes). Critically, mirroring the correlations in response magnitudes to diverse experimental conditions (Fig. 7a-d), LangCereb3 (Crus I/II/VIIb) showed the strongest functional connectivity with the neocortical language network during both rest and story listening (ps<0.0001, **Suppl. Table 40**; Fig. 7h). Therefore, in terms of both response profile similarity and patterns of functional correlations, LangCereb3 appears to be the cerebellar language region that is most functionally integrated with the left hemisphere cortical language network.

At the level of individual neocortical regions, the response profile of LangCereb3 (Crus I/II/VIIb) was numerically most similar to that of the inferior frontal language regions: in IFGorb and IFG (r=0.83 and r=0.81, respectively, Fig. 7e). However, LangCereb3 showed a statistically stronger functional correlation with the temporal rather than the frontal language regions (only during the naturalistic story condition; AntTemp: r=0.49; PostTemp: r=0.45; p<0.001; **Suppl. Table 42**, Fig. 7f). This discordance suggests that experimental responses and patterns of functional connectivity capture distinct aspects of a region’s function and highlight why functional connectivity data alone may be insufficient to fully characterize brain areas (see <u>Discussion</u>).

## Discussion

In the present study, we used precision fMRI to identify cerebellar regions that respond robustly to language across the written and auditory modalities. We identified four such regions in the right posterior cerebellum and subsequently characterized them across 26 diverse experimental and naturalistic paradigms. We found that all four cerebellar language regions bear similarity in their response profiles to the neocortical language regions and are functionally connected to them. However, the cerebellar regions differ in their degree of language selectivity and how closely their response profiles resemble those of the neocortical language regions. One cerebellar region (LangCereb3, spanning Crus I, Crus II, and lobule VIIb) is especially strongly integrated with the neocortical language network. This region responds selectively to language, is engaged by sentence-level meanings during both comprehension and production and shows sensitivity to linguistic processing difficulty. The remaining three cerebellar language regions show mixed-selective profiles, with strong responses to language but also to motor tasks (LangCereb4, spanning lobules VIIb and VIIIa), to demanding non-linguistic tasks (LangCereb1, spanning lateral lobule VI, and LangCereb4), and to meaningful visual stimuli (LangCereb2, spanning posterior lobule VI and Crus I). Below we discuss the significance of these findings and situate them in the context of prior work on the cerebellum and the neural infrastructure of language processing.

### The language network includes components in the right cerebellum

For decades, the neuroscientific community has overlooked the importance of the cerebellum in cognition and language. Even today, despite substantial neuropsychological and neuroimaging evidence to the contrary (Schmahmann & Sherman, 1998; Mariën et al., 2001; Stoodley & Schmahmann, 2010; E et al., 2014; Mariën et al., 2014, Guell et al., 2018a; 2018b; King et al., 2019; Satoer et al., 2024; Nettekoven et al., 2024; Saadon-Grossman et al., 2024), many in the field have remained skeptical of the cerebellum’s role in language beyond the motor aspects of speech articulation (LeBel & D’Mello, 2023; Wang et al., 2025). In the current study, we build on the foundation of prior studies of the linguistic cerebellum to provide a comprehensive assessment of the cerebellum’s contributions to language processing and its relationship with the neocortical language network.

We found one cerebellar language region—***LangCereb3***, spanning Crus I/II/VIIb—that selectively responds to linguistic input, showing little or no above-baseline response when participants perform demanding working memory tasks, listen to music, process socially-rich stimuli, or engage in other non-verbal tasks. This selectivity demonstrates that this region does not support motor demands, general cognitive demands, or general processing of hierarchical structure, social/communicative signals, or meaningful stimuli. Although future work may discover that this region is engaged as strongly in some non-linguistic tasks as it is during language processing, the evidence so far suggests that this region supports specifically linguistic computations, as discussed in more detail below (‘LangCereb3 is a cerebellar satellite of the neocortical language network’). The location of this region is also the most consistent—though not entirely overlapping—with where language responses have been reported in prior studies (e.g., Stoodley & Schmahmann, 2009a; Moberget et al., 2014; Lesage et al., 2017; D’Mello et al., 2017; Guell et al., 2018; King et al., 2019; Nettekoven et al., 2024).

In addition to the language-selective region, we identified three cerebellar regions that respond to language across modalities and types of semantic content, but also to at least one non-linguistic condition. ***LangCereb1*** (lateral lobule VI) responds strongly during cognitively demanding tasks, with stronger responses to more difficult conditions (Expt. 2b, 2e). In this way, this region combines functional signatures of the neocortical language and Multiple Demand (MD) networks (Duncan, 2010; Duncan et al., 2020)—a profile that has not been reported in the neocortex (but see Wolna et al., 2025 for a similar profile in the thalamus), where the two networks are robustly distinct (see Fedorenko & Blank, 2020 for a review). This region appears to additionally play a role in motor planning and/or execution across effectors (mouth and fingers, Expt. 2a). ***LangCereb2*** (posterior VI/Crus I) responds strongly to meaningful visual stimuli, including pictures of events (Expt. 2e-ii), and naturalistic videos (Expt. 2d, 2e-i). This region therefore appears to support semantic processing across verbal and pictorial modalities, similar to some neocortical regions (outside of the canonical left-hemisphere language network; Patterson et al., 2007; Lambon Ralph et al., 2010; Huth et al., 2016; Popham et al., 2021; Ivanova, 2022). However, this region also shows some response to demanding executive tasks, with a stronger response to the more demanding condition than to the less demanding condition, although this MD-network-like profile is not as pronounced as in LangCereb1 (or LangCereb4 discussed next). Finally, ***LangCereb4*** (VIIb/VIIIa) responds strongly to motor tasks, with a numerically stronger response during speech articulation compared to the hand-motor condition (Expt. 2a), and—similarly to LangCereb1—it responds during cognitively demanding tasks, with stronger responses to more difficult conditions (Expt. 2b, 2e). This region thus shows a mix of motor/articulatory control and an MD-network-like profile. The mixed selectivities of LangCereb1, 2, and 4 stand in stark contrast to the highly selective neocortical language regions (the ‘core’ language network; Fedorenko et al., 2024) and highlight a key divergence between the cerebellar and neocortical language circuits that merits additional investigation. It is also worth noting that the functional diversity of the cerebellar language-responsive regions is at odds with the idea that language representations in the cerebellum are redundant (Guell et al., 2018a).

### The mixed-selective cerebellar language regions may integrate information from multiple neocortical networks

What could the mixed response profiles observed in LangCereb1, 2, and 4 reflect? One possibility, inspired by the findings that the cerebellum integrates sensory and motor information (Wolpert et al., 1998; Wiestler et al., 2011), is that these regions integrate linguistic and non-linguistic information. In particular, the fact that these regions respond to language and a non-linguistic condition suggests that linguistic and non-linguistic representations are superimposed in these regions. This superimposition may allow these regions to integrate information across domains (see King et al., 2023 for concordant evidence that cerebellar responses are best predicted by multiple neocortical regions rather than a single region). This possibility is especially exciting because, despite the importance of cross-domain integration to many complex tasks, from solving verbal math problems to making social inferences during conversations, few proposals exist for where or how information from distinct neocortical systems is combined.

However, at present, the data do not allow us to rule out alternative explanations of mixed selectivity. For example, the mixed selectivity may result from nearby distinct neural populations, each supporting its own computation, that are not differentiated at the spatial (or temporal) scale of fMRI. This concern may be particularly relevant for cerebellar imaging studies given the much higher neural density in the cerebellum compared to the neocortex (Azevedo et al., 2009). Higher-field fMRI (e.g., 7T) or voxel decomposition approaches (e.g., Lee & Seung, 1999; Khosla et al. 2022) may help determine whether, in reality, multiple specialized neural populations comprise these regions. This said, it is worth noting that we observe language selectivity in the largest of the cerebellar language regions—suggesting that it is possible to observe language-selective responses in the cerebellum even at the resolution of our current approach. We can also dissociate the mixed-selective cerebellar language regions from nearby regions that more closely resemble, for example, the neocortical Multiple Demand regions and articulation regions. Namely, in the vicinity of both LangCereb1 and LangCereb4, we find neural populations that show a more canonical MD-network profile. Similarly, in the vicinity of LangCereb4, we additionally find a neural population that shows a more canonical articulation-region profile. These dissociations align with prior work showing that functional networks in the cerebellum are highly interdigitated but separable (Nettekoven et al., 2024; Saadon-Grossman et al., 2024) and provide additional support for the presence of an integrative circuit in regions with mixed-selective profiles.

Yet another possibility we cannot rule out is that a region that responds to both language and a non-linguistic condition implements a (single) computation required for both conditions. Based on the current understanding of perception, cognition, and motor control, the mixed-selectivity profiles discussed above do not readily suggest clear hypotheses for such shared underlying computations (e.g., shared among language comprehension, demanding executive tasks, and motor control for LangCereb1; see Diedrichsen et al., 2019 for related discussion). But this is not to say that such hypotheses could not be developed in the future.

### LangCereb3 is a cerebellar satellite of the neocortical language network

In discussing the contributions of the linguistic cerebellum to language processing, we have chosen to focus on the region that is selective for language: LangCereb3, spanning Crus I/II/VIIb. This decision was motivated by two considerations. First, the functional selectivity of a brain region significantly constrains the space of possibilities for the region’s computations. In other words, LangCereb3’s function can be explained in terms of linguistic processes alone. The lack of similar constraints on the computations of the non-selective cerebellar language regions makes it challenging to discern their function without additional experimentation. And second, the language selectivity of both LangCereb3 and the neocortical language network suggests that they may belong to the same *functional system*—a set of brain regions that are structurally connected, share functional properties, and work together in the service of a common goal; Yeo, Krienen et al., 2011; Buckner et al., 2011; Sporns, 2014; Fedorenko & Thompson-Schill, 2014; Medaglia et al., 2015). In contrast, the mixed selectivity of the other three cerebellar language regions suggests that they may work with multiple neocortical networks, as discussed above. Therefore, LangCereb3 presented the clearest target for a systematic comparative investigation of cerebellar vs. neocortical linguistic contributions.

In a widely used paradigm that tests for sensitivity to different cognitive components of language (Fedorenko et al., 2010), we found that LangCereb3 is primarily engaged in processing sentence-level *meanings*. However, we cannot rule out this region’s contributions to lexical access and syntactic structure building: although we do not find reliable sensitivity to lexical access and syntactic structure building in LangCereb3, we don’t find evidence that LangCereb3’s contributions to these components differ statistically from those of the neocortical network, which is robustly sensitive to both processes (Shain, Kean et al., 2024). As a result, we leave the question of whether and how LangCereb3 may contribute to word retrieval and syntactic processing to future work, with finer-grained manipulations. Although it may be tempting to take sensitivity to sentence ‘grammaticality’ in Expt. 3c as evidence for LangCere3b’s role in syntactic processing, the current data do not allow for this interpretation because in natural language stimuli (like those used in Expt. 3c), grammatical well-formedness strongly affects the ability to construct a sentence-level meaning (correlation between grammaticality and plausibility judgments in Expt. 3c: r=0.74).

We do have strong evidence that LangCereb3 is engaged by sentence-level meanings during both comprehension and production (see also Fabbro et al., 2000; Cook et al., 2004; Murdoch & Whelan, 2007; Stoodley & Schmahmann, 2009b; Guell et al., 2015; D’Mello et al., 2020b; Gatti et al., 2020a; 2020b; LeBel et al., 2021; Nakatani et al., 2023; Satoer et al., 2024 for evidence that the cerebellum plays a role in semantic processing). The fact that LangCereb3 is highly similar to the neocortical language network in i) its overall functional response profile to diverse experimental manipulations, ii) its fine-grained responses to individual sentences, iii) its sensitivity to linguistic processing difficulty, and iv) its lack of sensitivity to content type (e.g., social vs. nonsocial language), together with v) strong functional connectivity with the neocortical language areas, suggests that LangCereb3—more so than the other cerebellar language regions—works closely in concert with the neocortical language network. It may aid in semantic composition or iteratively refine the constructed meanings (similar to how the cerebellum refines, rather than generates, motor movements; Manto et al., 2012). Evidence from patients with brain damage suggests that in mature brains, the cerebellum plays a secondary role in linguistic semantic processing given that cerebellar damage in adults does not cause severe aphasia, but is instead associated with more subtle linguistic deficits (e.g., Satoer et al., 2024, see ‘Clinical implications’ below).

Future work should attempt to search for differences in the role of the neocortical language network and LangCereb3 in processing sentence-level meanings, including with approaches optimized for differentiating the cerebellar and neocortical contributions (e.g., Shahshahani et al., 2024), with temporally resolved methods, and by examining sensitivity to linguistic features and manipulations beyond those examined here.

### Complementary approaches to probing the functional architecture of the cerebellum (or the brain in general)

The question of which functions are distinct in the mind and brain is among the biggest and most controversial in cognitive neuroscience. Two approaches are commonly used to tackle these questions. The approach we adopted here—functional *localization* (Saxe et al., 2006)—targets a particular mental function (in our case, language processing), identifies a subset of the brain that supports this function, and subsequently characterizes these regions using diverse manipulations with the goal of understanding their computations. This approach has a long history and many success stories (see Kanwisher, 2010 for a review). An alternative approach that emerged more recently in fMRI research—functional *parcellation*—has also proven highly successful in uncovering the brain’s organization. In contrast to functional localization, parcellation approaches do not focus on a particular functional domain, and instead ask which brain areas are functionally dissociable, often in a more bottom-up, data-driven fashion. In this approach, parcellation algorithms assign individual voxels to a set/network based on their functional response properties or patterns of functional connectivity. Thus, both of these approaches have increased our understanding of the architecture of the mind and brain, yet their goals are distinct (albeit complementary): one targets a particular mental process of interest, and the other asks about the overall organization of a region or the whole brain. Encouragingly, the two approaches are so far yielding a highly consistent picture in the neocortex: well-validated ‘localizer’ tasks activate neocortical areas that correspond well to the areas discovered from bottom-up parcellation approaches (e.g., Braga et al., 2020; DiNicola et al., 2020; Du et al., 2025; Shain & Fedorenko, 2025; see Salvo et al., 2021 for review). We suspect that a similar convergence of findings will be observed for the cerebellum. We also hope that the functional localization approach becomes more widely used to study the cerebellum, given that rich functional characterization is critical to deciphering a brain region’s computation(s).

### The implications of the current findings for theories of cerebellar function

The idea that ‘structure determines function’ (e.g., Golgi, 1874; Ramon y Cajal, 1911)—now neuroscientific dogma—remains pervasive in cerebellar research. In particular, given the relative uniformity of the cerebellar cytoarchitecture—i.e., the presence of the same circuit motif throughout the cerebellar cortex—it is frequently argued that the cerebellum supports *one* algorithmic transformation of the input it receives (a so-called ‘Universal Cerebellar Transform’; Schmahmann, 1996). The idea is indeed appealing in its parsimony and elegance (for discussions, see Schmahmann, 1991; 2010; Diedrichsen et al., 2019), and many studies in the cerebellar field have attempted to uncover this common function, often drawing on knowledge from circuit-level models of the cerebellum (Marr, 1969; Albus, 1971; Ito & Kano, 1982) and from the domain of motor control, where the role of the cerebellum is better understood (Shadmehr et al., 2010; Manto et al., 2012). However, the universal cerebellar transform idea has not gone unchallenged. Contra early observations of cytoarchitectural uniformity, many recent studies have reported that the cerebellum is more diverse in its cell morphology and patterns of gene expression (Fujita et al., 2020; Busch & Hansel, 2023; Sepp et al., 2024; King et al., 2025), In line with this structural diversity, evidence is now abundant that the cerebellum contains a plethora of functionally diverse areas (Saadon-Grossman et al., 2024; Nettekoven et al., 2024; see also Diedrichsen et al., 2019 for discussion). Furthermore, computational work has highlighted the capacity of the cerebellar circuit to support diverse transformations of its input (e.g., Cayco-Gajic & Silver, 2019).

Our findings provide further evidence of functional heterogeneity within the cerebellum. Only a few areas in the cerebellum show robust responses during language processing, and even among the language-responsive regions—a relatively small fraction of the right posterior cerebellum—there exists remarkable functional diversity, which is difficult to reconcile with the idea that the entire cerebellum implements a single computation. We emphasize that diverse batteries of tasks, such as the one used here, are critical for revealing these functional differences (cf. relying on resting state data alone; see Nettekoven et al., 2024 for related discussion). Even LangCereb3, the most language-selective region, is likely engaged in a broad range of linguistic processes (cf. a single computation). In particular, because LangCereb3 so closely mirrors the neocortical language network, it may support many of the same diverse computations: recognizing individual words (Fedorenko et al., 2010; Shain, Kean et al., 2024), predicting upcoming linguistic input at all levels (e.g., Shain, Blank et al., 2020; Heilbron et al., 2022), and integrating incoming words into the evolving syntactic structure in memory (e.g., Shain et al., 2022).

Finally, it is worth noting that past proposals about the universal transform have focused on broad constructs, such as prediction (Sokolov et al., 2017; Moberget & Ivry, 2016; Argyropoulos, 2016; Gatti et al., 2021b), sequencing (Leggio & Molinari, 2015, Van Overwalle, 2024), and timing (Ivry et al., 2002). At a high level, these ideas can of course be applied to language processing, but they can also be tied to almost any aspect of perception, cognition, and motor control. What would help advance the field in our view are hypotheses that make specific testable predictions for the relevant domain (e.g., language) and that ideally connect to theorizing and to empirical neocortical findings in this domain. For example, inspired by the hypothesis that the cerebellum supports predictive processing across domains, some have argued for cerebellar contributions to linguistic prediction (e.g., Lesage et al., 2012; Moberget et al., 2014; Miall et al., 2016; Lesage et al., 2017; D’Mello et al., 2017; cf. King et al. 2022). However, the fact that neocortical language areas are also strongly implicated in predictive processing (Lopopolo et al. 2017; Shain, Blank et al. 2020; Heilbron et al., 2022; Tuckute et al., 2024) makes interpreting the cerebellar findings challenging (see Sokolov et al., 2017 for related discussion). For example, are these cerebellar areas contributing in a unique way, or are they showing these effects because any language-responsive area would show sensitivity to predictability?

### Clinical implications

Our findings have three key implications for clinical research and practice. First, given clear evidence of cerebellar contributions to language, language function should be comprehensively evaluated following cerebellar damage due to stroke, head trauma, or surgical resection. At present, neuropsychological evaluations after cerebellar damage are not the standard of care (Wang et al., 2025), despite the fact that patients frequently report language difficulties, which impact academic/professional success and general quality of life. Many research studies that investigate cognitive consequences of cerebellar damage focus on verbal fluency measures (i.e., semantic and phonemic fluency, e.g., Leggio et al., 2000), which require retrieving words from memory but also place significant demands on executive function. Such tasks may therefore show impaired performance either because of linguistic or executive difficulties, making interpretation challenging. Moreover, fluency tasks miss critical aspects of linguistic processing having to do with syntactic and combinatorial semantic processing. A critical takeaway from our work is the need to more systematically evaluate language function following cerebellar damage. Such work will be critical scientifically, for understanding the causal contributions of the cerebellar language-responsive areas, and clinically, for ensuring that patients get the right speech-language therapy in cases when it’s needed.

Second, in connecting linguistic and cognitive deficits to particular regions of the cerebellum, the functional heterogeneity of the cerebellum should be taken into account. As we have highlighted many times, the cerebellum is comprised of multiple functionally distinct areas (e.g., Nettekoven et al., 2024, Saadon-Grossman et al., 2024), each functionally connected to a distinct subset of the neocortex (e.g., Middleton & Strick, 2001; Kelly & Strick, 2003; King et al., 2023). Although individual-level fMRI data from prior to damage are typically not available, we can use probabilistic atlases for different functional areas and networks to make more informed guesses about the likely areas affected instead of relying on anatomy alone (see Blank et al., 2017 for a discussion of this point with respect to neocortical language areas in the study of aphasia). Furthermore, because of the substantial inter-individual variability in the precise functional topographies, even greater than in the neocortex (e.g., Marek et al., 2018), precision medicine approaches, which take this variability into account, might be even more important in the cerebellum than in other brain regions.

And third, developing efficient treatments for aphasia, which occurs in many cases of left-hemisphere damage, remains a critical goal of language research (Fridriksson & Hills, 2021, Fama & Turkeltaub, 2014). Given that LangCereb3 is highly similar in its functional profile to the neocortical language network, this region may serve as an important additional target for clinical interventions (e.g., non-invasive neuromodulation), to complement current approaches, which focus largely on neocortical areas.

Finally, we have here focused on mature brains, and in adults, cerebellar damage rarely leads to full-blown aphasia. However, evidence has been accumulating that the cerebellum may be more critical during linguistic and cognitive development (Riva & Giorgi, 2000; Tiemeler et al., 2010; Stoodley & Stein 2013; D’Mello et al., 2015; Badura et al., 2018; Olson et al., 2022), so studying cerebellar responses to language during development and understanding how early damage affects linguistic and cognitive outcomes remains a critical goal.

## Conclusions

In conclusion, we present a large-scale analysis of the cerebellum’s role in language processing, and we establish the existence of four cerebellar language regions—one language-selective region and three non-selective regions—that we argue should be the focus of future investigations of the cerebellum’s contributions to language. We join a growing number of researchers calling for the inclusion of the cerebellum in theories of neural language processing (Mariën et al., 2001; Murdoch et al., 2010; Mariën et al., 2014; LeBel & D’Mello, 2024; Satoer et al., 2024), and we hope that a greater focus on building domain-specific knowledge will ultimately enable the vast computational and theoretical literatures associated with cerebellum-like structures across biological species (Marr, 1969; Albus, 1971; Ito & Kano, 1982; Bell et al., 2008; Babadi & Sompolinsky, 2014; Litwin-Kumar et al., 2017; Cayco-Gajic & Silver, 2019, Muscinelli et al., 2023) to be applied to studies of the cognitive cerebellum.

## Methods

### Participants

903 unique individuals (age 18-81 years, average = 25.2, st. dev = 8.1; 526 female) participated in Expts. 1-4 for payment between 09/2007 and 04/2024 across 1,103 unique scanning sessions. All participants gave written informed consent in accordance with the Massachusetts Institute of Technology’s (MIT) Committee On the Use of Humans as Experimental Subjects (COUHES). All participants were native (age of first exposure <10 years old) or highly proficient (n=197) speakers of English (for evidence that the same language network is recruited for native and highly proficient speakers see Malik-Moraleda, Ayyash et al., 2022 and Malik-Moraleda, Jouravlev et al., 2024). The majority of participants were right-handed (n=795), as assessed with the Edinburgh Handedness Inventory or by self-report (see Willems et al., 2014 for a discussion about the importance of including left-handed participants in cognitive neuroscience studies). Detailed demographic information (i.e., age, gender, native language status, and handedness) is provided by experiment in **Table 1**. All participants completed a language localizer (Expts. 1a-b) in addition to at least one critical task (Expts. 2-4).

**Table 1:**
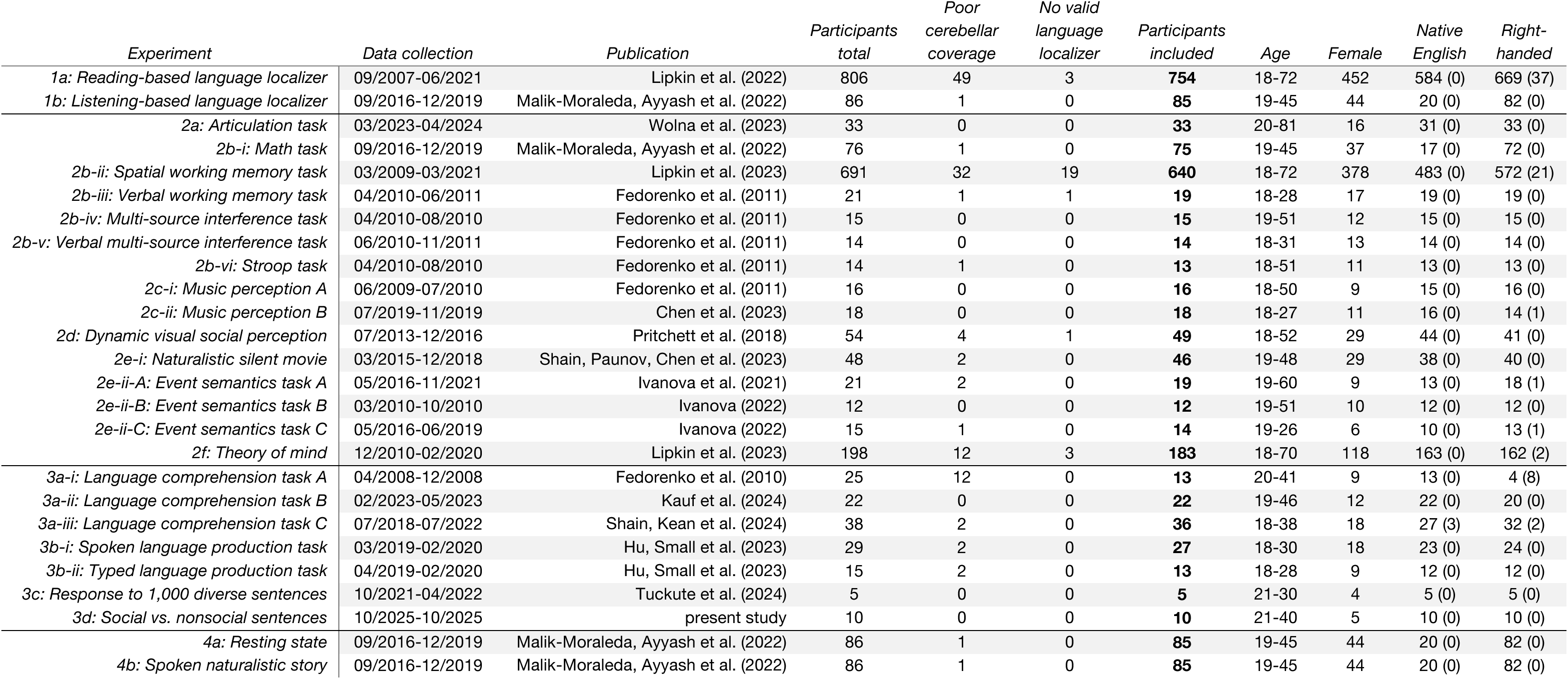
Summary of participants by experiment.

| *Experiment* | *Data collection* | *Publication* | *Participants*<br>*total* | *Poor*<br>*cerebellar*<br>*coverage* | *No valid*<br>*language*<br>*localizer* | *Participants*<br>*included* | *Age* | *Female* | *Native*<br>*English* | *Right-*<br>*handed* |
| --- | --- | --- | --- | --- | --- | --- | --- | --- | --- | --- |
| *1a: Reading-based language localizer* | 09/2007-06/2021 | Lipkin et al. (2022) | 806 | 49 | 3 | **754** | 18-72 | 452 | 584 (0) | 669 (37) |
| *1b: Listening-based language localizer* | 09/2016-12/2019 | Malik-Moraleda, Ayyash et al. (2022) | 86 | 1 | 0 | **85** | 19-45 | 44 | 20 (0) | 82 (0) |
| *2a: Articulation task* | 03/2023-04/2024 | Wolna et al. (2023) | 33 | 0 | 0 | **33** | 20-81 | 16 | 31 (0) | 33 (0) |
| *2b-i: Math task* | 09/2016-12/2019 | Malik-Moraleda, Ayyash et al. (2022) | 76 | 1 | 0 | **75** | 19-45 | 37 | 17 (0) | 72 (0) |
| *2b-ii: Spatial working memory task* | 03/2009-03/2021 | Lipkin et al. (2023) | 691 | 32 | 19 | **640** | 18-72 | 378 | 483 (0) | 572 (21) |
| *2b-iii: Verbal working memory task* | 04/2010-06/2011 | Fedorenko et al. (2011) | 21 | 1 | 1 | **19** | 18-28 | 17 | 19 (0) | 19 (0) |
| *2b-iv: Multi-source interference task* | 04/2010-08/2010 | Fedorenko et al. (2011) | 15 | 0 | 0 | **15** | 19-51 | 12 | 15 (0) | 15 (0) |
| *2b-v: Verbal multi-source interference task* | 06/2010-11/2011 | Fedorenko et al. (2011) | 14 | 0 | 0 | **14** | 18-31 | 13 | 14 (0) | 14 (0) |
| *2b-vi: Stroop task* | 04/2010-08/2010 | Fedorenko et al. (2011) | 14 | 1 | 0 | **13** | 18-51 | 11 | 13 (0) | 13 (0) |
| *2c-i: Music perception A* | 06/2009-07/2010 | Fedorenko et al. (2011) | 16 | 0 | 0 | **16** | 18-50 | 9 | 15 (0) | 16 (0) |
| *2c-ii: Music perception B* | 07/2019-11/2019 | Chen et al. (2023) | 18 | 0 | 0 | **18** | 18-27 | 11 | 16 (0) | 14 (1) |
| *2d: Dynamic visual social perception* | 07/2013-12/2016 | Pritchett et al. (2018) | 54 | 4 | 1 | **49** | 18-52 | 29 | 44 (0) | 41 (0) |
| *2e-i: Naturalistic silent movie* | 03/2015-12/2018 | Shain, Paunov, Chen et al. (2023) | 48 | 2 | 0 | **46** | 19-48 | 29 | 38 (0) | 40 (0) |
| *2e-ii-A: Event semantics task A* | 05/2016-11/2021 | Ivanova et al. (2021) | 21 | 2 | 0 | **19** | 19-60 | 9 | 13 (0) | 18 (1) |
| *2e-ii-B: Event semantics task B* | 03/2010-10/2010 | Ivanova (2022) | 12 | 0 | 0 | **12** | 19-51 | 10 | 12 (0) | 12 (0) |
| *2e-ii-C: Event semantics task C* | 05/2016-06/2019 | Ivanova (2022) | 15 | 1 | 0 | **14** | 19-26 | 6 | 10 (0) | 13 (1) |
| *2f: Theory of mind* | 12/2010-02/2020 | Lipkin et al. (2023) | 198 | 12 | 3 | **183** | 18-70 | 118 | 163 (0) | 162 (2) |
| *3a-i: Language comprehension task A* | 04/2008-12/2008 | Fedorenko et al. (2010) | 25 | 12 | 0 | **13** | 20-41 | 9 | 13 (0) | 4 (8) |
| *3a-ii: Language comprehension task B* | 02/2023-05/2023 | Kauf et al. (2024) | 22 | 0 | 0 | **22** | 19-46 | 12 | 22 (0) | 20 (0) |
| *3a-iii: Language comprehension task C* | 07/2018-07/2022 | Shain, Kean et al. (2024) | 38 | 2 | 0 | **36** | 18-38 | 18 | 27 (3) | 32 (2) |
| *3b-i: Spoken language production task* | 03/2019-02/2020 | Hu, Small et al. (2023) | 29 | 2 | 0 | **27** | 18-30 | 18 | 23 (0) | 24 (0) |
| *3b-ii: Typed language production task* | 04/2019-02/2020 | Hu, Small et al. (2023) | 15 | 2 | 0 | **13** | 18-28 | 9 | 12 (0) | 12 (0) |
| *3c: Response to 1,000 diverse sentences* | 10/2021-04/2022 | Tuckute et al. (2024) | 5 | 0 | 0 | **5** | 21-30 | 4 | 5 (0) | 5 (0) |
| *3d: Social vs. nonsocial sentences* | 10/2025-10/2025 | present study | 10 | 0 | 0 | **10** | 21-40 | 5 | 10 (0) | 10 (0) |
| *4a: Resting state* | 09/2016-12/2019 | Malik-Moraleda, Ayyash et al. (2022) | 86 | 1 | 0 | **85** | 19-45 | 44 | 20 (0) | 82 (0) |
| *4b: Spoken naturalistic story* | 09/2016-12/2019 | Malik-Moraleda, Ayyash et al. (2022) | 86 | 1 | 0 | **85** | 19-45 | 44 | 20 (0) | 82 (0) |

### Participant exclusion criteria

Across all experiments, participants who had poor cerebellar coverage during data acquisition were excluded from the analysis. Cerebellar coverage was systematically evaluated by calculating the number of voxels within a standard cerebellar anatomical mask (Diedrichsen et al., 2009) for which there was no signal in the raw file (after transforming the raw images into the MNI space). Participants were included if they had fewer than 50 voxels (of 20,943 total voxels in the mask, 0.24%) cut off during acquisition (during the critical or localizer scan). From qualitative inspection, the inferior-most portions of the cerebellum (e.g., lobule VIII) were the most likely to have been cut off during acquisition. The number of participants excluded by this criterion is detailed by experiment below, but it was typically fewer than 5 participants in a given experiment. Additionally, participants were excluded if they did not have two runs of the language localizer (needed to estimate the response magnitude in independent data). No other exclusion criteria (e.g., based on motion) were applied in order to maintain as large a sample size per experiment as possible, and in general, the data quality (including motion) was high. The exclusion criteria left 836 unique individuals (1,023 unique scanning sessions; age 18-81 years, average = 25.2, st. dev = 8.2; 495 female; n=177 proficient non-Native speakers; n=744 right-handed) across Expts. 1-4.

Summary of the data used in the current study, participant exclusions, and demographic information by experiment (Expts. 1-4). See ‘Participant exclusion criteria’ above for how poor cerebellar coverage was defined. Numbers in parentheses in the last two columns correspond to the number of participants for whom that information (i.e., native language status or handedness) was not collected.

### fMRI Data Acquisition, Preprocessing, and Analysis

#### fMRI data acquisition

Because the data were collected over the period of 16 years, several sequences were used. All the data were collected on 3 Tesla Siemens scanners at the Athinoula A. Martinos Imaging Center at the McGovern Institute for Brain Research. 926 of the 1,023 sessions were collected on a Trio scanner and the remaining 97 were collected after a Prisma upgrade. Data from most sessions were collected using one of the representative structural and functional sequences summarized in **Table 2**. Refer to OSF https://osf.io/y5t46/ for a complete description of the acquisition parameters used for each session. Although the scanning parameters differed slightly over the years of data collection, the parameters that would have had the biggest impact on the measured responses (e.g., TR, TE, voxel, field strength) were largely or entirely held constant. Most importantly, there were no systematic differences between tasks in these parameters so the minor differences across the sequences used over the years should not lead to any bias in interpretation.

**Table 2:** Summary of MRI sequence information.

| *series_description* | *matrix_size* | *slices* | *TR* | *TE* | *flip* | *IAA* | *ST* | *SDP* | *SD* | *AO* | *FR* | *FPE* | *IPRe* | *scanner* | *model* | *count* |
| --- | --- | --- | --- | --- | --- | --- | --- | --- | --- | --- | --- | --- | --- | --- | --- | --- |
| *T1_MPRAGE_1iso* | 256x256 | 176 slices | 2530.0 | 3.48 | 9.0 | 2 | 1.0 | 0.0 | 1.0 | NA | 256.0 | 256.0 | 1.000 x 1.000 | SIEMENS | TrioTim | 895 |
| *t1_mprage_sag_p2_iso_0.9mm_iso* | 288x288 | 240 slices | 1800.0 | 2.37 | 8.0 | NONE | 0.85 | 0.0 | 0.85 | NA | 250.0 | 250.0 | .868 x .788 | SIEMENS | Prisma | 96 |
| *T1_MPRAGE_4min_improved* | 256x256 | 128 slices | 2000.0 | 3.39 | 9.0 | 2 | 1.33 | 0.0 | 1.33 | NA | 256.0 | 256.0 | 1.000 x 1.333 | SIEMENS | TrioTim | 19 |
| *T1_MPRAGE_1mm_iso* | 256x256 | 176 slices | 2530.0 | 3.48 | 7.0 | NONE | 1.0 | 0.0 | 1.0 | NA | 256.0 | 256.0 | 1.000 x 1.000 | SIEMENS | TrioTim | 1 |
| *ge_func_2p1x2p1x4_PACE_improved* | 576x576 | 31 slices | 2000.0 | 30.0 | 90.0 | 2 | 4.0 | 9.99 | 4.40 | I | 200.0 | 200.0 | 2.083 x 2.083 | SIEMENS | TrioTim | 799 |
| *ge_functionals_PACE_wm* | 384x384 | 32 slices | 2000.0 | 30.0 | 90.0 | NONE | 4.0 | 9.99 | 4.40 | I | 200.0 | 200.0 | 3.125 x 3.125 | SIEMENS | TrioTim | 105 |
| *PRISMA-SMS2_TE30_2.0mm* | 832x832 | 52 slices | 2000.0 | 30.0 | 90.0 | 2 | 2.0 | 10.0 | 2.20 | A | 208.0 | 208.0 | 2.000 x 2.000 | SIEMENS | Prisma | 41 |
| *PRISMA-SMS3_TE30_2.0mm* | 936x936 | 72 slices | 2000.0 | 30.0 | 90.0 | 3 | 2.0 | 9.99 | 2.20 | A | 208.0 | 208.0 | 2.000 x 2.000 | SIEMENS | Prisma | 55 |

#### Preprocessing

fMRI data from all participants were analyzed using SPM12 (release 7487), CONN toolbox (Whitfield-Gabrieli & Nieto-Castanon, 2012) EvLab module (release 19b), and other custom MATLAB scripts. Each participant’s functional and structural data were converted from DICOM to NIFTI format. All functional scans were coregistered and resampled using the 4^th^ degree B-spline interpolation to the first scan of the first session (Friston et al., 1995). Potential outlier scans were identified from the resulting subject-motion estimates as well as from BOLD signal indicators using default thresholds in CONN preprocessing pipeline (5 standard deviations above the mean in global BOLD signal change, or framewise displacement values above 0.9 mm; Nieto-Castañón, 2020). Functional and structural data were independently normalized into a common space (the Montreal Neurological Institute [MNI] template; IXI549Space) using SPM12 unified segmentation and normalization procedure (Ashburner & Friston, 2005) with a reference functional image computed as the mean functional data after realignment across all timepoints omitting outlier scans. The output data were resampled to a common bounding box between MNI-space coordinates (−90, −126, −72) and (90, 90, 108), using 2mm isotropic voxels and 4^th^ degree B-spline interpolation for the functional data, and 1mm isotropic voxels and trilinear interpolation for the structural data. Last, the functional data were smoothed spatially using spatial convolution with a 4 mm FWHM Gaussian kernel.

fMRI data from the naturalistic cognition paradigms (Expts. 4a-b; Malik-Moraleda, Ayyash et al., 2022) included additional preprocessing steps prior to the spatial smoothing. First, to remove noise resulting from signal fluctuations originating from non-neuronal sources (for example, cardiac or respiratory activity), the first five BOLD signal timepoints extracted from the white matter and cerebrospinal fluid (CSF) were regressed out of each voxel’s time course. White matter and CSF voxels were identified based on segmentation of the anatomical image (Behzadi et al., 2007). Second, the residual signal was band-pass filtered at 0.008–0.09Hz to preserve only low-frequency signal fluctuations (Cordes et al., 2001).

Summary of representative structural (top) and functional (bottom) MRI sequences used across the 1023 scanning sessions. 12 scanning sessions did not include a structural scan (and instead relied on a previous structural scan collected from the same participant). Functional data were collected using a sequence other than one of the 6 detailed here for 23 of the scanning sessions. We note that the exact slice orientation and in plane rotation varied slightly by session, see OSF https://osf.io/y5t46/ for details. Abbreviations: IAA = image acquisition acceleration, ST = slice thickness in mm, SDP = slice distance percent, SD = slice distance in mm, AO = acquisition order (I=interleaved, A=ascending, SO = slice orientation, FR = FOV readout in mm, FPE = FOV phase encoding in mm, IPRe = in plane resolution, IPRo = in plane rotation.

#### First-level modeling

Responses to all experimental conditions from Expts. 1-3 (except Expt. 3c, see below) were estimated using the same modeling procedure. For each participant, responses in individual voxels were estimated using a General Linear Model (GLM) in which each experimental condition was modeled with a boxcar function convolved with the canonical hemodynamic response function (HRF) (fixation was modeled implicitly, such that all timepoints that did not correspond to one of the conditions were assumed to correspond to a fixation period). Temporal autocorrelations in the BOLD signal timeseries were accounted for by a combination of high-pass filtering with a 128 seconds cutoff, and whitening using an AR(0.2) model (first-order autoregressive model linearized around the coefficient a=0.2) to approximate the observed covariance of the functional data in the context of Restricted Maximum Likelihood estimation (ReML). In addition to experimental condition effects, the GLM design included first-order temporal derivatives for each condition (included to model variability in the HRF delays), as well as nuisance regressors to control for the effect of slow linear drifts, subject-specific motion parameters (6 parameters), and potential outlier scans (identified during preprocessing as described above) on the BOLD signal.

Responses to the 1,000 diverse sentences in Expt. 3c were modeled using GLMsingle (Prince et al., 2022), a robust framework for estimating responses to single-trial fMRI designs. A GLM was used to estimate the response amplitudes evoked by individual sentences (each sentence is treated a condition). As above, fixation was modelled implicitly. The data from different scanning sessions for a given participant were analyzed together. The beta weights returned by GLMsingle are in units of percent signal change (for more details, see: Tuckute et al., 2024; Prince et al., 2022).

#### Language Localizer

To identify cerebellar areas that are language-responsive, we use an extensively validated language ‘localizer’ paradigm (introduced by Fedorenko et al., 2010) that contrasts meaningful, well-formed sentences with a meaningless, perceptually-matched control stimulus. The sentences can either be written (e.g., Fedorenko et al., 2010), with lists of nonwords as the control condition (i.e., pseudowords that share orthographic properties with English but that carry no meaning and lack a syntactic frame), or auditorily (e.g., Malik-Moraleda, Ayyash et al., 2022) with foreign or muffled speech as the control condition (which share low-level acoustic properties with spoken language but, similar to nonwords, lack meaning and structure). This paradigm was designed to isolate language processing from lower-level perceptual processes and general cognitive demands. The localizer has been extensively used, in many variants (Fedorenko et al., 2010; 2016; Scott et al., 2017; Braga et al., 2020; Malik-Moraleda, Ayyash et al., 2022; Regev, Casto et al., 2024), and has been shown to reliably identify a network of brain regions in the neocortex that is i) highly selective for language (Fedorenko et al., 2011; Pritchett et al., 2018; Chen et al., 2019; Jouralev et al., 2019; Benn, Ivanova et al., 2023; Shain, Paunov, Chen et al., 2023), ii) robustly sensitive to linguistic manipulations at various scales (Fedorenko et al., 2010; 2016; Blank et al., 2016; Shain, Kean et al., 2024; Regev et al., 2024), iii) strongly interconnected (Saur et al., 2008; Friederici et al., 2009; Blank et al., 2014; Braga et al., 2020), and iv) causally important in linguistic ability (Goodglass, 1976; Wilson et al., 2023). For these reasons, we refer to this system as the “language network” (see Fedorenko et al., 2024 for a recent review) and use this localizer paradigm to search for language-responsive areas in the cerebellum. Every participant in the present study (regardless of the critical experiment that they performed) completed a language localizer at the begin of each scan.

#### Experiment 1a: Reading-based language localizer

Sentences and nonword lists were presented on the screen one word/nonword at a time. Several different versions of the localizer were used for the current set of participants (see Lipkin et al., 2022 for additional details), but the basic structure of the task and stimuli remained constant. We note that a small number of participants (n=13) completed a listening-based language localizer, but we included them in subsequent analyses to be consistent with the prior publication (Lipkin et al., 2022). We investigate the effect of presentation modality systematically below (see ‘Cerebellar Language Responses During Reading vs. Listening’).

#### Experiment 1b: listening-based language localizer

Participants listened to short passages (18 seconds long) from *Alice in Wonderland* in their native language, and—in the control condition—to acoustically degraded versions of the passages such that the speech was no longer intelligible. We used this dataset—over more closely matched, but smaller, datasets—to maximize our ability to accurately estimate response magnitudes and to increase power in our statistical comparisons. Generalizing the cerebellar language responses to speakers of typological diverse languages is an additional strength of using this particular dataset. The passages were degraded following the procedure introduced in Scott et al. (2017). See Malik-Moraleda, Ayyash et al. (2022) for more experimental details and discussion. Although all participants were highly proficient speakers of English, the majority (n=65) were not native English speakers and listened to passages in one of 45 languages (**Table 1**; for additional information about the participants’ language background, see OSF https://osf.io/y5t46/ for Suppl. Table 3 from Malik-Moraleda, Ayyash et al., 2022).

#### Functional Localization Approach

In the present study, we employ a within-participant functional localization approach that uses a ‘localizer’ paradigm to define functional regions of interest (fROIs) within individual participants. This approach accounts for known inter-participant variability in the precise anatomical locations of functional regions (e.g., regions that support language processing: Fedorenko et al., 2010; Frost & Goebel, 2012; Mahowald & Fedorenko, 2016). The first step is to identify areas where most participants show a response to the relevant contrast (e.g., Language>Control, see ‘Identifying common areas of language activation in the cerebellum’ below). These broad areas can then be used as spatial constraints, such that in each individual participant, the most responsive voxels within those areas are selected as fROIs (see ‘Definition of functional regions of interest (fROIs)’ below). The analyses described here were performed using the SPM_SS toolbox (available at https://www.evlab.mit.edu/resources).

#### Identifying common areas of language activation in the cerebellum

To identify spatially consistent areas of language activation, we used a group-constrained, subject-specific (GSS) analysis (Fedorenko et al., 2010). In this analysis, a set of ‘parcels’ (binary masks) was derived from a probabilistic overlap map of language-responsive voxels across the participants based on data from Expt. 1a (n=754). The probabilistic atlas was constructed by averaging over binarized maps of the most language-responsive voxels (i.e., top 10% across the whole brain) in individual participants. This probabilistic atlas was then smoothed using a 6 mm Gaussian kernel to avoid over-segmentation and only voxels in which at least 10% of the population showed a response were included; parcels were created using an image segmentation algorithm (the “watershed” algorithm; Meyer, 1991) (see Fedorenko et al., 2010 for additional details and discussion). This procedure yielded four common areas of language activation (‘parcels’) in the right posterior cerebellum (we note that these parcels differ slightly from the cerebellar language parcels published in a recent study, Wolna et al. 2025, due to small differences in how the GSS analysis was applied).

To allow for a more systematic comparison of language regions in the right and left cerebellar hemispheres, we constructed a symmetric version of the four cerebellar language regions by reflecting the right hemisphere parcels derived above across the cerebellar midline. Also, we note that although the GSS analysis described above provides a set of neocortical parcels as well, we chose to use previously generated publicly available neocortical language parcels (generated via a similar procedure on a smaller set of participants, and nearly identical the parcels generated here) for consistency with prior work (available at https://www.evlab.mit.edu/resources).

In addition to deriving a set of parcels from the whole-brain probabilistic atlas, we also created a set of parcels where the atlas was restricted to the cerebellum, and in the individual maps, we included the top 20% of language-responsive voxels within the cerebellum. This more liberal analysis yielded an additional parcel in right lobule IX, as well as four left cerebellar parcels that were roughly homotopic with the right cerebellar language regions derived above.

#### Definition of functional regions of interest (fROIs)

To define language functional regions of interest (fROIs) within individual participants, we selected the most language-responsive voxels (i.e., the top 10%) within each of the parcels above. This criterion has been used consistently across many prior studies in the Fedorenko Lab (e.g., Shain, Blank et al, 2020; Malik-Moraleda, Ayyash et al., 2022; Shain, Kean et al., 2024) and corresponds approximately to the whole-brain significance level of p<0.001. We have previously established that the fROIs defined based on the top 10% of voxels vs. using a fixed statistical significance threshold are near-identical (e.g., see Lipkin et al., 2022; Wolna et al., 2025). The advantage of the top 10% approach over a fixed statistical threshold approach is that it helps control for inter-individual variability in the overall strength of response and ensures that fROIs are of the same size across participants. Condition response magnitudes for a given fROI in a given participant reflect the univariate BOLD response, averaged over the voxels that comprise the fROI. Importantly, the responses to the localizer conditions are estimated using data that are independent from the data used to define the fROIs (a left out run).

For participants who completed multiple experiments and multiple versions of the language localizer, we used the same localizer scan to define their language fROIs across all experiments. Namely, we selected the most reliable language localizer scan available for a given participant across the period of data collection, where reliability was defined as the topographic consistency across runs (i.e., the spatial correlation between the odd and even runs of the paradigm). Spatial correlations here and elsewhere were calculated using the SPM_SS toolbox (available at https://www.evlab.mit.edu/resources), unless stated otherwise. Only language localizer scans with good cerebellar coverage were considered (see ‘Participant exclusion criteria’ above).

### Topography of Language Responses

#### Cerebellar flatmap visualization

Visualizations of language responses (and other contrasts of interest) on the cerebellar flatmap were performed using the Python (version 3.11.5, here and elsewhere) implementation of SUIT (SUITPy, version 1.3.1 here and elsewhere; Diedrichsen & Zotow, 2015). The default SUITPy parameters were used except twenty ‘depths’ were sampled (cf. the default six), and the ‘ignore_zeros’ argument was set to ‘True’ when visualizing parcels (i.e., binary masks) so that small parcels (e.g., LangCereb1, lateral VI) were visible on the surface. We emphasize that the flatmap was only used for visualization—all quantitative analyses were performed in the volume.

#### Topographic Variability Across Participants

To estimate the inter-individual topographic variability in the cerebellar (and neocortical, for comparison) language responses in Expt. 1a (n=754), we compared the pattern of language responses (Language>Control) between vs. within participants (pattern similarity within participants was computed between odd and even runs of the localizer). Similarity was defined as the Pearson correlation between unthresholded contrast maps, restricted to the cerebellar language parcels and the neocortical language parcels, for cerebellar and neocortical analyses, respectively. We then took the average of the n=753*2 between-participant correlations (for all possible participant/run pairings) to yield a single between-participant correlation value per participant. We then normalized the between-participant correlation value by the sum of the within- and between-participant correlations for that subject, giving us the percent of the total variance in a region that was shared across participants. We performed this procedure separately for the cerebellum and the neocortex. Topographic variability was computed in Python using the *nibabel* package (version 4.0.2, here and elsewhere), and Pearson correlations were calculated using the *scipy* package (version 1.11.3, here and elsewhere). The percent variance that was shared between participants in the cerebellum vs. the neocortex was statistically compared with the Wilcoxon signed-rank test (using the *scipy* package, here and elsewhere).

#### Lateralization

We defined language lateralization as:

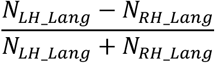

where *N* corresponds to the number of significant language voxels (Language>Control, t>2.576, p<0.01, uncorrected, two-sided t-test) in either the left or right cerebellar hemisphere (anatomical masks from Diedrichsen et al., 2009). The same procedure was applied to the neocortex. The significance threshold was made intentionally liberal to ensure that reliable responses in the cerebellum were not excluded due to the known lower signal quality in the cerebellum relative to the neocortex (Wang et al., 2025). Lateralization analyses were performed in Python using the *nibabel* package. We also computed a Pearson correlation between cerebellar and neocortical language lateralization across participants (n=754, Expt. 1a) using Python’s *scipy* package.

#### Overlap with Published Cerebellar Atlases

To quantify the overlap of our cerebellar language regions with two published cerebellar atlases, we first transformed our parcels (original in SPM IXI1549Space) into SUIT space. This transformation was performed with the *Functional_Fusion* package (version 1.0.0, publicly available at: https://functional-fusion.readthedocs.io/en/latest/index.html) and the *deform_data* function in Python. The transformation was performed for each region separately using linear interpolation, and the resulting parcels were binarized with a threshold of 0.2 (i.e., all values > 0.2 were set to 1). The individual parcels in SUIT space were then combined into a single file (available on OSF: https://osf.io/y5t46/). We then computed the overlap (Dice coefficient) between these parcels and two published cerebellar atlases: the Buckner et al. (2011) 7-network parcellation and the Nettekoven et al. (2024) symmetrical 32-region parcellation. Images for calculating Dice coefficients were loaded in Python using the *nibabel* package. The outline of our language parcels was overlayed with the published cerebellar atlases using Adobe Illustrator, and was only intended for qualitatively comparing the locations of our language regions with prior results.

We would also like to emphasize that our parcels are designed to be used in conjunction with functional localization within individuals. The parcels are, by design, relatively large, and contain not only language-responsive voxels, but also voxels that likely belong to distinct functional networks. As a result, using the full set of voxels in each parcel would drastically underestimate language selectivity and make it more difficult to decipher the linguistic areas’ computations.

#### Signal-to-Noise Ratio (SNR)

We computed a measure of the signal quality in each of the cerebellar language regions and the neocortical language network. In particular, for every participant in Expt. 1a and every cerebellar (and neocortical) fROI, we divided the observed response in a given region to the Sentence condition from the language localizer (a measure of the experimentally-induced signal fluctuations) by the residual sum-of-squares from the first-level GLM (a measure of the signal that could not be explained by the task). The residual sum-of-squares values were averaged over all voxels in the fROI. The resulting value represents the ratio of the task-relevant “signal” to the task-independent “noise”. This measure was chosen over the more commonly used tSNR measure as tSNR is less well-suited for task-based timeseries (see Welvaert & Rosseel, 2013, for additional discussion). SNR values were statistically compared between the neocortical language network and the cerebellar language regions with the Wilcoxon signed-rank test.

#### Reading vs. Listening

To systematically evaluate the similarity of cerebellar responses during reading and listening, we took two complementary approaches using the participants from Expt. 1b that completed both a reading- and listening-based language localizer (n=85, data from Malik-Moraleda, Ayyash et al., 2022). First, and most importantly, we tested whether the cerebellar language regions that were identified using a reading-based localizer (using the reading-derived language parcels) were also responsive during listening (over and above a low-level auditory control condition). To do so, we used linear mixed-effects (LME) models (one per cerebellar language region). Here and elsewhere, LME models were fit using the *lmer* function from the *lme4* package (version 1.1-35.5) in R (version 4.4.1). The LME models included two binary, treatment-coded fixed effects: ‘Effect’ (0 for the control conditions, 1 for language) and ‘Response’ (0 for the responses to the reading-based localizer, 1 for the responses to the listening-based localizer). The model included each of these individual terms and their interactions. The dependent variable for all LME models (here and elsewhere, unless indicated otherwise) was the response magnitude (% BOLD signal change) in a given subject-specific fROI, The LME models were fit with restricted maximum likelihood estimation (here and elsewhere, unless indicated otherwise) and included ‘Participant’ as a random effect.

Second, we computed the spatial correlations in the pattern of reading-vs. listening-responsive voxels at the group level and across individual participants. To compare patterns at the group level, a spatial correlation was computed between the probabilistic overlap maps from the two modalities (as defined above, see ‘Identifying common areas of language activation in the cerebellum’). A similar procedure was performed at the level of individual participants. Namely, spatial correlations were taken between i) runs of the reading-based localizer, ii) runs of the listening-based localizer, iii) runs of the reading- and listening-based localizers (averaging across all combinations of individual runs to match the amount of data between i and ii). This analysis was performed for each of the cerebellar language regions separately. Spatial correlations for the group- and individual-level comparisons were computed using Python’s *scipy* package.

We observed that the individual-level spatial correlations across reading vs. listening were similar to the correlations between runs of the listening-based localizer. To evaluate the statistical reliability of this similarity, we compared these correlations to the geometric mean of the reading-reading and the listening-listening correlations, evaluated with a one sample Wilcoxon signed-rank test. Under the null hypothesis that the two patterns of activation are exactly the same (reading and listening), we would expect the reading-listening correlations to be equal to the geometric mean of the within-modality correlations, accounting for noise differences between the modalities.

### Non-linguistic Experiments

#### Selection of experiments

To evaluate selectivity for language, we broadly sampled non-linguistic tasks, for which data were available in our lab’s database, that i) had been argued in prior work to engage the cerebellum (e.g., speech articulation, Ackermann et al., 2007; social perception, D’Mello et al., 2015; and semantic processing, LeBel et al., 2021; D’Mello et al., 2020b), and/or ii) were hypothesized in prior work to share cognitive and neural resources with language due to putatively similar computational demands (e.g., the processing of hierarchical structure in the case of music, Lerdahl & Jackendoff, 1977; suppressing irrelevant representations in the case of cognitive control mechanisms, Novick et al., 2005; etc.). The latter claims have typically been made for language in general, not considering language processing in the cerebellum specifically, and they do not find support with respect to the neocortical language areas (see Fedorenko et al., 2024 for a recent review); nevertheless, we wanted to evaluate these hypotheses for the cerebellar language regions to see whether their profiles may differ in meaningful ways from those of the neocortical language areas.

#### Experiment 2a: Articulation Task

Participants were instructed to repeat a sequence of 4 syllables (“ba, ga, ra, da”) or a simple finger-tapping motor sequence. Participants were shown an instruction slide for 2 seconds before each block which indicated the task that they were supposed to perform. During syllable production, a flashing icon of a mouth was presented on the screen. During finger tapping, an icon of a hand was shown with flashing indicators on top of the fingers. See Wolna et al. (in prep) for more experimental details and discussion.

#### Experiment 2b-i: Math Task

Participants were shown 2 numbers to add together. As with all the cognitively demanding tasks (Expt. 2b), the task contained both an Easy and a Hard condition. In the Easy condition, both numbers were single-digit, and in the Hard condition one number was single-digit, and the other was double-digit. Participants then had to select the correct sum in a forced-choice question with 2 options. Participants received feedback if they got the questions right or wrong. See Malik-Moraleda, Ayyash et al. (2022) for more experimental details and discussion.

#### Experiment 2b-ii: Spatial Working Memory Task

Participants were presented with a 3×4 grid and asked to keep track of where blue squares appeared. In the Easy condition, a sequence of 4 blue squares (one at a time) was shown; and in the Hard condition, a sequence of 8 blue squares (two at a time) was shown. Participants then had to select the correct cumulative set of blue square locations in a forced-choice question with two options. Participants received feedback if they were right or wrong. See Fedorenko et al. (2011) for more experimental details and discussion.

#### Experiment 2b-iii: Verbal Working Memory (Digit Span) Task

Participants were asked to keep track of a list of digit names (written out as words: e.g., “two”). In the Easy condition, a sequence of 4 digit names (one at a time) was shown, and in the Hard condition, a sequence of 8 digit names (two at a time) was shown. Participants then had to select the correct set of digits in a forced-choice question with two options. Participants received feedback if they were right or wrong. See Fedorenko et al. (2011) for more experimental details and discussion.

#### Experiment 2b-iv: Multi-Source Interference Task (MSIT)

Participants were presented with triplets of digits (each between 0 and 3) and asked to press the button (1, 2, or 3, laid out horizontally on the button box) that corresponded to the identity of the non-repeated digit. In the Easy condition, the identity of the digit was the same as its position and the distractors were not a possible button value (e.g., 100). In the Hard condition, the identity and the position of the digit did not match and the distractors were a possible button value (e.g., 212). See Bush & Shin (2006) and Fedorenko et al. (2011) for more experimental details and discussion.

#### Experiment 2b-v: Verbal Multi-Source Interference Task (vMSIT)

Participants were presented with triplets of words (‘none’, ‘left’, ‘middle’, ‘right’) and asked to press the button (left, middle, or right, laid out horizontally on the button box) that corresponded to the identity of the non-repeated word. In the Easy condition, the identity of the word was the same as its position and the distractors were not a possible button value (e.g., “none none right”). In the Hard condition, the identity and the position of the word did not match and the distractors were a possible button value (e.g., “right left left”). See Fedorenko et al. (2011) for more experimental details and discussion.

#### Experiment 2b-vi: Stroop Task

Participants were presented with a word and asked to overtly name the color of the word’s font. In the Easy condition, the words were non-color adjectives (e.g., close, huge), and in the Hard condition, the words denoted colors. The non-color adjectives were matched to the color adjectives in length and lexical frequency. In half of the hard trials, the color words did not match their font color. See Fedorenko et al. (2011) for more experimental details and discussion.

#### Experiment 2c-i: Music Perception A

Participants listened to 64 unfamiliar musical clips that were extracted from 1950s/1960s pop and rock music. In addition to the intact musical clips, participants heard versions of the excerpts that were scrambled in various ways to disrupt the musical structure. 8 of the participants were asked to passively listen (and press a button after each excerpt), and the remaining 4 participants were asked “How much do you like this piece?” after each excerpt and had to rate it on a scale from 1 (not at all) to 4 (very much) by pressing a button on the button box. See Fedorenko et al. (2011) for more experimental details and discussion.

#### Experiment 2c-ii: Music Perception B

Participants passively listened to 9-second audio clips of critical music stimuli—including orchestral music from classical orchestras or jazz bands, single-instrument music (e.g., cello, flute, guitar, piano, etc.), and drum music—or control, non-music stimuli (animal vocalizations or pitched environmental sounds). See Chen et al. (2023) for more experimental details and discussion.

#### Experiment 2d: Dynamic Visual Social Perception

Participants passively viewed 3-second videos of critical social stimuli (faces and body parts) or control stimuli (scenes, objects, and scrambled objects). The videos of faces and bodies depicted children playing in front of a black background that were framed close-up to only show their face or body. The videos of scenes were mostly taken from a car while driving slowly through leafy, pastoral suburbs. The videos of objects depicted small man-made objects that were chosen to avoid appearing animate. The actor moving the object was also limited from view. The scrambled object videos were constructed by splitting the video into a 15×15 grid and scrambling the location of the squares within the grid. See Pitcher et al. (2011) and Pritchett et al. (2018) for more experimental details and discussion.

#### Experiment 2e-i: Naturalistic Silent Movie

Participants passively viewed a silent 5-minute animated film that was rich with semantic, including social content. The film—Partly Cloudy (Pixar Animation Studios)—has been used in prior work as a nonverbal ‘Theory of Mind’ (ToM) localizer (see Jacoby et al., 2016; Richardson et al., 2018; 2020; Paunov et al., 2019; Kamps et al., 2022). Sections of the film were then coded as those that i) elicit mental state content (‘mental’ condition, 4 events, 44 seconds total), ii) depict physical events (‘physical’ condition, 3 events, 24 seconds total), iii) depict interactions between characters without a mental inference component (‘social’ condition, 5 events, 28 seconds total), and iv) depict the characters experiencing physical pain (‘pain’ condition, 7 events, 26 seconds total; the localizer is publicly available at https://saxelab.mit.edu/theory-mind-and-pain-matrix-localizer-movie-viewing-experiment/). See Jacoby et al. (2016) and Shain, Paunov, Chen et al. (2023) for more experimental details and discussion.

#### Experiments 2e-ii-A, 2e-ii-B, and 2e-ii-C: Event Semantics Tasks

Participants were shown pictures of events or linguistic event descriptions and asked to perform a semantic or perceptual task. In Expt. 2e-ii-A, participants were shown versions of agent-patient interactions that were either plausible or implausible (e.g., “The jester is entertaining the king.” vs. “The king is entertaining the jester.”). In Expt. 2e-ii-B, participants were shown versions of people interacting with everyday objects that were either plausible or implausible (e.g., “The man is peeling a carrot.” vs. “The man is peeling a candle.”). In both of these experiments, participants had to make a plausiblity judgment in the critical, semantic task. In Expt. 2e-ii-C, participants were shown versions of people performing actions on everyday objects, with the actions being reversible (i.e., they could be undone) or irreversible. For example, eating a clementine is irreversible while putting stables into a stapler is reversible. Participants had to make a reversibility judgment in the semantic task. The perceptual task for two of the experiments required participants to indicate the direction that the stimuli were moving (right or left), and for the other experiment, the task required participants to indicate whether the movement direction of the stimuli changed 3 or 4 times. See Ivanova et al. (2021) and Ivanova (2022) for more experimental details and discussion.

#### Statistical evaluation of non-linguistic responses

We statistically evaluated the non-linguistic responses relative to i) the fixation baseline, ii) the response to the control condition from the language localizer (nonword lists), and iii) the critical, language condition from the language localizer (sentences) using linear mixed-effects (LME) models. To evaluate the responses relative to the fixation baseline, we used an LME model with one dummy-coded predictor: ‘Effect’ (with 32 levels, one for each of the non-linguistic conditions). To evaluate the responses relative to the control or critical condition from the language localizer, we used an LME model with one treatment-coded predictor: ‘Effect’ (with 33 levels, one for each of the non-linguistic conditions and the control or critical condition from the language localizer which served as the reference level). All models included ‘Participant’ as a random effect, and models for the neocortical regions also included ‘ROI’ as a random effect (for the five ‘core’ left hemisphere language regions located in posterior temporal cortex, anterior temporal cortex, inferior frontal gyrus, orbital part of the inferior frontal gyrus, and middle frontal gyrus, here and elsewhere, Fedorenko et al., 2024).

We additionally tested for between-condition differences within certain non-linguistic domains. All within-domain comparisons were evaluated with LME models that all included ‘Participant’ and (for the neocortical language network only) ‘ROI’ as random effects. In particular, we tested i) whether there were stronger responses during articulation than during finger tapping (Expt. 2a), ii) whether there were overall stronger responses to the harder than the easier conditions in the cognitively demanding tasks (Expt. 2b; this model included an additional random intercept for ‘Experiment’: spatial working memory, verbal working memory, etc.), iii) whether there were stronger responses when listening to structured musical excerpts than to unstructured/non-music sounds (Expt. 2c), iv) whether there were stronger responses to visual social stimuli (faces and bodies) than non-social stimuli (scenes and objects, Expt. 2d), v) whether there were stronger responses to mental than physical content (Expt. 2e-i), vi) whether there were stronger responses to social than pain content (Expt. 2e-i), and vii) whether there were stronger responses during the event semantic task than the perceptual task (Expt. 2e-ii). All comparisons were modeled as binary, treatment-coded fixed effects. Comparisons were evaluated for each cerebellar ROI separately.

#### Measuring selectivity for language

We measured selectivity for language relative to the non-linguistic conditions using a standard definition of selectivity. In particular, we defined selectivity as the difference between a region’s response to language and the next highest response (to a non-linguistic condition), scaled by the sum of the two responses. A value of 1 would indicate that a region is entirely selective for language (i.e., it does not respond above baseline to any other task), and a value of 0 would indicate that a region responds as strongly to a non-linguistic condition as it does to language.

#### Dissociating cerebellar language regions from nearby cognitive and motor networks

Given the proximity of different functional networks to one another in the cerebellum (Nettekoven et al., 2024; Saadon-Grossman et al., 2024), we examined the degree to which the cerebellar language regions were dissociable from three nearby cognitive and motor networks: the Theory of Mind network (ToM, Saxe and Kanwisher, 2003), the Multiple Demand (MD) network (Duncan et al., 2010), and the motor articulation network (Wolna et al., in prep). To that end, we first defined a set of cerebellar ToM parcels using the False Belief>False Photo contrast from Expt. 2f (below), cerebellar MD parcels using the Hard>Easy contrast from the spatial working memory task (Expt. 2b-ii), and cerebellar articulation parcels using the Articulation>Finger Tapping conditions from the articulation task (Expt. 2a). We defined these parcels using the GSS analysis described above (see ‘Identifying common areas of language activation in the cerebellum’). We restricted the GSS analysis to the cerebellum only and used the top 10-20% of voxels within individual participants to construct the probabilistic overlap map (top 20% for the ToM and MD GSS analysis, top 10% for the articulation GSS analysis). We also note that because we need an independent run of data to estimate the fROIs’ responses, we excluded participants who only had a single run of each of these tasks, leaving 156 participants for the ToM task and 574 participants for the MD task (all 33 participants from the articulation task had multiple runs).

We then identified the cerebellar language regions (including both right and left cerebellar regions, as well as right lobule IX) that showed overlap with the ToM, MD, or articulation parcels. In total, three language parcels overlapped with the ToM parcels (right and left Crus I/II/VIIb, and right IX), two language parcels overlapped with the MD parcels (right lateral VI and right VIIb/VIIIa), and one language region overlapped with the articulation parcel (right VIIb/VIIIa). For each of these overlap cases, we constructed a new parcel that was the union of the original language parcel with the overlapping ToM/MD/articulation parcel (e.g., the union of the language and MD parcels in right lateral VI). Finally, in each of these combined parcels, we defined two fROIs (based on the language contrast or based on the ToM/MD/articulation contrast) and measured the responses to language and the other task.

Finally, to statistically evaluate whether language and ToM/MD/articulation fROIs were dissociable, we used a linear mixed-effect (LME) model (one per combined parcel) with three binary, sum-coded fixed effects: ‘Preference’ (0.5 the preferred conditions: sentences, false belief vignettes, hard spatial working memory, or articulation; −0.5 for non-preferred conditions: nonword lists, false photo vignettes, easy spatial working memory, or finger tapping), ‘Response’ (0.5 for the responses to the language localizer, −0.5 for the responses to the ToM, MD, or articulation localizer), and ‘Localizer’ (0.5 for when the language localizer was used to identify the fROI, −0.5 for when the ToM, MD, or articulation localizer was used). The model included each of these individual terms and their interactions. The LME models were fit with maximum likelihood estimation and included ‘Participant’ as a random effect.

#### Experiment 2f: Theory of Mind

Participants were presented with written vignettes that describe a situation where an agent holds a false belief about the world (e.g., someone having a false belief about the location of an object, the critical condition) and linguistically similar vignettes that describe a situation where a visual depiction of the world became outdated (e.g., a photo of a house that has since been demolished, the control condition). See Saxe and Kanwisher (2003) and Dodell-Feder et al. (2011) for more experimental details and discussion (the localizer is available at: http://saxelab.mit.edu/use-our-efficient-false-belief-localizer).

### Linguistic Experiments

#### Experiments 3a-i, 3a-ii, and 3a-iii: Language Comprehension Tasks

In three similar language comprehension experiments, participants passively read sentences (S), word lists (W), Jabberwocky sentences (J; sentences where content words are replaced by nonwords, such as “florped” or “blay”), and nonword lists (N). The experiments followed a 2×2 design, manipulating lexical access (which we sometimes refer to as “sensitivity to individual word meanings”) and syntactic structure building (which we sometimes refer to as “sensitivity to grammatical structure”). In particular, reading sentences requires both lexical access and syntactic structure building; reading word lists or Jabberwocky sentences requires only one of these processes (lexical access and syntactic structure building, respectively); and reading nonword lists requires neither process (but engages the same visual, orthographic, and attentional processes). Finally, if there is an interaction between lexical access and syntactic structure building—such that the response when both components are present (i.e., during sentences) is greater than the sum of the two components individually—then this indicates that a region is sensitive to compositional semantics (we refer to this in the main text as “processing sentence-level meanings”). This paradigm has been widely used in prior work for its ability to carve language into its component processes (Fedorenko et al., 2010; 2016; Kauf et al., 2024; Shain, Kean et al., 2024; Regev, Casto et al., 2024). Different items were used in each of the three experiments, but they were all 8 or 12 words/nonwords in length. See Fedorenko et al. (2010), Kauf et al. (2024), and Shain, Kean et al. (2024), for more experimental details and discussion.

#### Experiment 3b-i: Spoken Language Production Task

Mirroring the design of the language comprehension experiments, participants were asked to produce sentences, word-lists, and nonword-lists, while keeping their heads as still as possible. In the sentence production condition, participants were shown photographs of common events and asked to produce a description of the event (e.g., “The man is surfing.”). In the word list condition, participants were shown groups of 2-4 photographs of inanimate objects and asked to name each of them (e.g., “fruit, ball”). Objects were grouped in a way so as to minimize semantic associations between the objects to isolate the processes related to single-word production. In the nonword list condition, participants were shown 4 monosyllabic nonwords and asked to read them aloud (e.g., “bolt, sloal, sneaf, tworce”). See Hu, Small et al. (2023) for more experimental details and discussion.

#### Experiment 3b-ii: Typed Language Production Task

The design of the Expt. 3b-ii was identical to that of 3b-i, except that participants were instructed to type, rather than speak, their responses using a scanner-safe keyboard. This experiment was designed to test the generalizability of the results from the spoken version of this paradigm to another output modality (as the high-level demands of language production, such as lexical access and syntactic structure building, are present both when speaking and when typing). A similar, but nonoverlapping, set of materials that were presented to participants in Expt. 3b-i were presented here. See Hu, Small et al. (2023) for more experimental details and discussion.

#### Statistical evaluation of linguistic responses

As with the non-linguistic conditions, we statistically evaluated the linguistic responses relative to i) the fixation baseline, ii) the response to the control condition from the linguistic experiments (nonword list comprehension or production), and iii) the critical, language condition from the linguistic experiments (sentence comprehension or production) using linear mixed-effects (LME) models (see ‘Statistical evaluation of non-linguistic responses’ above for model details).

For the language comprehension experiments, we also statistically evaluated whether there was an interaction between lexical semantic and syntactic processing—which would indicate that a region is sensitive to the demands of semantic composition above and beyond lexical access and syntactic structure building demands. To do this, we used nested LME models and the likelihood ratio test. In particular, all models included two binary, treatment-coded fixed effects: ‘lexical’ (1 for conditions that required lexical access: sentences and word lists; 0 for conditions that did not: Jabberwocky sentences and nonword lists), and ‘syntax’ (1 for conditions that required syntactic structure building: sentences and Jabberwocky sentences; 0 for conditions that did not: word and nonword lists). Two models were fit per region—one with only these fixed effects, and another with these fixed effects plus their interaction. The nested LME models both included ‘Participant’ and ‘Experiment’ (i.e., Expt. 3a-i, ii, or iii) as random effects, and were fit with maximum likelihood estimation. The models for the neocortical language network additionally included ‘ROI’ as a random intercept (for the five ‘core’ left hemisphere language regions). We then evaluated the improvement in model performance with the addition of the ‘lexical:syntax’ interaction term using the likelihood ratio test (with R’s *anova* function).

#### Statistical comparison of linguistic responses in the neocortical language network vs. LangCereb3

To statistically compare the responses to the comprehension experiments (i.e., Expt. 3a only) in the neocortical language network vs. LangCereb3, we fit an LME model similar to the model described above, but with the addition of another binary, treatment-coded fixed effect: ‘cortex’ (0 for LangCereb3, 1 for the neocortical language network). The neocortical responses used in the model were the average over the five ‘core’ left hemisphere language regions. To account for the signal quality difference between the neocortical language network and LangCereb3, BOLD responses (the dependent variable) were normalized by the average response to the “S” condition (across participants) for a given region prior to model fitting. This normalization was applied for each of Expt. 3a’s sub-experiments separately (i.e., 3a-I, 3a-ii, 3a-iii). The LME model included ‘Participant’ as a random effect.

#### Experiment 3c: Responses to 1,000 diverse sentences

##### Materials

Participants were exposed to 1,000 linguistically diverse sentences that were all six words in length. The sentences were extracted from diverse naturalistic text corpora and the selection procedure was two-fold. First, 534 of the 1,000 sentences were selected to be semantically diverse by ensuring that sentences were close in GloVe space (Pennington et al., 2014) to 180 target semantic clusters (identified in Pereira et al., 2018). Second, the remaining 466 sentences were randomly sampled from eight stylistically diverse corpora (e.g., a subset of the Common Craw C4 corpus, the corpus of contemporary American English, etc.) in a stratified manner (see Tuckute et al., 2024 for additional details about the materials). All participants completed two 2-hour scanning sessions. Participants were instructed to read attentively and think about the sentence’s meaning, and they performed a short memory task after the end of the scan (outside of the scanner) to encourage engagement with the stimuli.

##### Linguistic and semantic features

We obtained a wide variety of features for each of the 1,000 sentences (per Tuckute et al. 2024). We considered three measures of surprisal (i.e., log probability): GPT2-XL-based surprisal, which considers the entire sentence and is bases on linguistic and semantic patterns learned in large amounts of text, 5-gram surprisal, which reflects predictability based on the sequences of particular words, and a probabilistic context-free grammar (PCFG) surprisal, which reflects the predictability of part-of-speech patterns. We also considered behavioral ratings from human participants. These ratings included how plausible a sentence is (“rate the sentence for whether it makes sense on a scale from 1 (does not make any sense) to 7 (makes perfect sense)”), how grammatically well-formed it is (“rate the sentence for how well it follows English grammar rules on a scale from 1 (completely ungrammatical) to 7 (perfectly grammatical)”), its perceived frequency (“rate each sentence according to how likely you think you are to encounter this sentence on a scale from 1 (not at all likely) to 7 (very likely)”), and its perceived frequency in conversational settings (“rate each sentence according to how likely you think it is to occur in a conversation between people on a scale from 1 (not at all likely) to 7 (very likely)”)—all measures related to a sentence’s processing difficulty (in addition to the surprisal-based measures). See Tuckute et al. (2024) for additional details and methods relating to the sentence-level features. To quantitatively evaluate the how strongly different sentence properties modulate responses in LangCereb3 (Crus I/II/VIIb) and the neocortical language network, we computed the partial correlation between responses in the two regions and each of the sentence properties individually (controlling for the effect of GPT2-xl surprisal). Partial correlations were calculated with the *pinguoin* package in Python (version 0.5.5).

We also statistically compared the similarity of responses to these 1,000 sentences in LangCereb3 and the neocortical language network, without respect to specific linguistic features. In particular, we compared the between-region similarity across participants (i.e., the correlation in responses across the two regions for all possible participant/region pairings) to the geometric mean of the two within-region similarities (computed across all participant pairings). If the pattern of response is the same in the two regions, then we would not expect the between-region correlations to differ from the geometric mean of the within-region reliabilities, which serves as a theoretical upper limit (taking into account the signal quality differences in the two regions). The between-region correlations were evaluated against the geometric mean of the within-region correlations with a one-sample Wilcoxon signed-rank test.

#### Experiment 3d: Social vs. Nonsocial Language Localizer

We designed two new versions of the language localizer to test if the content of the sentences affects the topography of the regions that were identified using our standard localizer.

To identify highly social and nonsocial sentences, we first collected behavioral ratings on a diverse set of 1,432 sentences (pooled from across three sets of experimental materials) for how “social” the content of the sentence is (48 sentences from the original language localizer, Expt. 1a; 1,000 sentences from Expt. 3c, and 384 sentences from a third experiment, items from Pereira et al., 2018). In particular, we asked online participants (via the Prolific platform for behavioral data collection) to rate each sentence for how much it made them think of other people’s experiences, thoughts, beliefs, desires, and/or emotions on a scale from 1 (not at all) to 5 (very much) (all 1,432 sentences and the associated behavioral ratings are now available on OSF https://osf.io/y5t46/). Each sentence was rated by 18-41 participants, and inter-rater agreement was high (average split-half reliability across 1,000 iterations: r=0.91±0.002 s.d. Expt. 1a; r=0.84±0.025 s.d. Expt. 3c; r=0.78±0.022 s.d. third experiment), suggesting a high degree of agreement on which sentences are more vs. less social in content, and substantial variability among the sentences.

We then used these ratings to train an *encoding model* to learn the mapping between a sentence’s embedding from a large language model (GPT2-xl, Radford et al., 2019) and its socialness rating. If successful, such an encoding model could be used to predict the socialness rating for new, unseen sentences. To fit this encoding model, we extracted sentence embeddings from all layers of GPT2-xl for the 1,432 sentences for which we had collected behavioral ratings. We extracted the model embeddings from the last token in the sentence (following Tuckute et al., 2024). We then selected the model layer that best predicted the socialness ratings using ridge regression with 5-fold cross validation. Performance was evaluated by computing the Pearson correlation between predicted and actual socialness ratings on the test set (and then averaged over the 5 folds). The regularization parameter (alpha) was learned using leave-one-out cross validation on the sentences in the training set. Therefore, the test sentences that were used to evaluate the model’s performance were not incorporated into the procedure for identifying alpha or learning the regression weights (i.e., the procedure was fully cross-validated). Layer 27 was the best performing GPT2-xl layer (r=0.87 predicted vs. actual) and was used in all subsequent analyses.

Using the sentence embeddings from this layer, we then learned the final encoding model across 1,432 sentences. The regularization parameter (alpha) for the ridge regression was again learned using leave-one-out cross validation, and we selected the regularization parameter with the lowest squared error of 60 logarithmically-spaced potential values (1×10−30, 1×10−29, …, 1×10^28, 1×10^29; following Tuckute et al., 2024).

We next extracted sentence embeddings for 21,230 new sentences from across 7 diverse corpora: the Brown corpus (1,127 sentences), a subset of the Universal Dependencies (UD) corpus (465 sentences), a subset of articles published in the Wall Street Journal (WSJ) (5,410 sentences), and four years of the Corpus of Contemporary American English (COCA) (2009: 3,896 sentences; 2010: 4,060 sentences; 2011: 4,031 sentences; 2012: 2,241 sentences). These 21,230 sentences were restricted to be 12 words long (to keep the stimuli as similar to the original language localizer as possible) and did not include i) digits, ii) uppercase letters (other than at the beginning of the sentence), or iii) special characters (e.g., ‘$’, ‘%’, etc.) and quotations. We then used the encoding model fit above to predict the socialness ratings for all 21,230 sentences. There was substantial variability across sentences in their predicted socialness ratings.

To select a subset of highly social and nonsocial sentences to be used in the new versions of the language localizer, we took a subset of the total 21,230 sentences that had a rating above the 97.5 percentile (for social sentences, n=531) or below the 2.5 percentile (for nonsocial sentences, n=531). We then randomly selected 60 sentences from the 531 highly social sentences, along with the 60 sentences from the nonsocial sentences that were most closely matched to these items on *surprisal* (known to affect responses in the language areas). Sentence-level surprisal was extracted from GPT2-xl by averaging the surprisal estimates across all tokens in the sentence. The 60 sentences in the two sets were then manually pruned to 48 final sentences (available on OSF https://osf.io/y5t46/), removing e.g., items with typos, items that discussed potentially triggering content, and items with stray characters that were not filtered in the initial preprocessing (e.g., dashes). Finally, behavioral ratings were collected from online participants on the two sets of sentences to ensure that the model predictions were accurate. There was high agreement between the model’s predicted social ratings for these items and the ratings of the new online participants (r=0.965), and the two sets of stimuli were robustly different in their socialness ratings.

We then collected fMRI data from 10 participants who completed four total runs of the language localizer— two with the highly social sentences and two with the highly nonsocial sentences. The experimental design was identical to Expt. 1a: sentences were presented one word at a time, with lists of nonwords interspersed, and the participants were instructed to passively read the sentences and press a button when they saw an image of a hand pressing a button. Then, for each participant, we identified the most language-responsive voxels (i.e., top 10%) within each of the four cerebellar parcels separately using the two sets of stimuli (our standard localization procedure). To quantify whether the functional regions of interest (fROIs) differ depending on the set of sentences that were used in the localizer, we computed the Dice coefficient between the fROIs defined by the social sentences and the fROIs defined by the nonsocial sentences. We then compared these coefficients to the Dice coefficients of the fROIs defined using odd vs. even runs of the same set of sentences, controlling for the amount of data.

### Naturalistic Cognition Paradigms

#### Experiment 4a: Resting State

Participants were instructed to close their eyes and let their mind wander for five minutes while trying their best to stay awake (e.g., as in Blank et al., 2014). The projector was turned off and the lights in the room were dimmed. We note that the participants were the same as those in Expt. 1b (n=85, and Expt. 4b below).

#### Experiment 4b: Spoken Naturalistic Story

Participants (the same as in Expt. 1b and 4a) listened to one of 3 ∼4.5-minute passages from *Alice in Wonderland* in their native language.

#### Computing functional correlations

To evaluate the degree of functional integration between cerebellar language regions and the neocortical language network, we calculated functional correlations during a resting state scan (Expt. 4a) and during a spoken naturalistic story (Expt. 4b). To compute functional correlations, fROI-level BOLD timeseries were first defined by averaging the BOLD timeseries across the top 10% of language-responsive voxels in a given fROI (see ‘Definition of functional regions of interest (fROIs)’). Timeseries consisted of timepoints from the entire duration for the resting state scan. For the story, the timeseries consisted of timepoints starting 6 seconds after the story began to account for the hemodynamic response and did not include timepoints during fixation (12 seconds at the beginning and end of the run). Then, the Pearson correlation was computed between all pairs of cerebellar and neocortical language regions (83 pairs in total) within an individual participant. Correlations were computed using the *scipy* package in Python. Left and right hemisphere ‘core’ language regions were included in the analysis. When creating summary plots (e.g., the average functional correlation between a given cerebellar language region and all left hemisphere neocortical language regions) correlations between pairs of regions were averaged within, rather than across, participants.

#### Statistical evaluation of functional correlations

We statistically evaluated the functional correlations in four ways using linear mixed-effects (LME) models. First, we tested whether the right cerebellar language regions were overall more correlated with the left-that the right-hemisphere neocortical regions using an LME with one binary, treatment-coded fixed effect: ‘hemisphere’ (0 for functional correlations with the left neocortical language network, 1 for functional correlations with the right neocortical language regions). The model included random intercepts for ‘Participant’, ‘cerebROI’ (for the four cerebellar language regions), ‘dataset’ (naturalistic story or resting state), and ‘ROI’ (for the five ‘core’ left hemisphere language regions).

Second, we tested whether functional correlations between the right cerebellar language regions and the left-hemisphere neocortical language network were higher during story listening than rest. The LME model that was used to test this claim included two treatment-coded fixed effects: ‘cerebROI’ (with four levels corresponding to each of the cerebellar language regions), and ‘dataset’ (0 for resting state, 1 for the naturalistic story). The model included random intercepts for ‘Participant’ and ‘ROI’ (for the five ‘core’ left hemisphere language regions), with random slopes for ‘Participant’ and ‘ROI’ for the ‘dataset’ fixed effect. We then estimated the difference in functional correlations between the datasets post hoc using the *emmeans* package (version 1.10.5) in R. We also used this model to test our third claim—whether LangCereb3 (Crus I/II/VIIb) was more functionally correlated with the left-hemisphere neocortical language network compared to the correlation between other cerebellar language regions and the neocortical network. We estimated the difference between pairs of cerebellar regions using the *emmeans* package and the resulting p-values were adjusted using the Holm correction.

And fourth, we tested whether functional correlations were higher between LangCereb3 (Crus I/II/VIIb) and the frontal vs. temporal left-hemisphere neocortical language regions. The LME model that was used to test this claim included two binary treatment-coded fixed effects: ‘lobe’ (0 for functional correlations with frontal language regions, 1 for functional correlations with temporal language regions) and ‘dataset’ (0 for resting state, 1 for the naturalistic story). The model included random intercepts for ‘Participant’ and ‘ROI’ (for the five ‘core’ left hemisphere language regions). The random-effects structures from all LME models in this section were selected by iteratively removing terms from a maximal random-effects structure until the fit was no longer singular, and all models were fit with restricted maximum likelihood estimation.

## Data Availability

Preprocessed data needed to reproduce the figures is available on OSF (https://osf.io/y5t46/).

## Code Availability

Code used to conduct analyses is available on GitHub (https://github.com/coltoncasto/cerebellum_PUBLIC).

## Acknowledgements

We acknowledge the Athinoula A. Martinos Imaging Center at the McGovern Institute for Brain Research at MIT, and its support team (Steve Shannon and Atsushi Takahashi), the many participants who took part in EvLab studies over the years, making this study possible, and the many former and current EvLab members who collected the data used in this study, especially Sara Swords, Anya Ivanova, and Carina Kauf. We also thank Nancy Kanwisher, Randy Buckner, Jörn Diedrichsen, Frank van Overwalle, Josh McDermott, Caroline Nettekoven, Elizabeth Lee, Emalie McMahon, members of the Kanwisher Lab, and the audiences at the Neurobiology of Language conference (2023, Marseille), the Cognitive Neuroscience Society meeting (2025, Boston), and the Cerebellum Gordon Research Conference (2025, Les Diablerets) for helpful discussions and comments on the analyses and manuscript. CC was supported by the Kempner Institute for the Study of Natural and Artificial Intelligence at Harvard University. GT was supported by The K. Lisa Yang ICoN Center Graduate Fellowship and MIT’s McGovern Institute for Brain Research. HS was supported by NSF GRFP DGE2139757. BL was supported by NSF GRFP DGE2141064. AD was supported by the Simons Foundation Autism Research Initiative (SFARI) Bridge to Independence Award. EF was partially supported by NIH awards R01-DC016607, R01-DC016950, and U01-NS121471, as well as by funds from MIT’s McGovern Institute for Brain Research, Quest for Intelligence, Department of Brain and Cognitive Sciences, and the Simons Center for the Social Brain.

## Author Contribution Statement

Conceptualization: CC and EF; Methodology: CC, MP, GT, HS, AW, BL, AD, EF; Software, Validation: CC, MP, GT, HS, AW; Formal analysis: CC, MP, GT, AW; Investigation: all authors; Data curation: CC, GT, HS, PS, AW, BL, EF; Writing-original draft: CC; Writing-review and editing: EF and AD, with additional input from all authors; Visualization: CC; Supervision: EF, AD; Project administration: EF.

## Supplementary

**Supplementary Figure 1:**
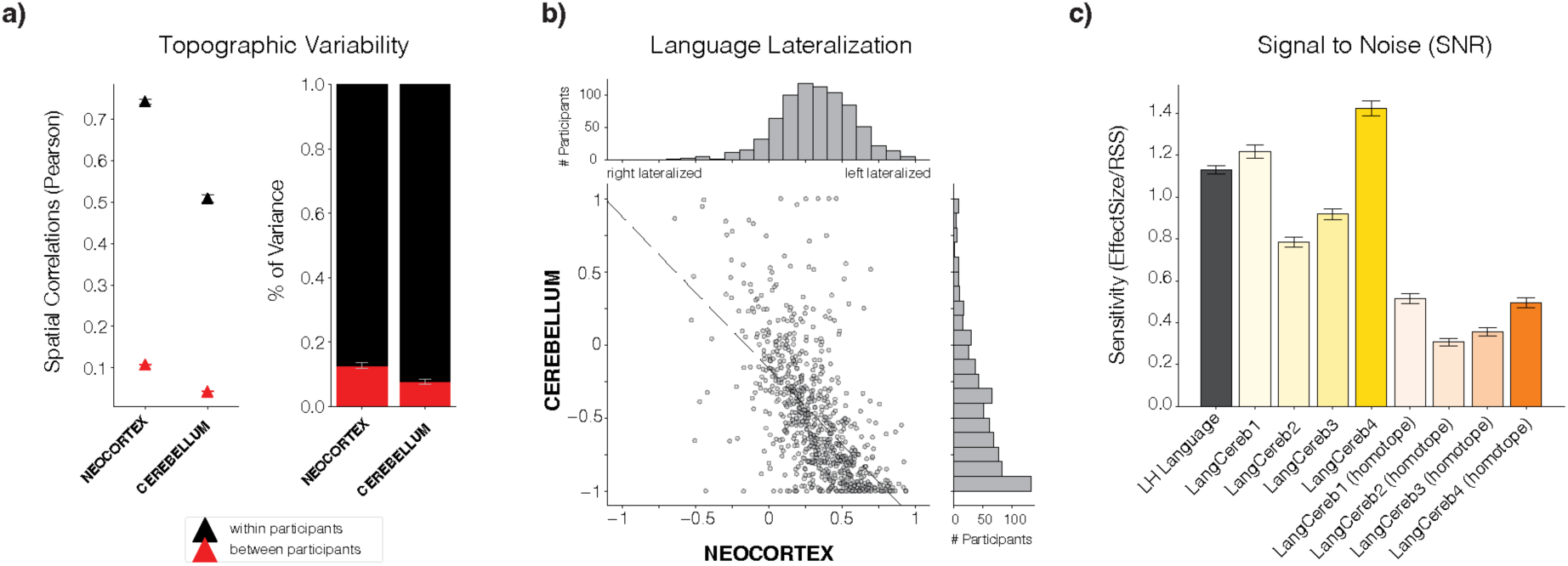
Inter-participant topographic variability, language lateralization, and signal quality. **a)** Spatial correlations of language responses (Language-Control contrast, **Fig. 1a**) within-(black) vs. between-participants (red) in the cerebellum and the neocortex. Within-participant spatial correlations were computed between odd and even runs of the language localizer. Between-participant spatial correlations were computed as the average of the spatial correlations between a given participant and all possible run/participant pairings. We then normalized the between-participant spatial correlations by the sum of the between- and within-participant correlations. We found that on average 13.7% of the pattern variance was shared across individuals in the neocortical language network whereas only 8.6% of the pattern variance was shared across individuals in the cerebellar language regions, indicating that language responses in the cerebellum are more topographically variable than in the neocortex (p<0.001, Wilcoxon signed-rank test). Error bars reflect standard error of the mean over participants. **b)** Lateralization of language responses within individual participants in the cerebellum vs. the neocortex. Lateralization was computed as the difference between the number of significant language-responsive voxels (uncorrected t > 2.576, p<0.01, two-sided t-test) in the left {cerebellum, neocortex} minus the number of significant voxels in the right {cerebellum, neocortex}, over the sum of the total number of significant voxels in the {cerebellum, neocortex}. **c)** A measure of the signal-to-noise ratio in the neocortical language network and the cerebellar language regions. This value was calculated for all participants in Expt. 1a (n=754) by dividing the response in a given region to the Sentence condition from the language localizer by the residual sum-of-squares (RSS) from the first-level GLM (averaged over all voxels in the fROI). This value represents the ratio of the task-relevant “signal” to the task-independent “noise”. This analysis indicates that the SNR is significantly worse in the LangCereb3 than in the neocortical language network (p<0.001, Wilcoxon signed-rank test). Error bars reflect standard error of the mean over participants.

**Supplementary Figure 2:**
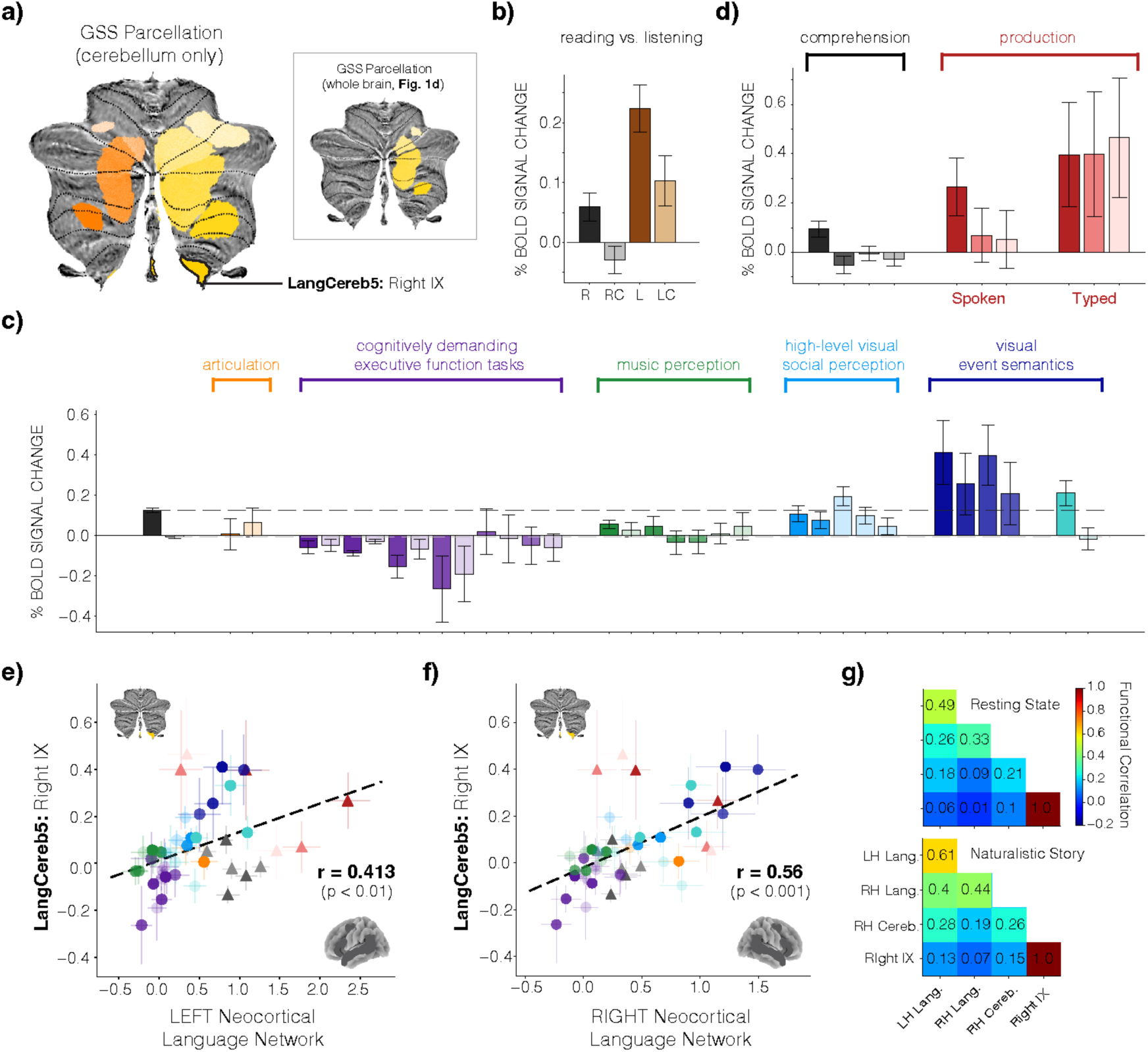
Responses in LangCereb5 located in right lobule IX. **a)** GSS analysis applied exclusively to the cerebellum using the top 20% (cf. 10%) of language-responsive voxels. The analysis reveals an additional cerebellar language region (LangCereb5) located in right lobule IX. **b)** Responses to reading (black) and listening (brown) in LangCereb5 (IX; as in **Fig. 2c-g**). **c)** Responses to the suite of non-linguistic inputs and tasks (Expts. 2a-e) in LangCereb5 (IX; as in **Fig. 3**). LangCereb5 responds more strongly to visual semantic information (e.g., Expts. 2e-i and 2e-ii, blue bars) than to language. The identify of the bars follows **Fig. 3**. Error bars reflect the standard error of the mean over participants (see **Fig. 3** and <u>Methods</u> for the number of participants per experiment). **d)** Responses to Expts. 3a-b in LangCereb5 (IX; as in Fig. 4). Error bars reflect the standard error of the mean over participants (see **Fig. 4** and <u>Methods</u> for the number of participants per experiment). **e)** Comparing the response profile of LangCereb5 (IX) with the left neocortical language network (the average of the five ‘core’ regions, <u>Methods</u>). Individual points correspond to experimental conditions from **c-d** (excluding the Sentence and Nonword list comprehension conditions, as in **Fig. 6**, <u>Methods</u>). Error bars on individual points reflect the standard error of the mean over participants for the given experiment. The line of best fit is plotted in black along with the Pearson correlation between the response profiles of LangCereb5 and the neocortical language network**. f)** Comparing the response profile of LangCereb5 (IX) with the right neocortical language network (the average response of the homotopes of the five ‘core’ left hemisphere regions). **g)** Functional correlations during rest (top) and naturalistic story listening (bottom) between LangCereb5 and the left neocortical language network, the right neocortical language network, and the other right cerebellar language regions (LangCereb1-4; functional correlations calculated as in **Fig. 6**).

**Supplementary Figure 3:**
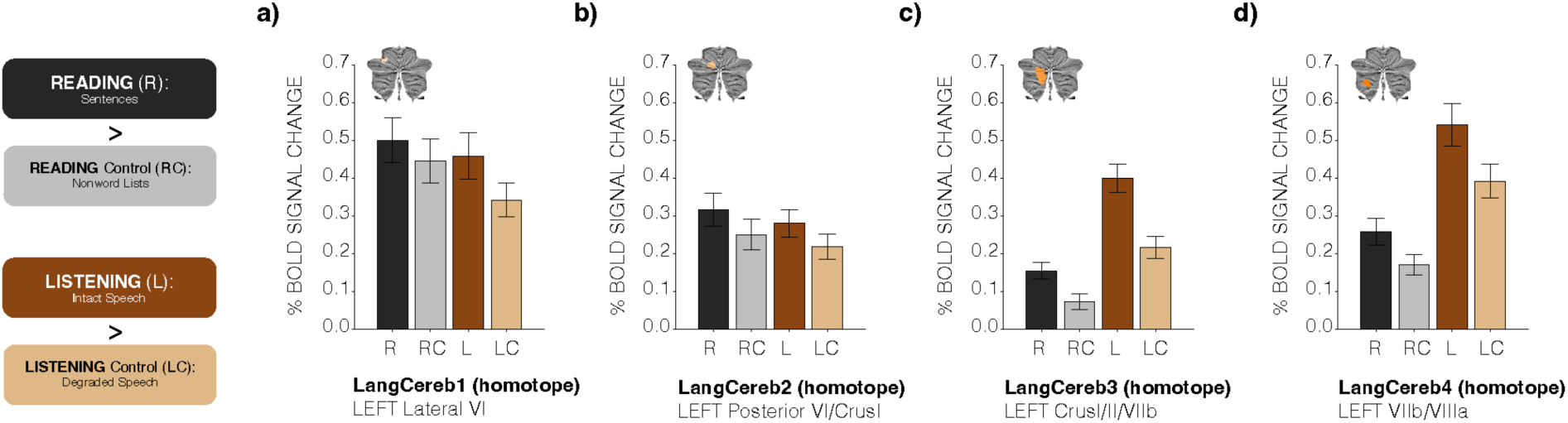
Engagement of the left cerebellar homotopes during reading and listening. a-d) Responses during reading and listening in the four, left-cerebellar homotopes. The left cerebellar homotopes respond less consistently across reading and listening. Responses reflect the percent BOLD signal change in the individually-defined language fROIs (Methods), and error bars reflect the standard error of the mean over participants.

**Supplementary Figure 4:**
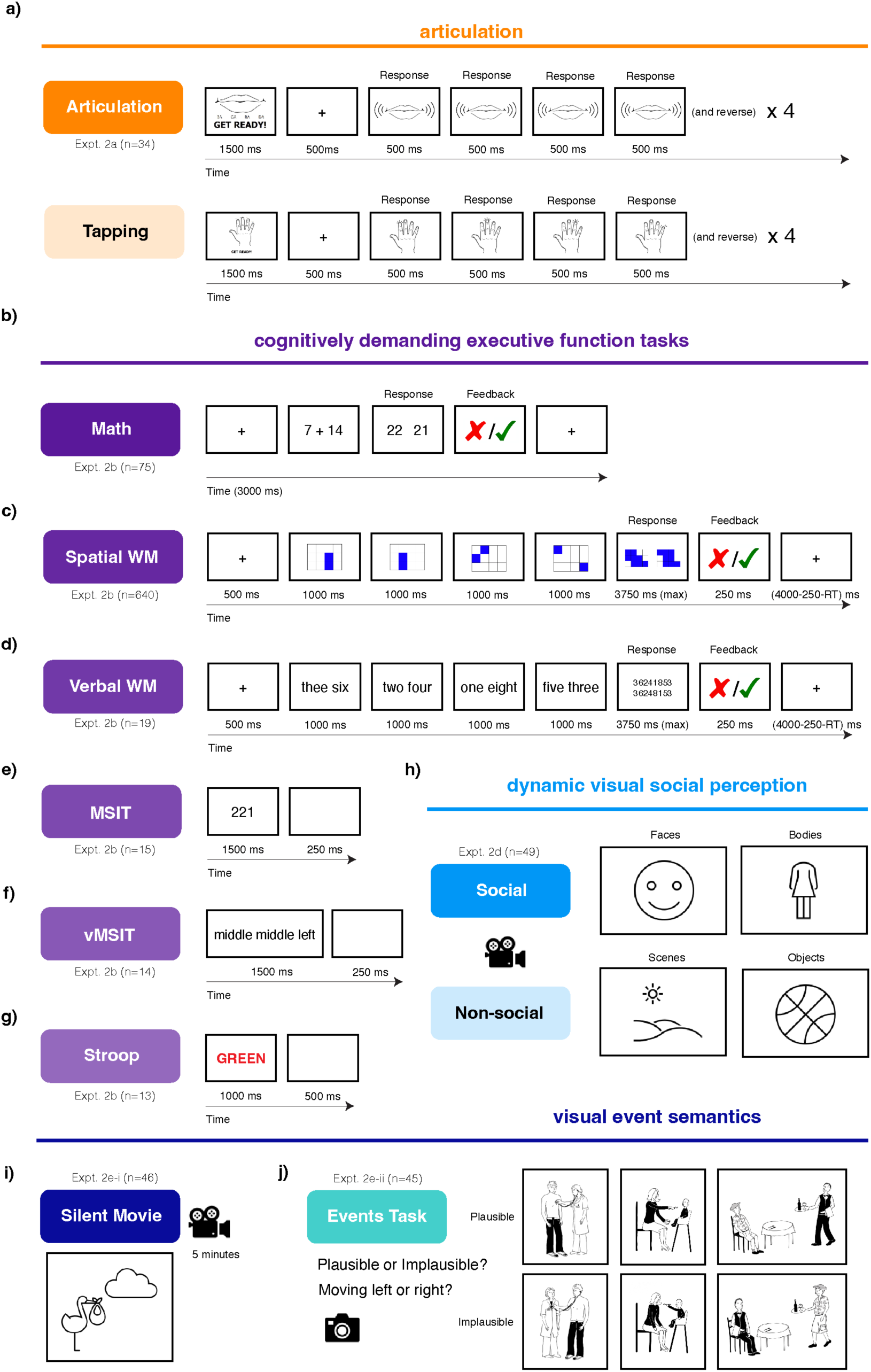
Example items from the suite of non-linguistic conditions. a) Expt. 2a (n=33), articulation task (data from Wolna et al., in prep.). Participants were instructed to overtly recite a syllable sequence “ba-da-ga-ra” (top) or tap a sequence using their fingers (bottom). **b)** Expt. 2b-I (n=75) arithmetic task (data from Malik-Moraleda, Ayyash et al., 2022). **c)** Expt. 2b-ii (n=640) spatial working memory task (data from Fedorenko et al., 2011; Lipkin et al., 2023). **d)** Expt. 2b-iii (n=19) verbal working memory task (data from Fedorenko et al., 2011 for this and all subsequent executive function tasks). **e)** Expt. 2b-iv (n=15) multi-source interference task. **f)** Expt. 2b-v (n=14) verbal multi-source interference task. **g)** Expt. 2b-vi (n=13) Stroop task. **h)** Expt. 2d (n=49) dynamic visual social perception task (data from Pritchett et al., 2018). Participants viewed videos of faces and bodies (both social) or scenes and objects (both non-social). **i)** Expt. 2e-i (n=46) silent movie viewing (Pixar’s *Partly Cloudy;* data from Shain, Paunov, Chen et al., 2023). **j)** Expt. 2e-ii (n=45) event semantics task (data from Ivanova et al., 2021; Ivanova, 2022). Participants were either instructed to evaluate the semantic plausibility of the event in the image or determine whether the image was moving to the left or right. See Methods for additional experimental details.

**Supplementary Figure 5:**
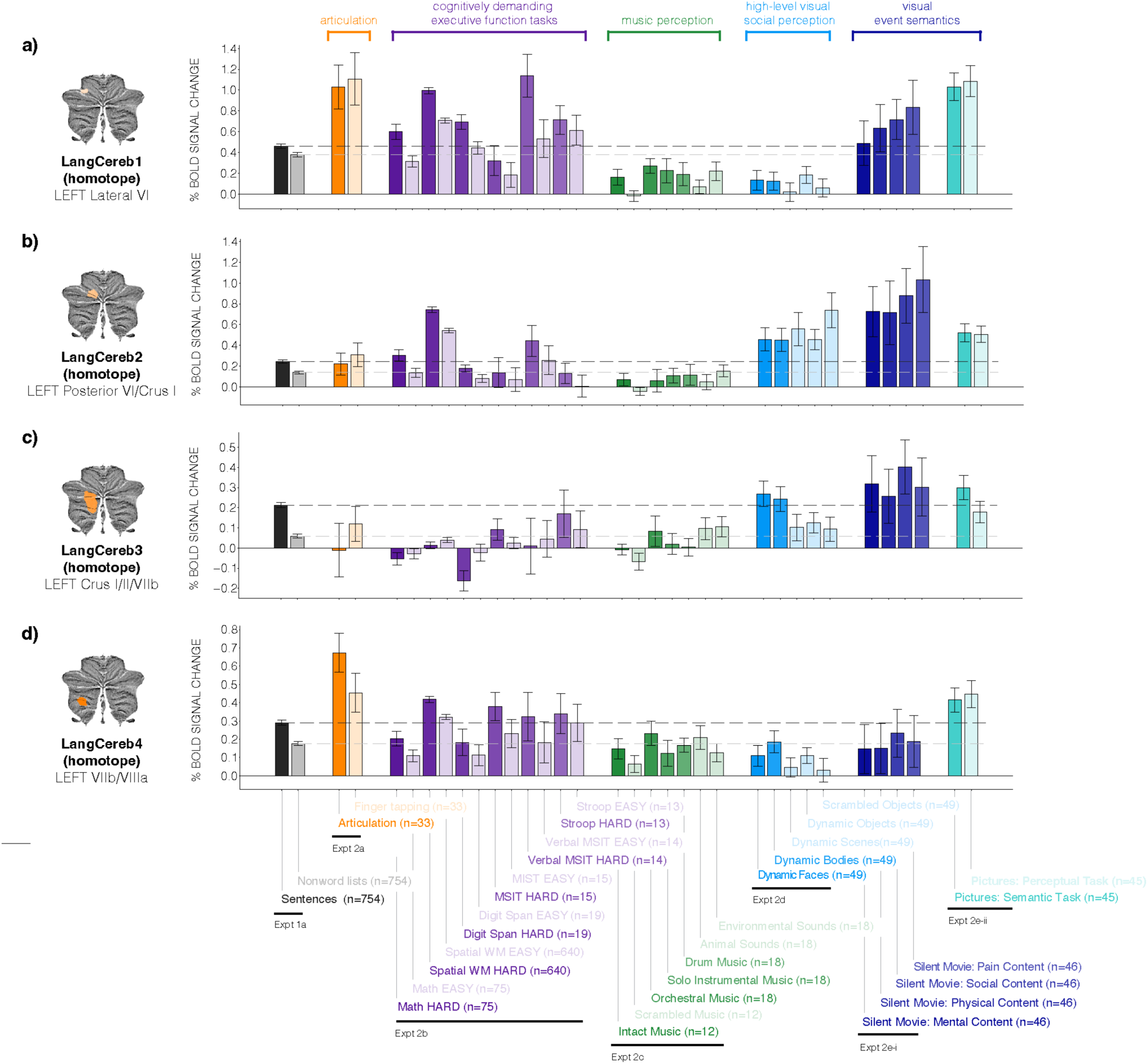
Selectivity of left cerebellar language regions for language over non-linguistic inputs and tasks. a-d) Responses in the left cerebellar homotopic regions of LangCereb1-4 to the suite of non-linguistic tasks (Expts. 2a-e; as in **Fig. 3**). Across all experiments, control conditions are denoted by lightly shaded bars. Responses reflect the percent BOLD signal change in the individually-defined language fROIs (<u>Methods</u>). All error bars reflect the standard error of the mean over participants for the given condition. The dashed black line indicates the response of a given region to language; the dashed gray line indicates the response of a given region to the control condition from the language localizer.

**Supplementary Figure 6:**
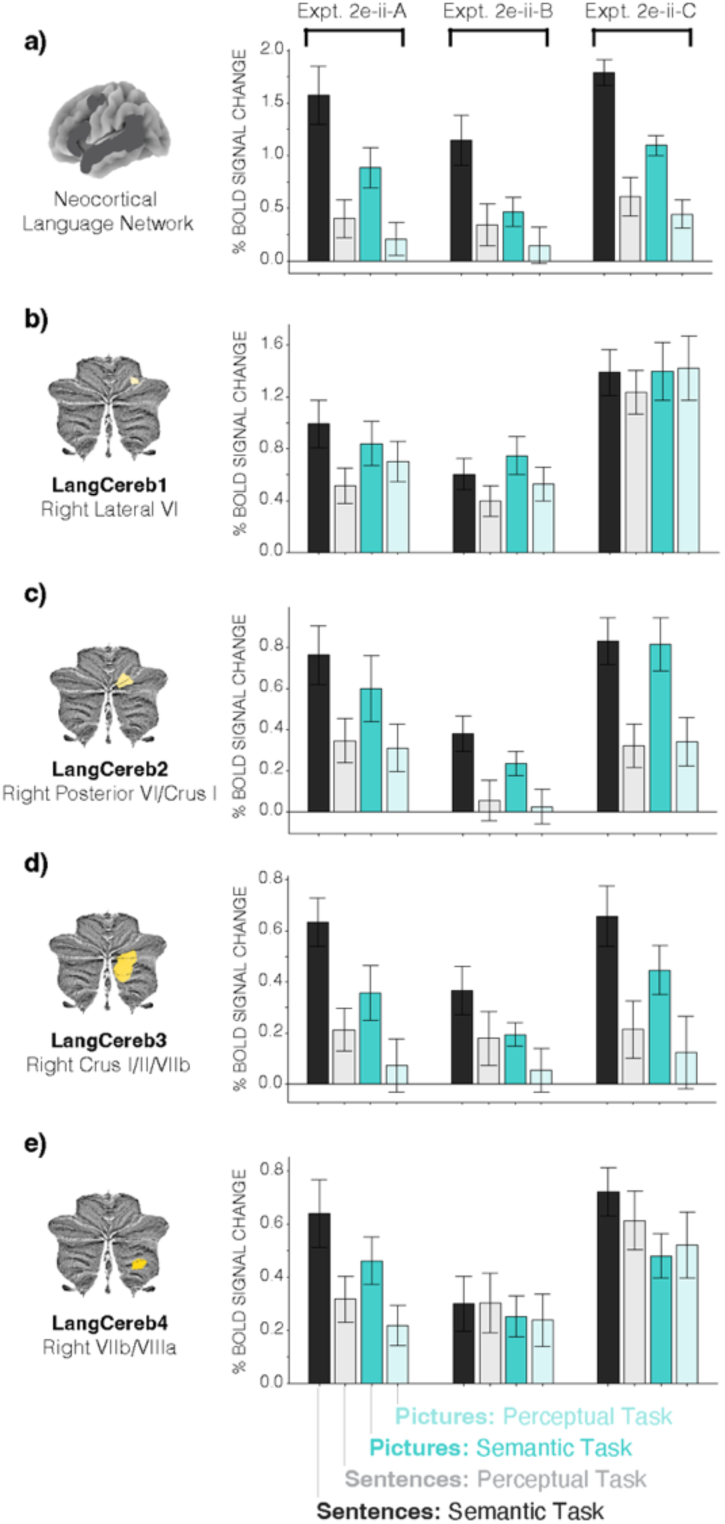
Responses in the neocortical language network and LangCereb1-4 to all event semantics experiments visualized separately (Expt. 2e). a-e) Responses to the three events semantics experiments (Expts. 2e-ii-A, B, C) in the neocortical language network and LangCereb1-4 visualized separately. In Expt. 2e-ii-A, participants were shown pictures and sentences of agent-patient interactions that were either plausible or implausible. In Expt. 2e-ii-B, participants were shown pictures and sentences of people interacting with everyday objects that were either plausible or implausible. In Expt. 2e-ii-C, participants were shown pictures and sentences of people performing actions on everyday objects, with the actions being reversible or irreversible (see Methods for details). In addition to the two non-linguistic (visual) conditions (teal, included in Fig. 3), responses to linguistic conditions are provided as well (black). Participants were shown a sentence and either asked if the event was plausible or implausible (‘Semantic Task’; for Expt. 2e-ii-C participants were asked if the action was reversable or irreversible) or asked if the sentence was moving slightly to the left or the right (‘Perceptual Task’). Responses reflect the percent BOLD signal change in the individually-defined language fROIs (Methods). All error bars reflect the standard error of the mean over participants for the given condition. The neocortical language network responds to the visual semantic task, but it has a preference for semantic content that is delivered linguistically. LangCereb3 (Crus I/II/VIIb) shows a similar pattern of response. LangCereb2 (posterior VI/Crus I) responds similarly to visual and linguistic semantic content. LangCereb1 (lateral VI) and 4 (VIIb/VIIIa) respond as strongly to the perceptual task as they do to the semantic task for multiple experiments (in both modalities) suggesting they respond to generally engaging tasks.

**Supplementary Figure 7:**
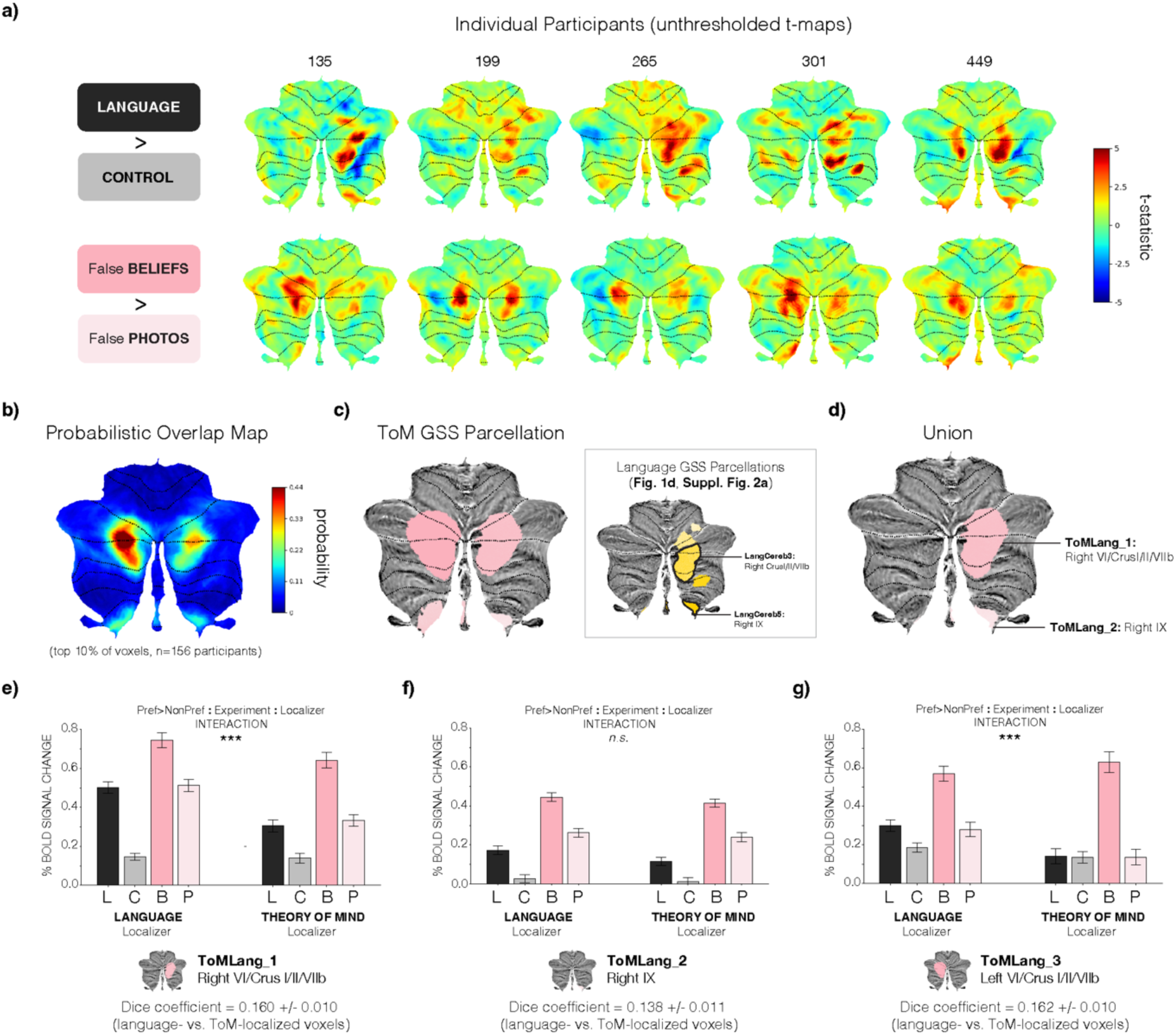
Dissociation of language and Theory of Mind networks in the cerebellum. **a)** Cerebellar responses to language (Language-Control, top) and Theory of Mind (ToM, Expt. 2f, n=156, Saxe & Kanwisher, 2003; False Beliefs-False Photos, bottom) in 5 individual participants, visualized on a flattened cerebellar cortical surface. Responses reflect an unthresholded t-statistic and cerebellar lobule boundaries are marked in black. In the Theory of Mind task, participants were presented with written vignettes that describe a situation where an agent holds a false belief about the world, and vignettes that describe a situation where a visual depiction of the word became outdated (see <u>Methods</u> for details). **b)** Probabilistic overlap map across n=156 participants constructed from individual binary masks of ToM-responsive voxels, visualized on a flattened cerebellar cortical surface (as in **Fig. 1c**). Individual binary masks took a value of 1 for the top 10% of ToM-responsive voxels across the whole brain and 0 otherwise. Color shading of the overlap map reflects the percentage of participants for whom that area was in their top 10% of most ToM-responsive voxels. **c)** Parcellation resulting from the group-constrained subject-specific (GSS) analysis which defines common areas of activation across a group of participants (<u>Methods</u>). Four functional regions of interest (fROIs) were identified. *Inset:* Language parcels from **Fig. 1d** and **Suppl. Fig. 2a** with the regions that most overlap with the ToM parcels marked in black (LangCereb3, right Crus I/II/VIIb; LangCereb5, right IX). **d)** Union of the ToM parcels and the language parcels highlighted in **c**. **e-g**) Responses to the language (black) and ToM (pink) localizers in the most language-(left) or ToM-responsive (right) voxels in a given region. When measuring responses to the same task that was used to identify the voxels, separate runs of the data were used. If language and ToM networks are dissociable within a given region, then we expect to observe a ‘Preference>Non-Preference’ by ‘Experiment’ by ‘Localizer’ interaction. In particular, the difference between the preferred (Language or False Beliefs) and non-preferred (Control or False Photos) conditions should differ by the localizer that was used (evaluated with a linear mixed-effects model, <u>Methods</u>). Language and ToM voxels are dissociable within ToM+Lang_1 (right VI/Crus I/II/VIIb; p<0.001; we note that the remaining B>P response could be due to a known linguistic confound in the ToM stimuli, see Shain, Paunov, Chen et al., 2023). Language and ToM voxels are not dissociable in ToM+Lang_2 (right IX). Finally, language and ToM voxels are dissociable in ToM+Lang_3 (left VI/Crus I/II/VIIb, defined as the union between the LangCereb1 left homotope and the ToM parcels from **c**). However, importantly, this region shows stronger ToM responses than ToM+Lang_1, as the language-localized voxels exhibit a larger B-P difference, and conversely, the ToM-localized voxels do not exhibit a language response (L-C is approximately 0). We also provide the average Dice coefficient of language- and ToM-localized voxels across participants, as a measure of how overlapping the fROIs are within a given parcel. The provided error values reflect standard error of the mean over participants.

**Supplementary Figure 8:**
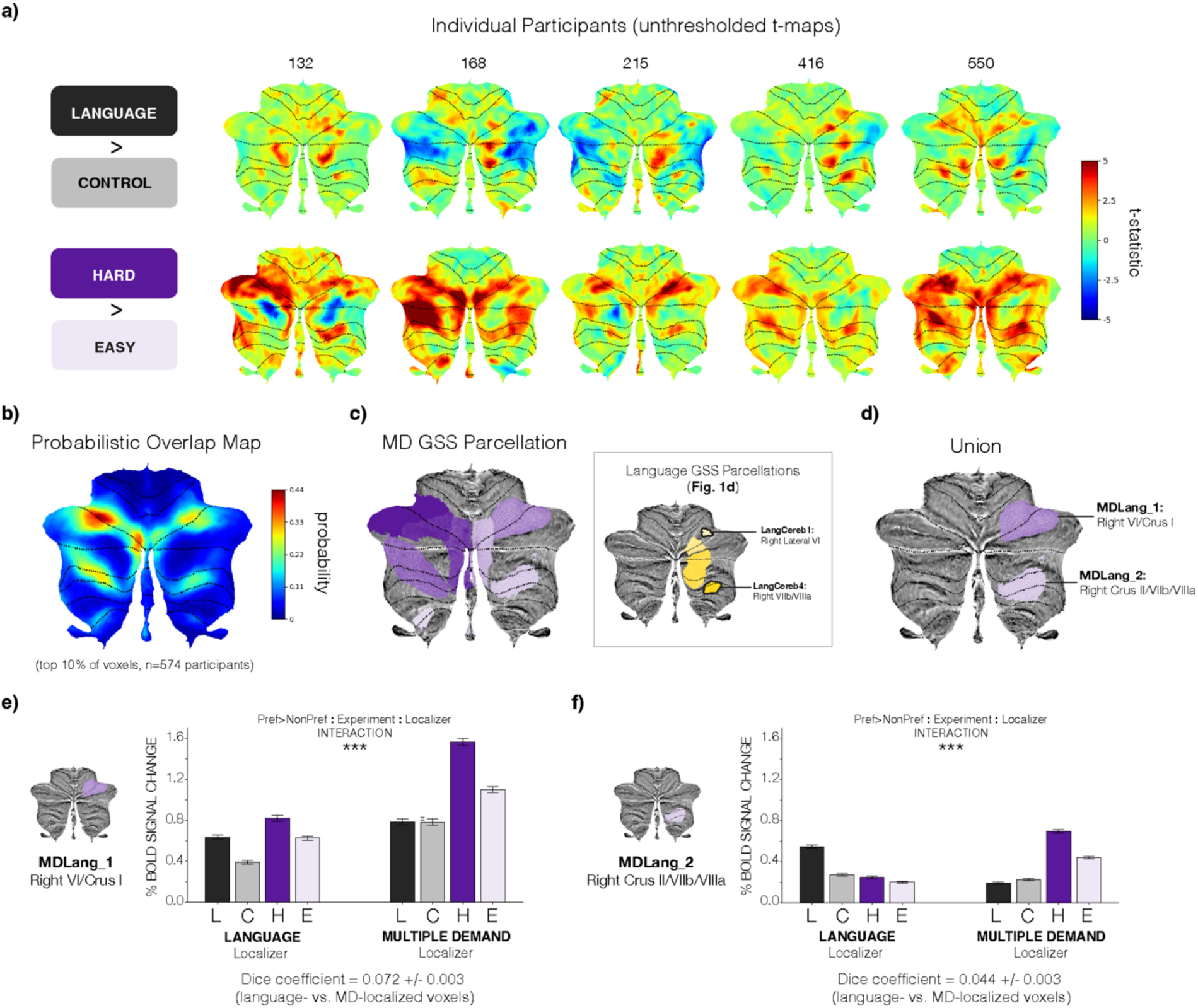
Dissociation of language and Multiple Demand networks in the cerebellum. **a)** Cerebellar responses to language (Language-Control, top) and a spatial working memory task commonly used to localize the ‘Multiple Demand’ (MD) network (Hard-Easy, Expt. 2b-ii, bottom) in 5 individual participants, visualized on a flattened cerebellar cortical surface. Responses reflect an unthresholded t-statistic and cerebellar lobule boundaries are marked in black. In the spatial working memory task, participants were presented with a 3×4 grid and asked to keep track of where blue squares appeared. There was a hard and an easy condition (see <u>Methods</u> for details). **b)** Probabilistic overlap map across n=574 participants constructed from individual binary masks of MD-responsive voxels, visualized on a flattened cerebellar cortical surface. Individual binary masks took a value of 1 for the top 10% of MD-responsive voxels across the whole brain and 0 otherwise. **c)** Parcellation resulting from the group-constrained subject-specific (GSS) analysis which defines common areas of activation across a group of participants (<u>Methods</u>). Eight functional regions of interest (fROIs) were identified. *Inset:* Language parcels from **Fig. 1d** with the regions that most overlap with the MD parcels marked in black (LangCereb1, right lateral VI; LangCereb4, right VIIb/VIIIa). **d)** Union of the MD parcels and the language parcels highlighted in **c**. **e-f**) Responses to the language (black) and MD (purple) localizers in the most language-(left) or MD-responsive (right) voxels in a given region. When measuring responses to the same task that was used to identify the voxels, separate runs of the data were used. As in **Suppl. Fig. 6**, if language and MD networks are dissociable within a given region, then we expect to observe a ‘Preference>Non-Preference’ by ‘Experiment’ by ‘Localizer’ interaction (evaluated with a linear mixed-effects model, <u>Methods</u>). Language and MD voxels are dissociable within MD+Lang_1 (right VI/Crus I; p<0.001) and MD+Lang_2 (right Crus II/VIIb/VIIIa). We also provide the average Dice coefficient of language- and MD-localized voxels across participants, as a measure of how overlapping the fROIs are within a given parcel. The provided error values reflect standard error of the mean over participants.

**Supplementary Figure 9:**
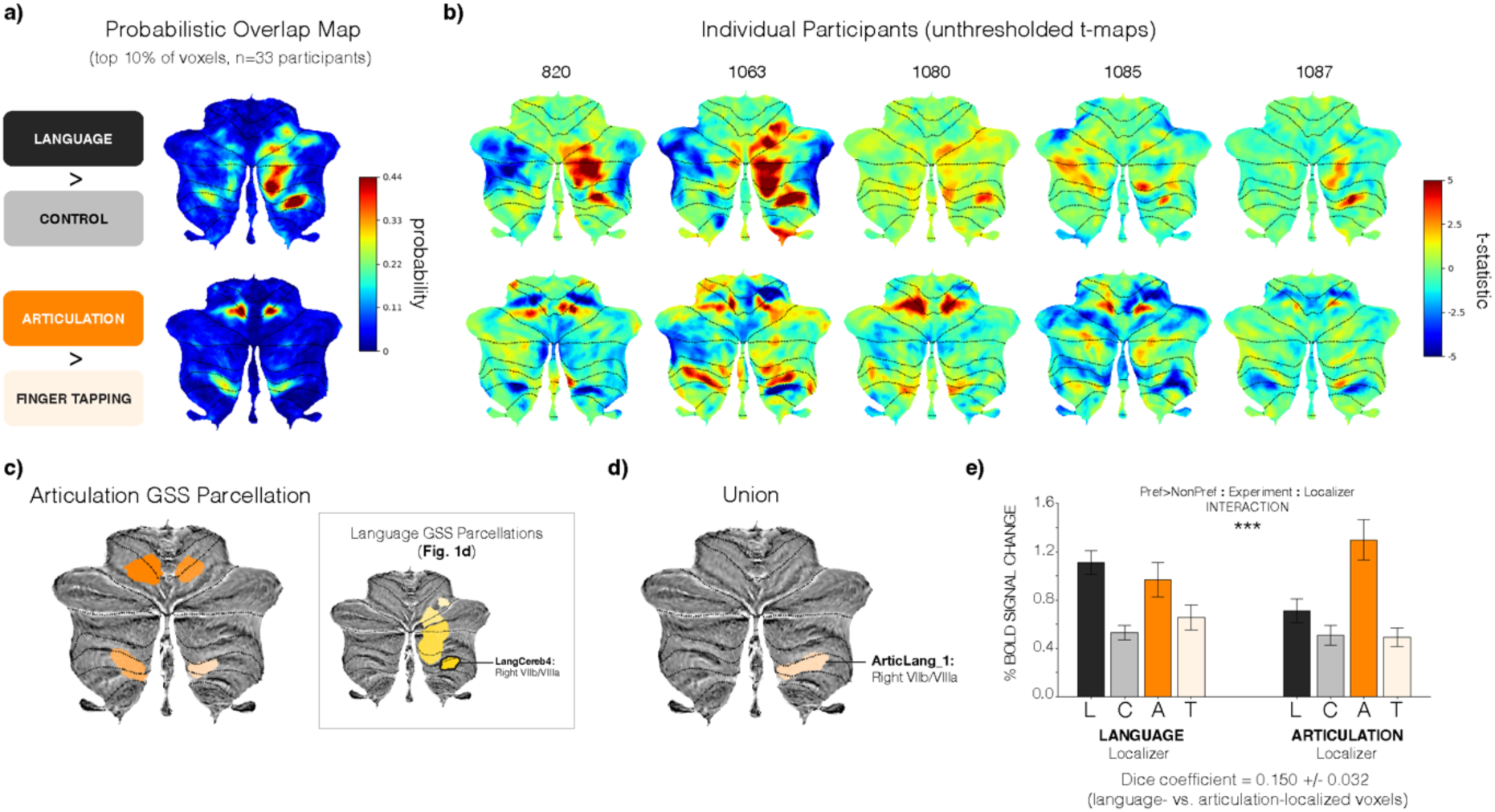
Dissociation of language and articulation networks in the cerebellum. **a)** Probabilistic overlap map across n=33 participants of language-responsive voxels (Language-Control, top) and articulation-responsive voxels (Articulation-Finger Tapping, bottom) constructed from individual binary masks, visualized on a flattened cerebellar cortical surface. Individual binary masks took a value of 1 for the top 10% of language- or articulation-responsive voxels across the whole brain and 0 otherwise. In the articulation task participants were instructed to repeat a sequence of 4 syllables (“ba, ga, ra, da”) or a simple finger-tapping motor sequence (see <u>Methods</u> for details). **b)** Cerebellar responses to language (Language-Control, top) and articulation (Articulation-Finger Tapping, Expt. 2b-ii, bottom) in 5 individual participants, visualized on a flattened cerebellar cortical surface. Responses reflect an unthresholded t-statistic and cerebellar lobule boundaries are marked in black. **c)** Parcellation resulting from the group-constrained subject-specific (GSS) analysis which defines common areas of activation across a group of participants (<u>Methods</u>). Four functional regions of interest (fROIs) were identified. *Inset:* Language parcels from **Fig. 1d** with the region that most overlaps with the articulation parcels marked in black (LangCereb4, right VIIb/VIIIa). **d)** Union of the articulation parcels and the language parcel highlighted in **c**. **e**) Responses to the language (black) and articulation (orange) localizers in the most language-(left) or articulation-responsive (right) voxels Artic+Lang_1. When measuring responses to the same task that was used to identify the voxels, separate runs of the data were used. As in **Suppl. Figs. 6-7**, if language and articulation networks are dissociable within a given region, then we expect to observe a ‘Preference>Non-Preference’ by ‘Experiment’ by ‘Localizer’ interaction (evaluated with a linear mixed-effects model, <u>Methods</u>). Language and articulation voxels are dissociable within Artic+Lang_1 (right VIIb/VIIIa; p<0.001). We also provide the average Dice coefficient of language- and articulation-localized voxels across participants, as a measure of how overlapping the fROIs are within the union parcel. The provided error values reflect standard error of the mean over participants.

**Supplementary Figure 10:**
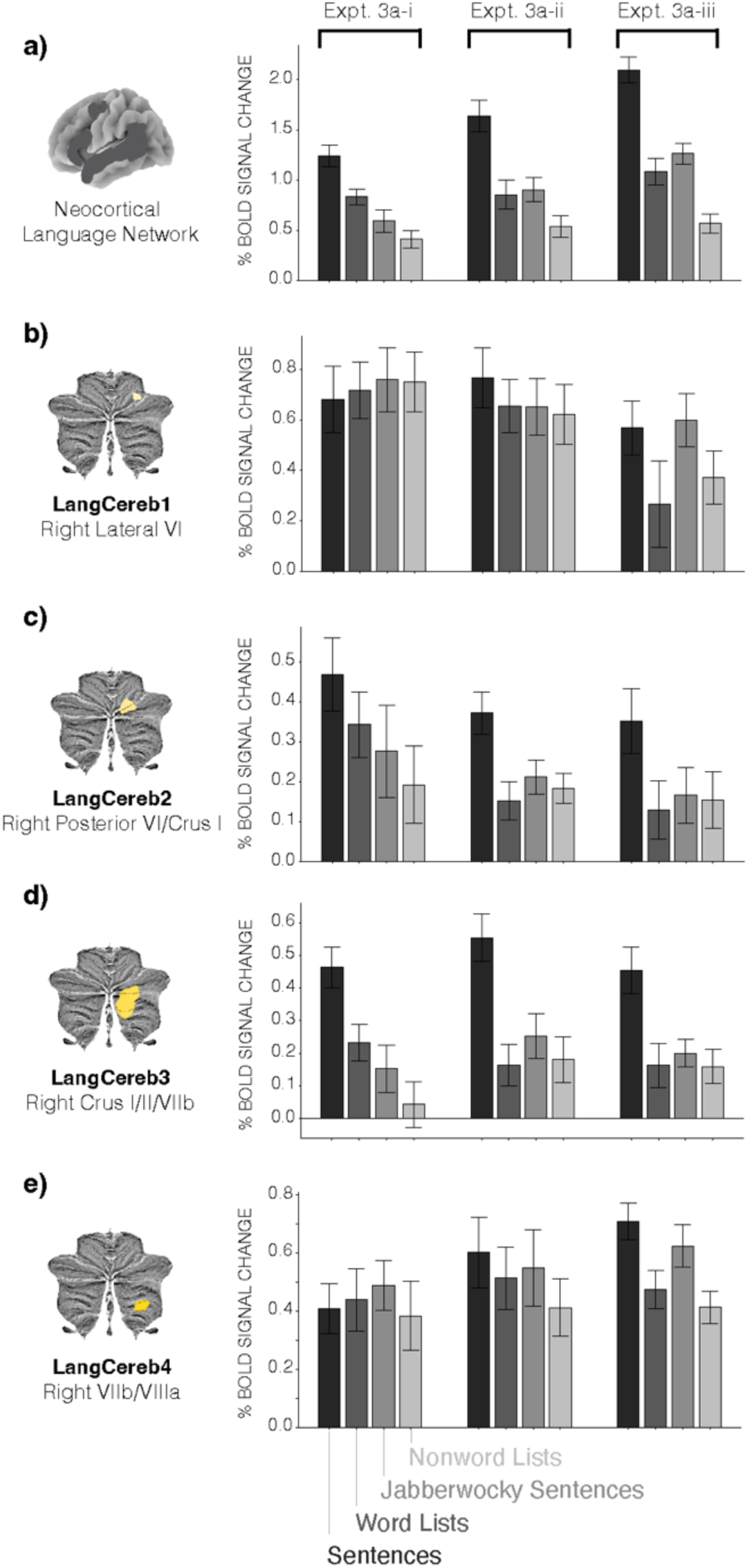
Responses in LangCereb1-4 to all language comprehension experiments. a-d) Responses to the three language comprehension experiments (Expts. 3e-i, ii, and iii) in LangCereb1-4 visualized separately. In all three experiments, participants passively read sentences (S), word lists (W), Jabberwocky sentences (J; sentences where content words are replaced by nonwords, such as “florped” or “blay”), and nonword lists (N). The experiments followed a 2×2 design, manipulating lexical access and syntactic structure building (see <u>Methods</u> for details). Responses reflect the percent BOLD signal change in the individually-defined language fROIs (<u>Methods</u>). All error bars reflect the standard error of the mean over participants for the given condition.

**Supplementary Figure 11:**
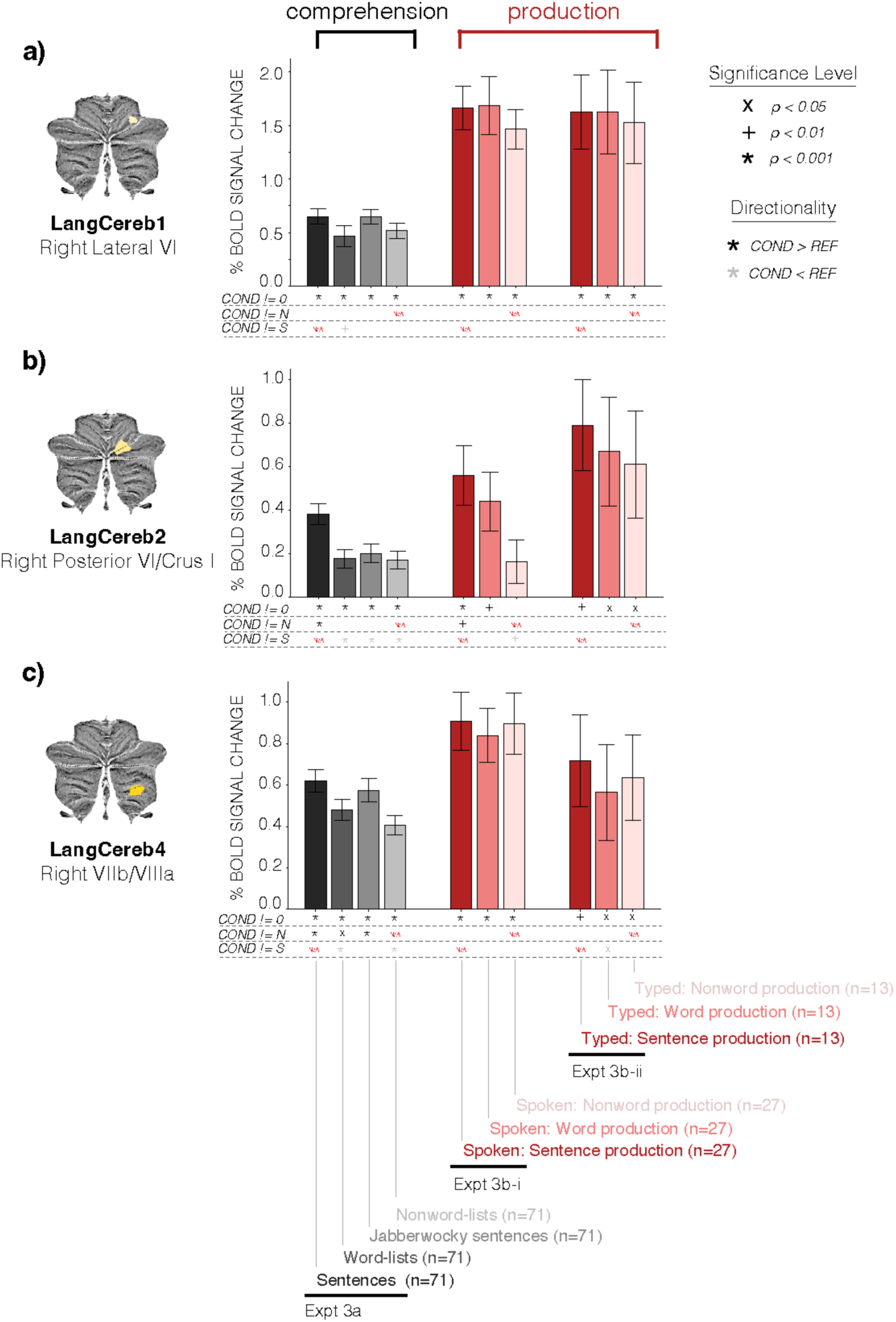
Linguistic contributions of the non-selective cerebellar language regions during comprehension and production. **a-c)** Responses in LangCereb1 (right lateral IV, **a**), LangCereb2 (right posterior VI/Crus I, **b**), and LangCereb4 (right VIIb/VIIIa, **c**) to the language comprehension experiment (Expt. 3a, n=71, data from Fedorenko et al., 2011; Shain, Kean et al., 2024; and Kauf et al., 2024) and the language production experiment (Expt. 3b, n=40, data from Hu, Small et al., 2023), as in **Fig. 4**. Responses reflect the percent BOLD signal change in the individually-defined language fROIs (<u>Methods</u>). All error bars reflect the standard error of the mean over participants for the given condition. Responses were statistically evaluated relative to i) a fixation baseline, ii) the response to nonword lists (in the relevant experiment), and iii) the response to sentences (in the relevant experiment) using linear mixed-effects models (see **Suppl. Tables 29-30**, **32**, <u>Methods</u>).

**Supplementary Figure 12:**
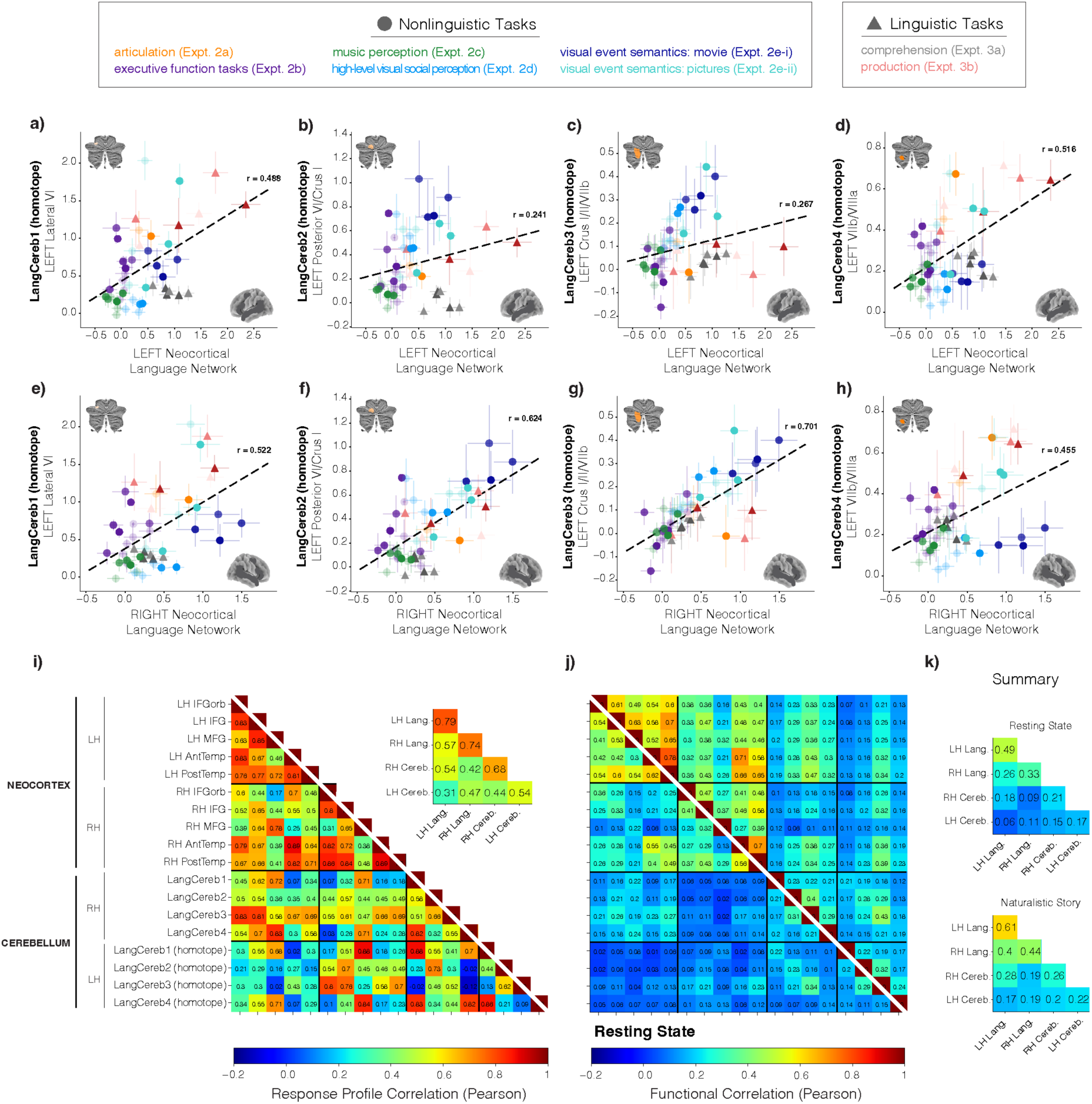
Response profile similarity and functional correlations of the left cerebellar language regions to the neocortical language regions. **a-h)** Correlation in the response profiles (i.e., the magnitudes of response from Expts. 2a-e and 3a-b, **Fig. 3** and **4**) of the left cerebellar language regions vs. the left-hemisphere neocortical language network (**a-d**) and the right-hemisphere neocortical language regions (**e-h**), as in **Fig. 6**. Responses to individual conditions are shown in the same colors as in **Fig. 3** and **4**. Error bars around individual points reflect the standard error of the mean over participants for that condition. Black dashed lines correspond to the line of best fit. **i)** Correlation in the response profiles between the left-hemisphere cerebellar language regions and individual neocortical regions. *Inset:* Summary of response profile correlations by gross anatomical region (left-hemisphere neocortex, right-hemisphere neocortex, and left cerebellum). **j)** Functional correlations between the left-hemisphere cerebellar and neocortical language regions during resting state (lower triangle) or naturalistic story listening (upper triangle; Expt. 4a-b, n=85, data from Malik-Moraleda, Ayyash et al., 2022; <u>Methods</u>). **k)** Summary of functional correlations by gross anatomical region.

**Supplementary Figure 13:**
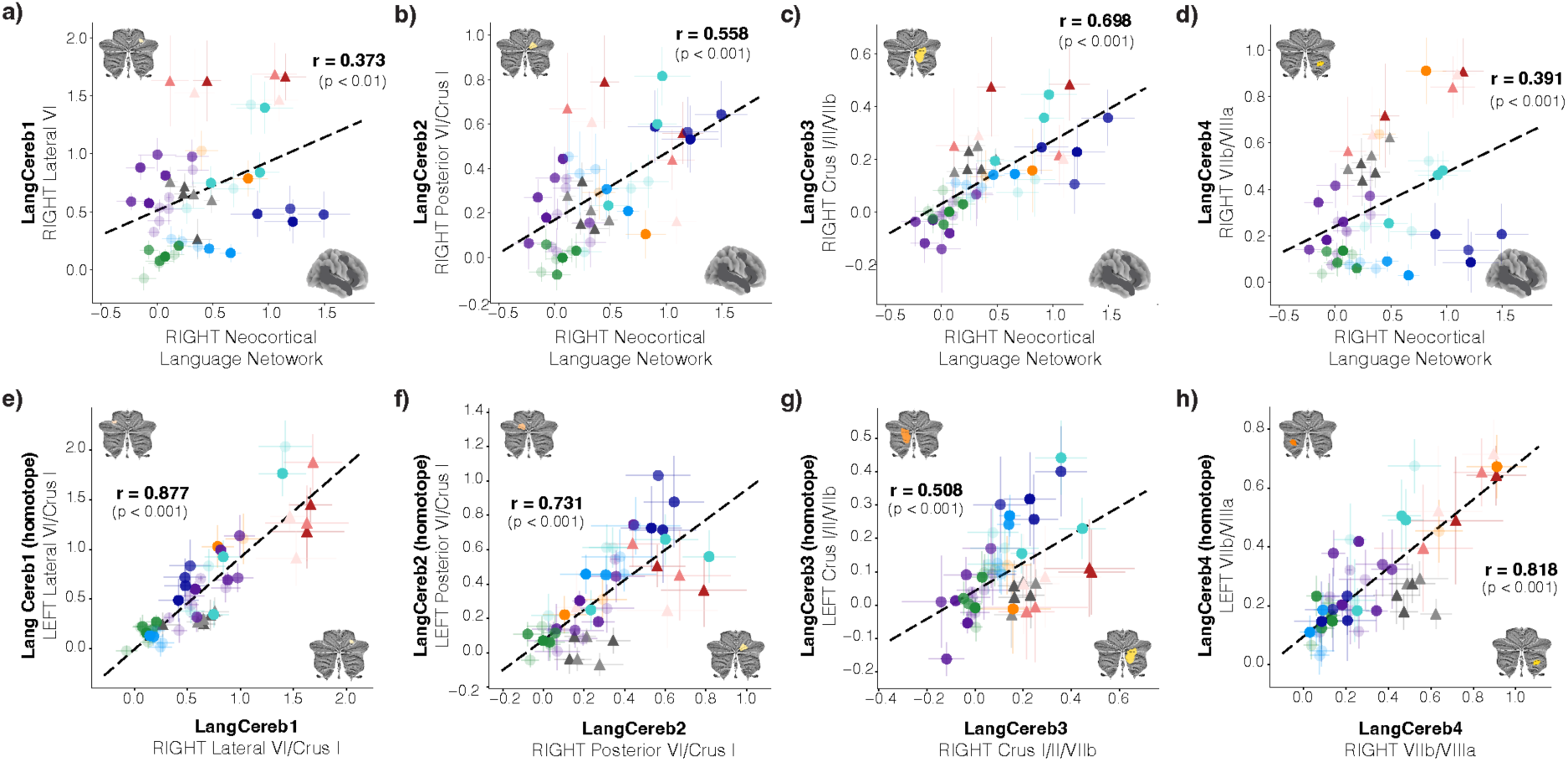
Response profile similarity of the right cerebellar language regions to their left cerebellar homotopes, and to the right neocortical language regions. a-h) Correlation in the response profiles (i.e., the magnitudes of response from Expts. 2a-e and 3a-b, Fig. 3 and 4) of the right-hemisphere cerebellar language regions vs. the right-hemisphere neocortical language regions (a-d) and vs. the corresponding left cerebellar homotopic area (e-h). Responses to individual conditions are shown in the same colors as in Fig. 3 and 4. Error bars around individual points reflect the standard error of the mean over participants for that condition. Black dashed lines correspond to the line of best fit.

---

**Supplemental Table 1:** Language (L) vs. a perceptually-matched control condition (C) in the cerebellar language regions. Linear mixed-effects (LME) models were used to test whether the cerebellar language regions responded more strongly to language than to a control condition (nonword lists; **Fig. 1g**). A separate model was fit per cerebellar language region. ‘EffectS’ was a binary predictor (1 for the Language condition, 0 for the Control condition). The LME models were fit with maximum likelihood estimation and included ‘Participant’ as a random effect. Bolded terms are significant (p < 0.05).

|  | *ROI* | *Estimate* | *Std..Error* | *df* | *t.value* | *P.uncorrected* | *P.corrected* |
| --- | --- | --- | --- | --- | --- | --- | --- |
| **(Intercept)** | **Right_CrusI_II_VIIb** | **0.1059** | **0.010822** | **1111** | **9.7857** | **9.5172e-22** | **2.5866e-21** |
| **Effect** | **Right_CrusI_II_VIIb** | **0.36759** | **0.0097065** | **754** | **37.87** | **1.3054e-176** | **2.8382e-175** |
| **(Intercept)** | **Right_posterior_VI_CrusI** | **0.1582** | **0.013353** | **963.3** | **11.847** | **2.5199e-30** | **7.8272e-30** |
| **EffectS** | **Right_posterior_VI_CrusI** | **0.32208** | **0.0094051** | **754** | **34.245** | **9.2097e-156** | **1.0012e-154** |
| **(Intercept)** | **Right_VIIb_VIIIa** | **0.34127** | **0.01285** | **976.78** | **26.557** | **2.0456e-117** | **1.1119e-116** |
| **EffectS** | **Right_VIIb_VIIIa** | **0.29476** | **0.0093117** | **754** | **31.655** | **1.4645e-140** | **1.0614e-139** |
| **(Intercept)** | **Right_lateral_VI** | **0.54847** | **0.020464** | **884.58** | **26.802** | **2.5955e-116** | **1.1287e-115** |
| **EffectS** | **Right_lateral_VI** | **0.25646** | **0.011594** | **754** | **22.12** | **6.0403e-84** | **2.1889e-83** |

**Supplemental Table 2:** Overlap of cerebellar language regions with published cerebellar atlases. Dice coefficients comparing the cerebellar language regions reported in the present study with two published cerebellar atlases (Buckner et al., 2011; Nettekoven et al., 2024). Our symmetric cerebellar parcels (**Fig. 1d**, originally in SPM IXI549Space space) were first converted into SUIT space (Diedrichsen et al., 2009) prior to performing any comparisons with the existing atlases (both in SUIT space). *Top:* Overlap of the cerebellar language regions with the Buckner et al. (2011) 7-network parcellation. *Bottom:* Overlap with the Nettekoven et al. (2024) symmetrical 32-region parcellation. LangCereb3 (Crus I/II/VIIb) and LangCereb2 (posterior VI/Crus I) overlap substantially with Network7 (i.e., the Default Network) in Buckner et al. (2011) and the S1R and S2R (i.e., sociolinguistic) regions in Nettekoven et al. (2024). The other cerebellar language regions (LangCereb1 and 4) primarily overlap with the Network 4 and 6 regions from Buckner et al. (2011) and the D3R (i.e., demand) region from Nettekoven et al. (2024).

|  | LangCereb1 | LangCereb2 | LangCereb3 | LangCereb4 | LangCereb1<br>(homotope) | LangCereb2<br>(homotope) | LangCereb3<br>(homotope) | LangCereb4<br>(homotope) |
| --- | --- | --- | --- | --- | --- | --- | --- | --- |
| Network1 | 0.0 | 0.0 | 0.0 | 0.0 | 0.0 | 0.0 | 0.0 | 0.0 |
| Network2 | 0.0 | 0.0027 | 0.0 | 0.0541 | 0.0 | 0.0 | 0.0 | 0.0026 |
| Network3 | 0.0 | 0.0479 | 0.2191 | 0.2422 | 0.0 | 0.0139 | 0.2053 | 0.1707 |
| Network4 | 0.3158 | 0.0281 | 0.0 | 0.6158 | 0.4521 | 0.0733 | 0.0232 | 0.8165 |
| Network5 | 0.0 | 0.0 | 0.0016 | 0.0 | 0.0 | 0.0 | 0.0 | 0.0 |
| Network6 | 0.6842 | 0.2847 | 0.0766 | 0.0878 | 0.5479 | 0.7385 | 0.187 | 0.0103 |
| Network7 | 0.0 | 0.6366 | 0.7027 | 0.0 | 0.0 | 0.1744 | 0.5846 | 0.0 |
| M1L | 0.0 | 0.0 | 0.0 | 0.0 | 0.0 | 0.0 | 0.0 | 0.0 |
| M2L | 0.0 | 0.0 | 0.0 | 0.0 | 0.0 | 0.0381 | 0.0 | 0.0034 |
| M3L | 0.0 | 0.0 | 0.0 | 0.0 | 0.0 | 0.0 | 0.0 | 0.0986 |
| M4L | 0.0 | 0.0 | 0.0 | 0.0 | 0.0 | 0.0 | 0.0 | 0.0 |
| A1L | 0.0 | 0.0 | 0.0 | 0.0 | 0.0 | 0.0007 | 0.0539 | 0.0189 |
| A2L | 0.0 | 0.0 | 0.0 | 0.0 | 0.0 | 0.0 | 0.0 | 0.0652 |
| A3L | 0.0 | 0.0 | 0.0 | 0.0 | 0.0 | 0.0 | 0.0 | 0.0592 |
| D1L | 0.0 | 0.0 | 0.0 | 0.0 | 0.14 | 0.0784 | 0.0396 | 0.0352 |
| D2L | 0.0 | 0.0 | 0.0 | 0.0 | 0.0 | 0.0242 | 0.0401 | 0.0129 |
| D3L | 0.0 | 0.0 | 0.0 | 0.0 | 0.8575 | 0.0762 | 0.0682 | 0.7067 |
| D4L | 0.0 | 0.0 | 0.0 | 0.0 | 0.0025 | 0.0352 | 0.0137 | 0.0 |
| S1L | 0.0 | 0.0 | 0.0 | 0.0 | 0.0 | 0.707 | 0.4103 | 0.0 |
| S2L | 0.0 | 0.0 | 0.0 | 0.0 | 0.0 | 0.0403 | 0.3718 | 0.0 |
| S3L | 0.0 | 0.0 | 0.0 | 0.0 | 0.0 | 0.0 | 0.0 | 0.0 |
| S4L | 0.0 | 0.0 | 0.0 | 0.0 | 0.0 | 0.0 | 0.0024 | 0.0 |
| S5L | 0.0 | 0.0 | 0.0 | 0.0 | 0.0 | 0.0 | 0.0 | 0.0 |
| M1R | 0.0 | 0.0 | 0.0 | 0.0 | 0.0 | 0.0 | 0.0 | 0.0 |
| M2R | 0.0 | 0.0376 | 0.0 | 0.0027 | 0.0 | 0.0 | 0.0 | 0.0 |
| M3R | 0.0 | 0.0 | 0.0 | 0.1056 | 0.0 | 0.0 | 0.0 | 0.0 |
| M4R | 0.0 | 0.0 | 0.0 | 0.0 | 0.0 | 0.0 | 0.0 | 0.0 |
| A1R | 0.0 | 0.0007 | 0.0527 | 0.0133 | 0.0 | 0.0 | 0.0 | 0.0 |
| A2R | 0.0 | 0.0 | 0.0 | 0.0701 | 0.0 | 0.0 | 0.0 | 0.0 |
| A3R | 0.0 | 0.0 | 0.0 | 0.055 | 0.0 | 0.0 | 0.0 | 0.0 |
| D1R | 0.1213 | 0.0801 | 0.0348 | 0.0355 | 0.0 | 0.0 | 0.0 | 0.0 |
| D2R | 0.0 | 0.0226 | 0.0431 | 0.0133 | 0.0 | 0.0 | 0.0 | 0.0 |
| D3R | 0.8741 | 0.076 | 0.066 | 0.7045 | 0.0 | 0.0 | 0.0 | 0.0 |
| D4R | 0.0046 | 0.0418 | 0.0165 | 0.0 | 0.0 | 0.0 | 0.0 | 0.0 |
| S1R | 0.0 | 0.7009 | 0.4101 | 0.0 | 0.0 | 0.0 | 0.0 | 0.0 |
| S2R | 0.0 | 0.0404 | 0.3753 | 0.0 | 0.0 | 0.0 | 0.0 | 0.0 |
| S3R | 0.0 | 0.0 | 0.0 | 0.0 | 0.0 | 0.0 | 0.0 | 0.0 |
| S4R | 0.0 | 0.0 | 0.0016 | 0.0 | 0.0 | 0.0 | 0.0 | 0.0 |
| S5R | 0.0 | 0.0 | 0.0 | 0.0 | 0.0 | 0.0 | 0.0 | 0.0 |

**Supplemental Table 3:** Reading vs. listening in the cerebellar language regions. A linear mixed-effects (LME) model was used to evaluate the responses during reading vs. listening in the cerebellar language regions. The model included two binary, treatment-coded fixed effects: ‘Effect’ (0.5 for language, −0.5 for the control condition) and ‘Response’ (0.5 for the responses to the listening-based localizer, −0.5 for the responses to the reading-based localizer). The model included each of these individual terms and their interactions. A separate model was fit per cerebellar language region. The LME models were fit with restricted maximum likelihood estimation and included ‘Participant’ as a random effect. Bolded terms are significant (p < 0.05).

|  | *ROI* | *Estimate* | *Std..Error* | *df* | *t.value* | *Pr...t..* |
| --- | --- | --- | --- | --- | --- | --- |
| **(Intercept)** | **Right_lateral_CrusI** | **0.6071** | **0.05245** | **168** | **11.57** | **3.713e-23** |
| **EffectS** | **Right_lateral_CrusI** | **0.263** | **0.04823** | **252** | **5.453** | **1.181e-07** |
| **ResponseAuditory** | **Right_lateral_CrusI** | **-0.2396** | **0.04823** | **252** | **-4.968** | **1.247e-06** |
| *EffectS:ResponseAuditory* | *Right_lateral_CrusI* | *-0.03808* | *0.06821* | *252* | *-0.5583* | *0.5772* |
| **(Intercept)** | **Right_posterior_VI_CrusI** | **0.1923** | **0.03819** | **215.7** | **5.037** | **9.992e-07** |
| **EffectS** | **Right_posterior_VI_CrusI** | **0.2837** | **0.04073** | **252** | **6.965** | **2.845e-11** |
| *ResponseAuditory* | *Right_posterior_VI_CrusI* | *0.07738* | *0.04073* | *252* | *1.9* | *0.05862* |
| *EffectS:ResponseAuditory* | *Right_posterior_VI_CrusI* | *-0.06437* | *0.0576* | *252* | *-1.117* | *0.2649* |
| **(Intercept)** | **Right_CrusI_II_VIIb** | **0.1321** | **0.03094** | **182.8** | **4.269** | **3.155e-05** |
| **EffectS** | **Right_CrusI_II_VIIb** | **0.3206** | **0.03005** | **252** | **10.67** | **3.587e-22** |
| **ResponseAuditory** | **Right_CrusI_II_VIIb** | **0.1675** | **0.03005** | **252** | **5.574** | **6.38e-08** |
| *EffectS:ResponseAuditory* | *Right_CrusI_II_VIIb* | *-0.001361* | *0.0425* | *252* | *-0.03204* | *0.9745* |
| **(Intercept)** | **Right_VIIb_VIIIa** | **0.3325** | **0.03983** | **156.5** | **8.347** | **3.457e-14** |
| **EffectS** | **Right_VIIb_VIIIa** | **0.3381** | **0.03479** | **252** | **9.717** | **3.677e-19** |
| *ResponseAuditory* | *Right_VIIb_VIIIa* | *0.06125* | *0.03479* | *252* | *1.76* | *0.07958* |
| *EffectS:ResponseAuditory* | *Right_VIIb_VIIIa* | *-0.06923* | *0.04921* | *252* | *-1.407* | *0.1607* |

**Supplemental Table 4:** The neocortical language network’s non-linguistic responses relative to baseline. A linear mixed-effects (LME) model was used to evaluate responses in the neocortical language network to the non-linguistic tasks (Expts. 2a-e, **Fig. 3a**) relative to a fixation baseline. The model included one dummy-coded fixed effect with 32 levels (one for each of the non-linguistic conditions): ‘Effect’. The LME models were fit with restricted maximum likelihood estimation and included ‘Participant’ and ‘ROI’ (for the five ‘core’ left hemisphere language regions, <u>Methods</u>) as random effects. Bolded terms are significant (p<0.05).

|  | *Estimate* | *Std..Error* | *df* | *t.value* | *Pr...t..* |
| --- | --- | --- | --- | --- | --- |
| **EffectArticLoc_P** | **0.5587** | **0.2327** | **7.392** | **2.401** | **0.0456** |
| EffectArticLoc_T | 0.3323 | 0.2327 | 7.392 | 1.428 | 0.1942 |
| **EffectCloudy_Mental** | **0.5681** | **0.2132** | **5.205** | **2.665** | **0.04285** |
| EffectCloudy_Pain | 0.2867 | 0.2132 | 5.205 | 1.345 | 0.2343 |
| EffectCloudy_Physical | 0.4525 | 0.2132 | 5.205 | 2.123 | 0.08501 |
| **EffectCloudy_Social** | **0.8381** | **0.2132** | **5.205** | **3.932** | **0.01021** |
| EffectDigSpan_E | 0.1091 | 0.2415 | 8.567 | 0.4517 | 0.6627 |
| EffectDigSpan_H | 0.1117 | 0.2415 | 8.567 | 0.4628 | 0.6551 |
| EffectDyLoc_Bodies | 0.3486 | 0.2118 | 5.073 | 1.646 | 0.1599 |
| EffectDyLoc_Faces | 0.3973 | 0.2118 | 5.073 | 1.876 | 0.1187 |
| EffectDyLoc_Objects | 0.219 | 0.2118 | 5.073 | 1.034 | 0.348 |
| EffectDyLoc_Scenes | 0.3253 | 0.2118 | 5.073 | 1.536 | 0.1843 |
| EffectDyLoc_ScrObjects | 0.07864 | 0.2118 | 5.073 | 0.3713 | 0.7254 |
| EffectMath_E | -0.05051 | 0.2071 | 4.639 | -0.2439 | 0.8177 |
| EffectMath_H | -0.0188 | 0.2071 | 4.639 | -0.09078 | 0.9315 |
| EffectMDloc_E | 0.04858 | 0.2012 | 4.13 | 0.2415 | 0.8207 |
| EffectMDloc_H | -0.06162 | 0.2012 | 4.13 | -0.3063 | 0.7742 |
| EffectMSIT_E | -0.03078 | 0.2452 | 9.115 | -0.1255 | 0.9028 |
| EffectMSIT_H | -0.2231 | 0.2452 | 9.115 | -0.9098 | 0.3864 |
| EffectMusic_2009_B | -0.08999 | 0.2817 | 15.82 | -0.3195 | 0.7536 |
| EffectMusic_2009_I | -0.08396 | 0.2817 | 15.82 | -0.298 | 0.7695 |
| EffectMusic_2019_ANIM | -0.1009 | 0.2283 | 6.844 | -0.4422 | 0.672 |
| EffectMusic_2019_ENV_A | -0.2235 | 0.2283 | 6.844 | -0.9791 | 0.3609 |
| EffectMusic_2019_MUS_DR | -0.2803 | 0.2283 | 6.844 | -1.228 | 0.2601 |
| EffectMusic_2019_MUS_ORCH | -0.0006855 | 0.2283 | 6.844 | -0.003003 | 0.9977 |
| EffectMusic_2019_MUS_SOLO | -0.3213 | 0.2283 | 6.844 | -1.408 | 0.203 |
| EffectPic_Perc | 0.354 | 0.2181 | 5.707 | 1.623 | 0.1583 |
| **EffectPic_Sem** | **0.9297** | **0.2181** | **5.707** | **4.262** | **0.005946** |
| EffectvMSIT_E | 0.05653 | 0.243 | 8.793 | 0.2326 | 0.8214 |
| EffectvMSIT_H | -0.1031 | 0.243 | 8.793 | -0.4243 | 0.6816 |
| EffectvstroopBL_E | 0.148 | 0.2487 | 9.643 | 0.595 | 0.5656 |
| EffectvstroopBL_H | 0.2537 | 0.2487 | 9.643 | 1.02 | 0.3326 |

**Supplemental Table 5:** The neocortical language network’s non-linguistic responses relative to the control condition from the language localizer. A linear mixed-effects (LME) model was used to evaluate responses in the neocortical language network to the non-linguistic tasks (Expts. 2a-e, **Fig. 3a**) relative to the control condition from the language localizer. The model included one treatment-coded fixed effect with 33 levels (one for each of the non-linguistic conditions, and one for the control condition from the language localizer which was also the reference level): ‘Effect’. The LME models were fit with restricted maximum likelihood estimation and included ‘Participant’ and ‘ROI’ (for the five ‘core’ left hemisphere language regions, <u>Methods</u>) as random effects. Bolded terms are significant (p<0.05).

|  | Estimate | Std..Error | df | t.value | Pr...t.. |
| --- | --- | --- | --- | --- | --- |
| **(Intercept)** | **0.5913** | **0.1914** | **4.091** | **3.09** | **0.03551** |
| *EffectArticLoc_P* | -0.03261 | 0.1079 | 1594 | -0.3023 | 0.7624 |
| **EffectArticLoc_T** | **-0.259** | **0.1079** | **1594** | **-2.401** | **0.01645** |
| *EffectCloudy_Mental* | 0.0515 | 0.06849 | 13990 | 0.752 | 0.4521 |
| **EffectCloudy_Pain** | **-0.2299** | **0.06849** | **13990** | **-3.357** | **0.0007894** |
| *EffectCloudy_Physical* | -0.06407 | 0.06849 | 13990 | -0.9355 | 0.3496 |
| **EffectCloudy_Social** | **0.3215** | **0.06849** | **13990** | **4.694** | **2.703e-06** |
| *EffectDigSpan_E* | -0.3696 | 0.1096 | 11350 | -3.373 | 0.0007456 |
| *EffectDigSpan_H* | -0.367 | 0.1096 | 11350 | -3.349 | 0.0008141 |
| *EffectDyLoc_Bodies* | -0.2675 | 0.06353 | 14930 | -4.21 | 2.568e-05 |
| *EffectDyLoc_Faces* | -0.2188 | 0.06353 | 14930 | -3.443 | 0.0005768 |
| *EffectDyLoc_Objects* | -0.3971 | 0.06353 | 14930 | -6.25 | 4.222e-10 |
| *EffectDyLoc_Scenes* | -0.2907 | 0.06353 | 14930 | -4.575 | 4.79e-06 |
| *EffectDyLoc_ScrObjects* | -0.5374 | 0.06353 | 14930 | -8.458 | 2.962e-17 |
| *EffectMath_E* | -0.6514 | 0.05005 | 14920 | -13.01 | 1.646e-38 |
| *EffectMath_H* | -0.6197 | 0.05005 | 14920 | -12.38 | 4.938e-35 |
| *EffectMDloc_E* | -0.5625 | 0.02099 | 14910 | -26.8 | 1.616e-154 |
| *EffectMDloc_H* | -0.6728 | 0.02099 | 14910 | -32.05 | 5.868e-218 |
| *EffectMSIT_E* | -0.5465 | 0.117 | 14000 | -4.669 | 3.056e-06 |
| *EffectMSIT_H* | -0.7388 | 0.117 | 14000 | -6.312 | 2.837e-10 |
| *EffectMusic_2009_B* | -0.5083 | 0.1458 | 8124 | -3.486 | 0.0004922 |
| *EffectMusic_2009_I* | -0.5023 | 0.1458 | 8124 | -3.445 | 0.000574 |
| *EffectMusic_2019_ANIM* | -0.7506 | 0.103 | 14950 | -7.285 | 3.378e-13 |
| *EffectMusic_2019_ENV_A* | -0.8732 | 0.103 | 14950 | -8.474 | 2.584e-17 |
| *EffectMusic_2019_MUS_DR* | -0.9299 | 0.103 | 14950 | -9.025 | 2.006e-19 |
| *EffectMusic_2019_MUS_ORCH* | -0.6504 | 0.103 | 14950 | -6.312 | 2.833e-10 |
| *EffectMusic_2019_MUS_SOLO* | -0.971 | 0.103 | 14950 | -9.424 | 4.987e-21 |
| *EffectPic_Perc* | -0.2139 | 0.07481 | 10530 | -2.86 | 0.00425 |
| *EffectPic_Sem* | 0.3618 | 0.07481 | 10530 | 4.836 | 1.342e-06 |
| *EffectvMSIT_E* | -0.4705 | 0.1176 | 14560 | -4.002 | 6.304e-05 |
| *EffectvMSIT_H* | -0.6301 | 0.1176 | 14560 | -5.36 | 8.435e-08 |
| *EffectvstroopBL_E* | -0.3781 | 0.1232 | 14440 | -3.069 | 0.002152 |
| *EffectvstroopBL_H* | -0.2723 | 0.1232 | 14440 | -2.211 | 0.02708 |

**Supplemental Table 6:** The neocortical language network’s non-linguistic responses relative to language. A linear mixed-effects (LME) model was used to evaluate responses in the neocortical language network to the non-linguistic tasks (Expts. 2a-e, **Fig. 3a**) relative to language. The model included one treatment-coded fixed effect with 33 levels (one for each of the non-linguistic conditions, and one for the control condition from the language localizer which was also the reference level): ‘Effect’. The LME models were fit with restricted maximum likelihood estimation and included ‘Participant’ and ‘ROI’ (for the five ‘core’ left hemisphere language regions, <u>Methods</u>) as random effects. Bolded terms are significant (p<0.05).

|  | *Estimate* | *Std..Error* | *df* | *t.value* | *Pr...t..* |
| --- | --- | --- | --- | --- | --- |
| **(Intercept)** | **2.012** | **0.2129** | **4.095** | **9.45** | **0.0006246** |
| **EffectArticLoc_P** | **-1.454** | **0.1227** | **1442** | **-11.85** | **5.732e-31** |
| **EffectArticLoc_T** | **-1.68** | **0.1227** | **1442** | **-13.69** | **3.248e-40** |
| **EffectCloudy_Mental** | **-1.462** | **0.07639** | **14080** | **-19.14** | **1.165e-80** |
| **EffectCloudy_Pain** | **-1.744** | **0.07639** | **14080** | **-22.83** | **2.765e-113** |
| **EffectCloudy_Physical** | **-1.578** | **0.07639** | **14080** | **-20.66** | **2.078e-93** |
| **EffectCloudy_Social** | **-1.192** | **0.07639** | **14080** | **-15.61** | **1.814e-54** |
| **EffectDigSpan_E** | **-1.608** | **0.1225** | **11400** | **-13.13** | **4.389e-39** |
| **EffectDigSpan_H** | **-1.605** | **0.1225** | **11400** | **-13.11** | **5.822e-39** |
| **EffectDyLoc_Bodies** | **-1.683** | **0.07076** | **14950** | **-23.78** | **1.013e-122** |
| **EffectDyLoc_Faces** | **-1.634** | **0.07076** | **14950** | **-23.09** | **6.011e-116** |
| **EffectDyLoc_Objects** | **-1.812** | **0.07076** | **14950** | **-25.61** | **1.269e-141** |
| **EffectDyLoc_Scenes** | **-1.706** | **0.07076** | **14950** | **-24.11** | **5.179e-126** |
| **EffectDyLoc_ScrObjects** | **-1.953** | **0.07076** | **14950** | **-27.59** | **1.582e-163** |
| **EffectMath_E** | **-2.074** | **0.05569** | **14890** | **-37.23** | **3.494e-290** |
| **EffectMath_H** | **-2.042** | **0.05569** | **14890** | **-36.66** | **8.268e-282** |
| **EffectMDloc_E** | **-2.003** | **0.02336** | **14890** | **-85.75** | **0** |
| **EffectMDloc_H** | **-2.113** | **0.02336** | **14890** | **-90.46** | **0** |
| **EffectMSIT_E** | **-1.809** | **0.1306** | **14080** | **-13.86** | **2.258e-43** |
| **EffectMSIT_H** | **-2.001** | **0.1306** | **14080** | **-15.33** | **1.3e-52** |
| **EffectMusic_2009_B** | **-1.811** | **0.1634** | **8091** | **-11.08** | **2.427e-28** |
| **EffectMusic_2009_I** | **-1.805** | **0.1634** | **8091** | **-11.05** | **3.641e-28** |
| **EffectMusic_2019_ANIM** | **-2.228** | **0.1147** | **14970** | **-19.42** | **5.694e-83** |
| **EffectMusic_2019_ENV_A** | **-2.351** | **0.1147** | **14970** | **-20.49** | **5.233e-92** |
| **EffectMusic_2019_MUS_DR** | **-2.407** | **0.1147** | **14970** | **-20.98** | **2.385e-96** |
| **EffectMusic_2019_MUS_ORCH** | **-2.128** | **0.1147** | **14970** | **-18.54** | **6.425e-76** |
| **EffectMusic_2019_MUS_SOLO** | **-2.448** | **0.1147** | **14970** | **-21.34** | **1.512e-99** |
| **EffectPic_Perc** | **-1.642** | **0.08367** | **10590** | **-19.62** | **3.139e-84** |
| **EffectPic_Sem** | **-1.066** | **0.08367** | **10590** | **-12.74** | **6.535e-37** |
| **EffectvMSIT_E** | **-1.723** | **0.131** | **14600** | **-13.15** | **2.923e-39** |
| **EffectvMSIT_H** | **-1.882** | **0.131** | **14600** | **-14.37** | **1.747e-46** |
| **EffectvstroopBL_E** | **-1.647** | **0.1373** | **14500** | **-11.99** | **5.485e-33** |
| **EffectvstroopBL_H** | **-1.541** | **0.1373** | **14500** | **-11.22** | **4.107e-29** |

**Supplemental Table 7:** The neocortical language network’s responses within non-linguistic domains. Linear mixed-effects (LME) models were used to evaluate within-domain hypotheses in the neocortical language network (**Fig. 3a**). Each row corresponds to a term from an LME model testing a particular within-domain hypothesis. All LME models were fit with maximum likelihood estimation and additionally included ‘Participant’ and ‘ROI’ (for the five ‘core’ left hemisphere language regions, <u>Methods</u>) as a random effect. To test if there were stronger responses during articulation than during finger tapping (Expt. 2a) an LME model with one binary, treatment-coded fixed effect was used (0 for finger tapping, 1 for articulation; ‘EffectArticLoc_P’). To test whether there were overall stronger responses to harder cognitively demanding executive function tasks (Expt. 2b), an LME model was used with one binary, treatment-coded fixed effect (0 for the Easy conditions, 1 for the Hard conditions; ‘ConditionHard’) and an additional random intercept for ‘Experiment’ (spatial working memory, verbal working memory, etc.). To test if there were stronger responses while listening to structured musical excerpts than while listening to unstructured sounds (Expt. 2c) an LME model was used with one binary, treatment-coded fixed effect (0 for conditions with unstructured sounds, 1 for conditions with structured musical excerpts; ‘ConditionStructured’) with an additional random intercept for ‘Experiment’. To test whether there were stronger responses to visual social stimuli (faces and bodies) than non-social stimuli (scenes and objects, Expt. 2d) an LME model was used with one binary, treatment-coded fixed effect (0 for non-social conditions, 1 for social conditions; ‘ConditionSocial’). To test whether there were stronger responses to mental than physical content (Expt. 2e-i) an LME model was used with one binary, treatment-coded fixed effect (0 for physical content, 1 for mental content; ‘EffectCloudy_Mental’). To test whether there were stronger responses to social than pain content (Expt. 2e-i) an LME model was used with one binary, treatment-coded fixed effect (0 for pain content, 1 for social content; ‘EffectCloudy_Social’). To test whether there were stronger responses when participants were engaged in an event semantics task than when they were engaged in a perceptual task with the same stimuli (Expt. 2e-i) an LME model was used with one binary, treatment-coded fixed effect (0 for the perceptual task, 1 for the semantic task; ‘EffectPic_Sem’). Bolded terms are significant (p<0.05).

|  | *Effect* | *ROI* | *Estimate* | *Std..Error* | *df* | *t.value* | *P* |
| --- | --- | --- | --- | --- | --- | --- | --- |
| *EffectArticLoc_P* | Articulation_vs_Tapping | LH_Language | 0.2264 | 0.1228 | 292.9 | 1.843 | 0.0663 |
| ***ConditionHard*** | **Hard_vs_Easy_executive_function_tasks** | **LH_Language** | **-0.09259** | **0.01606** | **7073** | **-5.764** | **8.546e-09** |
| *ConditionStructured* | Structured_vs_Unstructured_music_perception | LH_Language | -0.02937 | 0.04104 | 538 | -0.7157 | 0.4745 |
| ***ConditionSocial*** | **Social_vs_Non-social_high_level_visual_perception** | **LH_Language** | **0.1653** | **0.03839** | **1172** | **4.305** | **1.812e-05** |
| *EffectCloudy_Mental* | Mental_vs_Physical_visual_event_semantics | LH_Language | 0.1156 | 0.1102 | 409.9 | 1.049 | 0.2948 |
| ***EffectCloudy_Social*** | **Social_vs_Pain_visual_event_semantics** | **LH_Language** | **0.5515** | **0.1186** | **414** | **4.649** | **4.495e-06** |
| ***EffectPic_Sem*** | **Semantic_vs_Perceptual_visual_task** | **LH_Language** | **0.5757** | **0.07812** | **406.2** | **7.369** | **9.675e-13** |

**Supplemental Table 8:** LangCereb1 (right lateral VI)’s non-linguistic responses relative to baseline. A linear mixed-effects (LME) model was used to evaluate responses in LangCereb1 to the non-linguistic tasks (Expts. 2a-e, **Fig. 3b**) relative to a fixation baseline, as in **Suppl. Table 4**, except ‘ROI’ was not included as a random effect. Bolded terms are significant (p<0.05).

|  | *Estimate* | *Std..Error* | *df* | *t.value* | *Pr...t..* |
| --- | --- | --- | --- | --- | --- |
| **EffectArticLoc_P** | **0.7867** | **0.118** | **1258** | **6.664** | **3.974e-11** |
| **EffectArticLoc_T** | **1.027** | **0.118** | **1258** | **8.702** | **1.007e-17** |
| **EffectCloudy_Mental** | **0.4627** | **0.08015** | **2126** | **5.772** | **8.958e-09** |
| **EffectCloudy_Pain** | **0.5734** | **0.08015** | **2126** | **7.154** | **1.153e-12** |
| **EffectCloudy_Physical** | **0.5278** | **0.08015** | **2126** | **6.585** | **5.715e-11** |
| **EffectCloudy_Social** | **0.5233** | **0.08015** | **2126** | **6.529** | **8.237e-11** |
| **EffectDigSpan_E** | **0.5643** | **0.1392** | **1978** | **4.053** | **5.257e-05** |
| **EffectDigSpan_H** | **0.909** | **0.1392** | **1978** | **6.528** | **8.43e-11** |
| EffectDyLoc_Bodies | 0.08293 | 0.07619 | 2076 | 1.089 | 0.2765 |
| EffectDyLoc_Faces | 0.04643 | 0.07619 | 2076 | 0.6094 | 0.5423 |
| EffectDyLoc_Objects | 0.1011 | 0.07619 | 2076 | 1.327 | 0.1847 |
| EffectDyLoc_Scenes | 0.1467 | 0.07619 | 2076 | 1.926 | 0.05426 |
| **EffectDyLoc_ScrObjects** | **0.1666** | **0.07619** | **2076** | **2.187** | **0.02888** |
| **EffectMath_E** | **0.3165** | **0.05972** | **2055** | **5.299** | **1.287e-07** |
| **EffectMath_H** | **0.5641** | **0.05972** | **2055** | **9.445** | **9.283e-21** |
| **EffectMDloc_E** | **0.6181** | **0.02625** | **1360** | **23.55** | **3.958e-103** |
| **EffectMDloc_H** | **0.8072** | **0.02625** | **1360** | **30.75** | **4.135e-158** |
| **EffectMSIT_E** | **0.4534** | **0.1484** | **2192** | **3.055** | **0.00228** |
| **EffectMSIT_H** | **0.6216** | **0.1484** | **2192** | **4.188** | **2.919e-05** |
| EffectMusic_2009_B | -0.07325 | 0.1958 | 1258 | -0.3742 | 0.7083 |
| EffectMusic_2009_I | 0.1144 | 0.1958 | 1258 | 0.5846 | 0.5589 |
| EffectMusic_2019_ANIM | 0.1719 | 0.1201 | 1867 | 1.432 | 0.1523 |
| EffectMusic_2019_ENV_A | 0.09944 | 0.1201 | 1867 | 0.8282 | 0.4077 |
| EffectMusic_2019_MUS_DR | 0.2139 | 0.1201 | 1867 | 1.782 | 0.07493 |
| **EffectMusic_2019_MUS_ORCH** | **0.2501** | **0.1201** | **1867** | **2.083** | **0.03736** |
| EffectMusic_2019_MUS_SOLO | 0.1194 | 0.1201 | 1867 | 0.9944 | 0.3201 |
| **EffectPic_Perc** | **0.864** | **0.09172** | **2153** | **9.42** | **1.122e-20** |
| **EffectPic_Sem** | **0.9722** | **0.09172** | **2153** | **10.6** | **1.26e-25** |
| **EffectvMSIT_E** | **0.4845** | **0.1455** | **2219** | **3.329** | **0.0008865** |
| **EffectvMSIT_H** | **0.9533** | **0.1455** | **2219** | **6.55** | **7.138e-11** |
| **EffectvstroopBL_E** | **0.9082** | **0.1553** | **2216** | **5.848** | **5.696e-09** |
| **EffectvstroopBL_H** | **1.023** | **0.1553** | **2216** | **6.587** | **5.581e-11** |

**Supplemental Table 9:** LangCereb1 (right lateral VI)’s non-linguistic responses relative to the control condition from the language localizer. A linear mixed-effects (LME) model was used to evaluate responses in LangCereb1 to the non-linguistic tasks (Expts. 2a-e, **Fig. 3b**) relative to the control condition from the language localizer, as in **Suppl. Table 5**, except ‘ROI’ was not included as a random effect. Bolded terms are significant (p<0.05).

|  | *Estimate* | *Std..Error* | *df* | *t.value* | *Pr...t..* |
| --- | --- | --- | --- | --- | --- |
| **(Intercept)** | **0.5303** | **0.02302** | **2061** | **23.04** | **1.066e-104** |
| **EffectArticLoc_P** | **0.2564** | **0.1151** | **1844** | **2.226** | **0.02611** |
| **EffectArticLoc_T** | **0.4969** | **0.1151** | **1844** | **4.315** | **1.678e-05** |
| *EffectCloudy_Mental* | -0.03478 | 0.08247 | 2749 | -0.4217 | 0.6733 |
| *EffectCloudy_Pain* | 0.07598 | 0.08247 | 2749 | 0.9212 | 0.357 |
| *EffectCloudy_Physical* | 0.03035 | 0.08247 | 2749 | 0.368 | 0.7129 |
| *EffectCloudy_Social* | 0.02589 | 0.08247 | 2749 | 0.3139 | 0.7536 |
| *EffectDigSpan_E* | -0.02351 | 0.1299 | 2921 | -0.181 | 0.8564 |
| **EffectDigSpan_H** | **0.3212** | **0.1299** | **2921** | **2.472** | **0.01349** |
| **EffectDyLoc_Bodies** | **-0.3487** | **0.07738** | **2545** | **-4.506** | **6.917e-06** |
| **EffectDyLoc_Faces** | **-0.3852** | **0.07738** | **2545** | **-4.977** | **6.877e-07** |
| **EffectDyLoc_Objects** | **-0.3305** | **0.07738** | **2545** | **-4.271** | **2.018e-05** |
| **EffectDyLoc_Scenes** | **-0.2849** | **0.07738** | **2545** | **-3.681** | **0.000237** |
| **EffectDyLoc_ScrObjects** | **-0.265** | **0.07738** | **2545** | **-3.424** | **0.0006258** |
| **EffectMath_E** | **-0.2317** | **0.06145** | **2374** | **-3.77** | **0.0001673** |
| *EffectMath_H* | 0.01593 | 0.06145 | 2374 | 0.2593 | 0.7954 |
| **EffectMDloc_E** | **0.09289** | **0.02581** | **2318** | **3.599** | **0.000326** |
| **EffectMDloc_H** | **0.282** | **0.02581** | **2318** | **10.93** | **3.916e-27** |
| *EffectMSIT_E* | -0.1366 | 0.141 | 2733 | -0.9683 | 0.333 |
| *EffectMSIT_H* | 0.03173 | 0.141 | 2733 | 0.225 | 0.822 |
| **EffectMusic_2009_B** | **-0.5248** | **0.1697** | **2963** | **-3.092** | **0.002005** |
| **EffectMusic_2009_I** | **-0.3371** | **0.1697** | **2963** | **-1.986** | **0.04709** |
| **EffectMusic_2019_ANIM** | **-0.4069** | **0.1256** | **2532** | **-3.24** | **0.001211** |
| **EffectMusic_2019_ENV_A** | **-0.4794** | **0.1256** | **2532** | **-3.817** | **0.0001381** |
| **EffectMusic_2019_MUS_DR** | **-0.3649** | **0.1256** | **2532** | **-2.906** | **0.003697** |
| **EffectMusic_2019_MUS_ORCH** | **-0.3287** | **0.1256** | **2532** | **-2.617** | **0.008911** |
| **EffectMusic_2019_MUS_SOLO** | **-0.4595** | **0.1256** | **2532** | **-3.658** | **0.0002589** |
| **EffectPic_Perc** | **0.3252** | **0.08826** | **2955** | **3.685** | **0.0002327** |
| **EffectPic_Sem** | **0.4335** | **0.08826** | **2955** | **4.911** | **9.532e-07** |
| *EffectvMSIT_E* | -0.07263 | 0.1424 | 2625 | -0.5099 | 0.6102 |
| **EffectvMSIT_H** | **0.3962** | **0.1424** | **2625** | **2.781** | **0.005451** |
| **EffectvstroopBL_E** | **0.3306** | **0.149** | **2660** | **2.218** | **0.02661** |
| **EffectvstroopBL_H** | **0.4453** | **0.149** | **2660** | **2.988** | **0.002833** |

**Supplemental Table 10:** LangCereb1 (right lateral VI)’s non-linguistic responses relative to language. A linear mixed-effects (LME) model was used to evaluate responses in LangCereb1 to the non-linguistic tasks (Expts. 2a-e, **Fig. 3b**) relative to language, as in **Suppl. Table 6**, except ‘ROI’ was not included as a random effect. Bolded terms are significant (p<0.05).

|  | Estimate | Std..Error | df | t.value | Pr...t.. |
| --- | --- | --- | --- | --- | --- |
| **(Intercept)** | **0.7871** | **0.02331** | **2078** | **33.77** | **1.195e-199** |
| *EffectArticLoc_P* | -0.0004822 | 0.1165 | 1861 | -0.004137 | 0.9967 |
| **EffectArticLoc_T** | **0.24** | **0.1165** | **1861** | **2.06** | **0.03956** |
| **EffectCloudy_Mental** | **-0.3254** | **0.08388** | **2755** | **-3.879** | **0.0001074** |
| **EffectCloudy_Pain** | **-0.2146** | **0.08388** | **2755** | **-2.559** | **0.01056** |
| **EffectCloudy_Physical** | **-0.2602** | **0.08388** | **2755** | **-3.103** | **0.001938** |
| **EffectCloudy_Social** | **-0.2647** | **0.08388** | **2755** | **-3.156** | **0.001618** |
| *EffectDigSpan_E* | -0.2273 | 0.1321 | 2924 | -1.721 | 0.08527 |
| *EffectDigSpan_H* | 0.1174 | 0.1321 | 2924 | 0.8889 | 0.3741 |
| **EffectDyLoc_Bodies** | **-0.5974** | **0.07875** | **2552** | **-7.586** | **4.59e-14** |
| **EffectDyLoc_Faces** | **-0.6339** | **0.07875** | **2552** | **-8.05** | **1.26e-15** |
| **EffectDyLoc_Objects** | **-0.5792** | **0.07875** | **2552** | **-7.355** | **2.551e-13** |
| **EffectDyLoc_Scenes** | **-0.5336** | **0.07875** | **2552** | **-6.776** | **1.529e-11** |
| **EffectDyLoc_ScrObjects** | **-0.5137** | **0.07875** | **2552** | **-6.524** | **8.243e-11** |
| **EffectMath_E** | **-0.4891** | **0.06256** | **2379** | **-7.819** | **7.938e-15** |
| **EffectMath_H** | **-0.2416** | **0.06256** | **2379** | **-3.861** | **0.0001158** |
| **EffectMDloc_E** | **-0.1684** | **0.02628** | **2320** | **-6.41** | **1.761e-10** |
| *EffectMDloc_H* | 0.02066 | 0.02628 | 2320 | 0.7863 | 0.4318 |
| **EffectMSIT_E** | **-0.33** | **0.1434** | **2739** | **-2.3** | **0.0215** |
| *EffectMSIT_H* | -0.1617 | 0.1434 | 2739 | -1.127 | 0.2597 |
| **EffectMusic_2009_B** | **-0.7135** | **0.1723** | **2962** | **-4.14** | **3.564e-05** |
| **EffectMusic_2009_I** | **-0.5259** | **0.1723** | **2962** | **-3.051** | **0.002298** |
| **EffectMusic_2019_ANIM** | **-0.6807** | **0.1278** | **2539** | **-5.326** | **1.094e-07** |
| **EffectMusic_2019_ENV_A** | **-0.7532** | **0.1278** | **2539** | **-5.893** | **4.294e-09** |
| **EffectMusic_2019_MUS_DR** | **-0.6387** | **0.1278** | **2539** | **-4.997** | **6.214e-07** |
| **EffectMusic_2019_MUS_ORCH** | **-0.6025** | **0.1278** | **2539** | **-4.714** | **2.56e-06** |
| **EffectMusic_2019_MUS_SOLO** | **-0.7332** | **0.1278** | **2539** | **-5.737** | **1.079e-08** |
| *EffectPic_Perc* | 0.07896 | 0.08968 | 2957 | 0.8804 | 0.3787 |
| **EffectPic_Sem** | **0.1872** | **0.08968** | **2957** | **2.087** | **0.03694** |
| **EffectvMSIT_E** | **-0.2964** | **0.1449** | **2630** | **-2.045** | **0.04096** |
| *EffectvMSIT_H* | 0.1724 | 0.1449 | 2630 | 1.19 | 0.2342 |
| *EffectvstroopBL_E* | 0.1409 | 0.1516 | 2666 | 0.9294 | 0.3527 |
| *EffectvstroopBL_H* | 0.2556 | 0.1516 | 2666 | 1.686 | 0.0919 |

**Supplemental Table 11:** LangCereb1 (right lateral VI)’s responses within non-linguistic categories. Linear mixed-effects (LME) models were used to evaluate within-category hypotheses in LangCereb1 (**Fig. 3b**). Each row corresponds to a term from an LME model testing a particular within-category hypothesis (as in **Suppl. Table 7**). All LME models were fit with maximum likelihood estimation and additionally included ‘Participant’ and ‘ROI’ (for the five ‘core’ left hemisphere language regions, <u>Methods</u>) as a random effect. See **Suppl. Table 7** and <u>Methods</u> for additional model details. Bolded terms are significant (p<0.05).

|  | *Effect* | *ROI* | *Estimate* | *Std..Error* | *df* | *t.value* | *P.uncorrected* |
| --- | --- | --- | --- | --- | --- | --- | --- |
| *EffectArticLoc_P* | Articulation_vs_Tapping | Right_lateral_VI | -0.2405 | 0.1718 | 33 | -1.4 | 0.1708 |
| ***ConditionHard*** | **Hard_vs_Easy_executive_function_tasks** | **Right_lateral_VI** | **0.202** | **0.01232** | **869.4** | **16.4** | **7.659e-53** |
| *ConditionStructured* | Structured_vs_Unstructured_music_perception | Right_lateral_VI | 0.09036 | 0.05472 | 85.67 | 1.651 | 0.1023 |
| *ConditionSocial* | Social_vs_Non-social_high_level_visual_perception | Right_lateral_VI | -0.07346 | 0.04199 | 196 | -1.749 | 0.08178 |
| *EffectCloudy_Mental* | Mental_vs_Physical_visual_event_semantics | Right_lateral_VI | -0.06513 | 0.08463 | 46 | -0.7695 | 0.4455 |
| *EffectCloudy_Social* | Social_vs_Pain_visual_event_semantics | Right_lateral_VI | -0.05009 | 0.1037 | 46 | -0.4828 | 0.6315 |
| *EffectPic_Sem* | Semantic_vs_Perceptual_visual_task | Right_lateral_VI | 0.1082 | 0.09827 | 50.57 | 1.101 | 0.2759 |

**Supplemental Table 12:**
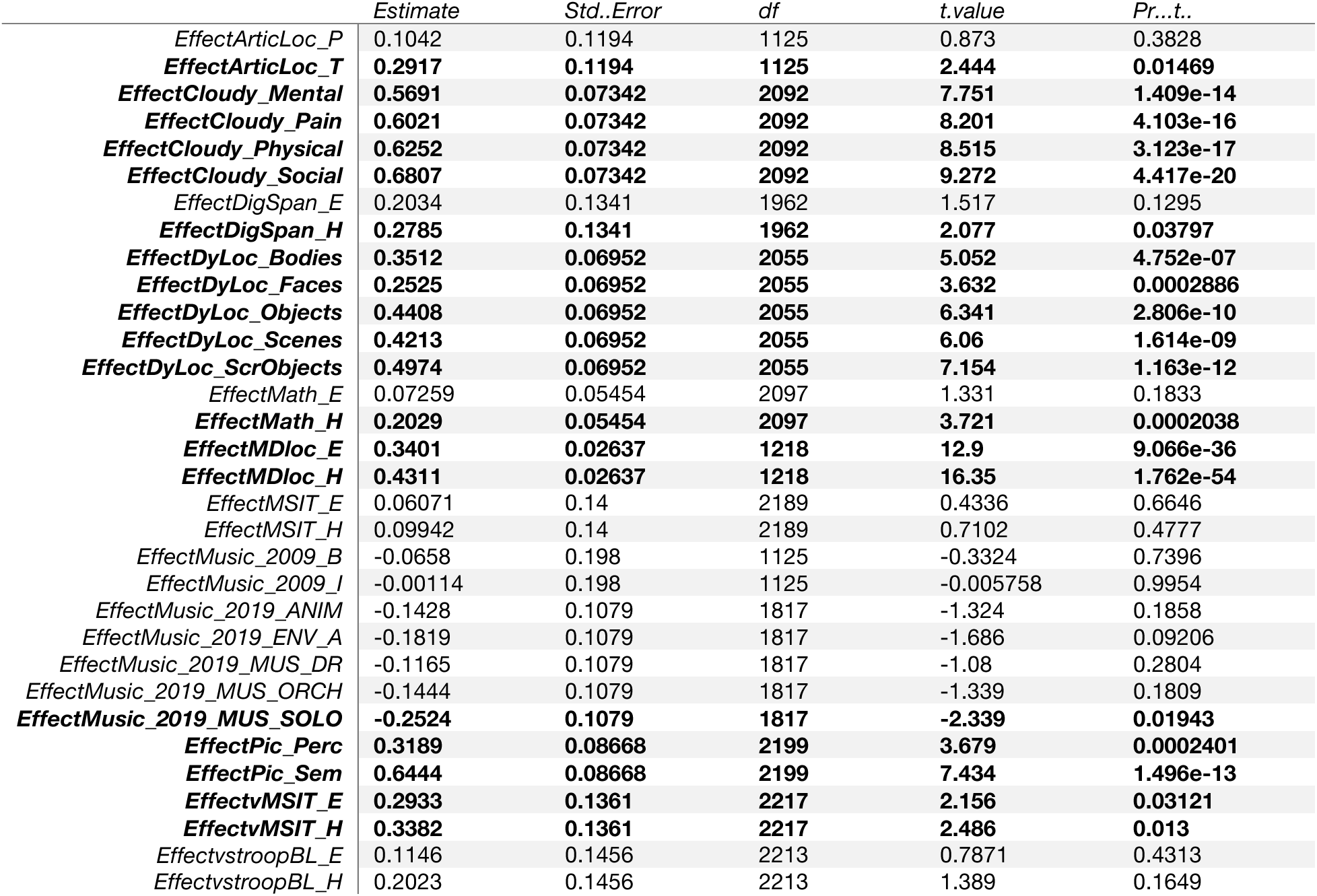
LangCereb2 (right posterior VI/Crus I)’s non-linguistic responses relative to baseline. A linear mixed-effects (LME) model was used to evaluate responses in LangCereb2 to the non-linguistic tasks (Expts. 2a-e, **Fig. 3c**) relative to a fixation baseline, as in **Suppl. Table 8**. Bolded terms are significant (p<0.05).

**Supplemental Table 13:**
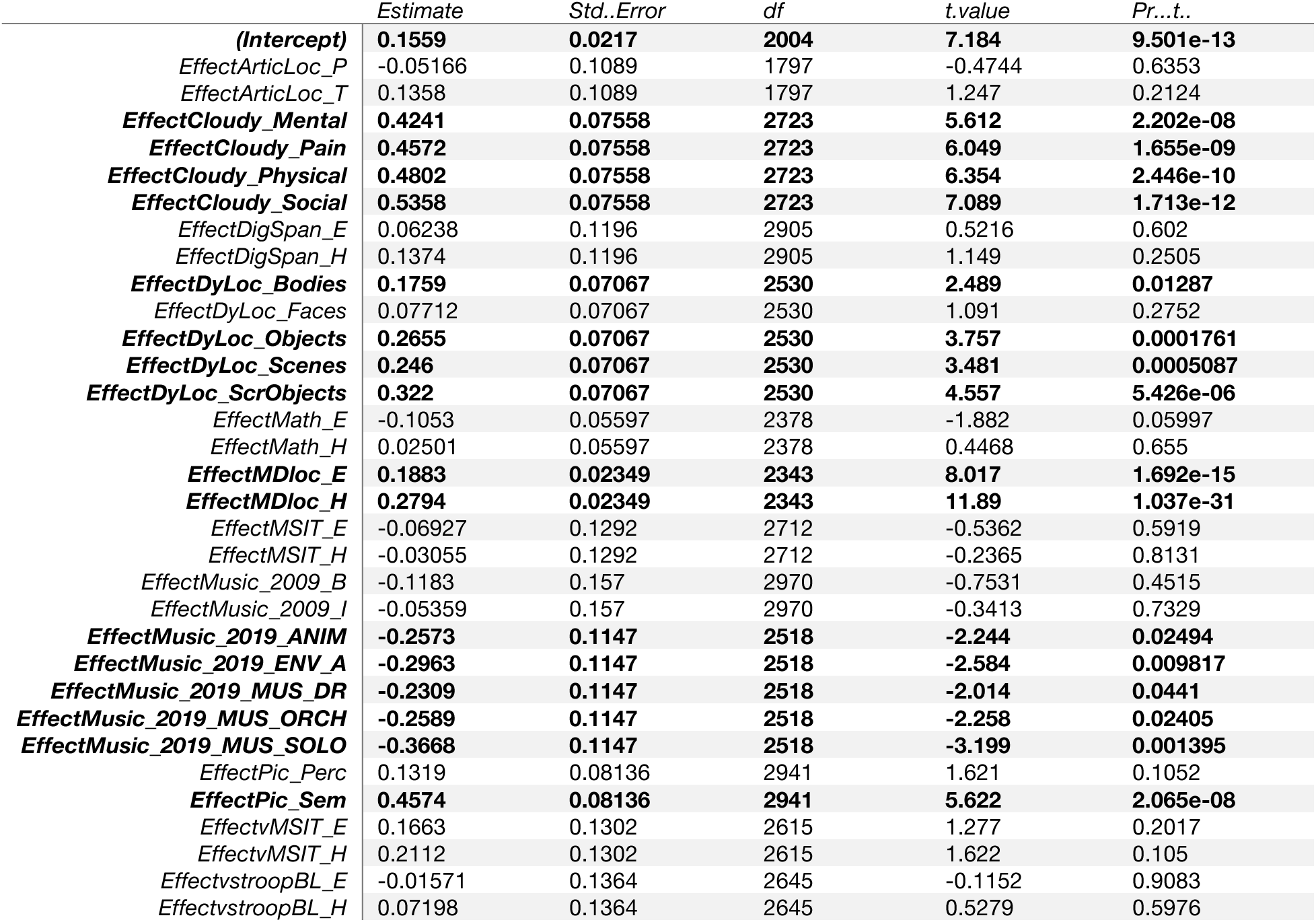
LangCereb2 (right posterior VI/Crus I)’s non-linguistic responses relative to the control condition from the language localizer. A linear mixed-effects (LME) model was used to evaluate responses in LangCereb2 to the non-linguistic tasks (Expts. 2a-e, **Fig. 3c**) relative to the control condition from the language localizer, as in **Suppl. Table 9**. Bolded terms are significant (p<0.05).

|  | *Estimate* | *Std..Error* | *df* | *t.value* | *Pr...t..* |
| --- | --- | --- | --- | --- | --- |
| **(Intercept)** | **0.1559** | **0.0217** | **2004** | **7.184** | **9.501e-13** |
| *EffectArticLoc_P* | -0.05166 | 0.1089 | 1797 | -0.4744 | 0.6353 |
| *EffectArticLoc_T* | 0.1358 | 0.1089 | 1797 | 1.247 | 0.2124 |
| **EffectCloudy_Mental** | **0.4241** | **0.07558** | **2723** | **5.612** | **2.202e-08** |
| **EffectCloudy_Pain** | **0.4572** | **0.07558** | **2723** | **6.049** | **1.655e-09** |
| **EffectCloudy_Physical** | **0.4802** | **0.07558** | **2723** | **6.354** | **2.446e-10** |
| **EffectCloudy_Social** | **0.5358** | **0.07558** | **2723** | **7.089** | **1.713e-12** |
| *EffectDigSpan_E* | 0.06238 | 0.1196 | 2905 | 0.5216 | 0.602 |
| *EffectDigSpan_H* | 0.1374 | 0.1196 | 2905 | 1.149 | 0.2505 |
| **EffectDyLoc_Bodies** | **0.1759** | **0.07067** | **2530** | **2.489** | **0.01287** |
| *EffectDyLoc_Faces* | 0.07712 | 0.07067 | 2530 | 1.091 | 0.2752 |
| **EffectDyLoc_Objects** | **0.2655** | **0.07067** | **2530** | **3.757** | **0.0001761** |
| **EffectDyLoc_Scenes** | **0.246** | **0.07067** | **2530** | **3.481** | **0.0005087** |
| **EffectDyLoc_ScrObjects** | **0.322** | **0.07067** | **2530** | **4.557** | **5.426e-06** |
| *EffectMath_E* | -0.1053 | 0.05597 | 2378 | -1.882 | 0.05997 |
| *EffectMath_H* | 0.02501 | 0.05597 | 2378 | 0.4468 | 0.655 |
| **EffectMDloc_E** | **0.1883** | **0.02349** | **2343** | **8.017** | **1.692e-15** |
| **EffectMDloc_H** | **0.2794** | **0.02349** | **2343** | **11.89** | **1.037e-31** |
| *EffectMSIT_E* | -0.06927 | 0.1292 | 2712 | -0.5362 | 0.5919 |
| *EffectMSIT_H* | -0.03055 | 0.1292 | 2712 | -0.2365 | 0.8131 |
| *EffectMusic_2009_B* | -0.1183 | 0.157 | 2970 | -0.7531 | 0.4515 |
| *EffectMusic_2009_I* | -0.05359 | 0.157 | 2970 | -0.3413 | 0.7329 |
| **EffectMusic_2019_ANIM** | **-0.2573** | **0.1147** | **2518** | **-2.244** | **0.02494** |
| **EffectMusic_2019_ENV_A** | **-0.2963** | **0.1147** | **2518** | **-2.584** | **0.009817** |
| **EffectMusic_2019_MUS_DR** | **-0.2309** | **0.1147** | **2518** | **-2.014** | **0.0441** |
| **EffectMusic_2019_MUS_ORCH** | **-0.2589** | **0.1147** | **2518** | **-2.258** | **0.02405** |
| **EffectMusic_2019_MUS_SOLO** | **-0.3668** | **0.1147** | **2518** | **-3.199** | **0.001395** |
| *EffectPic_Perc* | 0.1319 | 0.08136 | 2941 | 1.621 | 0.1052 |
| **EffectPic_Sem** | **0.4574** | **0.08136** | **2941** | **5.622** | **2.065e-08** |
| *EffectvMSIT_E* | 0.1663 | 0.1302 | 2615 | 1.277 | 0.2017 |
| *EffectvMSIT_H* | 0.2112 | 0.1302 | 2615 | 1.622 | 0.105 |
| *EffectvstroopBL_E* | -0.01571 | 0.1364 | 2645 | -0.1152 | 0.9083 |
| *EffectvstroopBL_H* | 0.07198 | 0.1364 | 2645 | 0.5279 | 0.5976 |

**Supplemental Table 14:** LangCereb2 (right posterior VI/Crus I)’s non-linguistic responses relative to language. A linear mixed-effects (LME) model was used to evaluate responses in LangCereb2 to the non-linguistic tasks (Expts. 2a-e, **Fig. 3c**) relative to language, as in **Suppl. Table 9**. Bolded terms are significant (p<0.05).

|  | Estimate | Std..Error | df | t.value | Pr...t.. |
| --- | --- | --- | --- | --- | --- |
| **(Intercept)** | **0.4781** | **0.02211** | **1973** | **21.63** | **3.021e-93** |
| **EffectArticLoc_P** | **-0.3739** | **0.111** | **1767** | **-3.367** | **0.0007764** |
| EffectArticLoc_T | -0.1864 | 0.111 | 1767 | -1.678 | 0.09344 |
| EffectCloudy_Mental | 0.09378 | 0.07652 | 2711 | 1.226 | 0.2205 |
| EffectCloudy_Pain | 0.1268 | 0.07652 | 2711 | 1.657 | 0.09758 |
| EffectCloudy_Physical | 0.1499 | 0.07652 | 2711 | 1.959 | 0.05026 |
| **EffectCloudy_Social** | **0.2054** | **0.07652** | **2711** | **2.684** | **0.007312** |
| **EffectDigSpan_E** | **-0.2592** | **0.1212** | **2899** | **-2.138** | **0.03257** |
| EffectDigSpan_H | -0.1841 | 0.1212 | 2899 | -1.519 | 0.1289 |
| EffectDyLoc_Bodies | -0.1377 | 0.0715 | 2516 | -1.926 | 0.05417 |
| **EffectDyLoc_Faces** | **-0.2365** | **0.0715** | **2516** | **-3.308** | **0.0009532** |
| EffectDyLoc_Objects | -0.04816 | 0.0715 | 2516 | -0.6736 | 0.5006 |
| EffectDyLoc_Scenes | -0.06766 | 0.0715 | 2516 | -0.9464 | 0.3441 |
| EffectDyLoc_ScrObjects | 0.008424 | 0.0715 | 2516 | 0.1178 | 0.9062 |
| **EffectMath_E** | **-0.4161** | **0.0566** | **2365** | **-7.352** | **2.679e-13** |
| **EffectMath_H** | **-0.2858** | **0.0566** | **2365** | **-5.049** | **4.779e-07** |
| **EffectMDloc_E** | **-0.1347** | **0.02375** | **2333** | **-5.672** | **1.586e-08** |
| EffectMDloc_H | -0.04369 | 0.02375 | 2333 | -1.839 | 0.06601 |
| **EffectMSIT_E** | **-0.4183** | **0.1308** | **2700** | **-3.198** | **0.001398** |
| **EffectMSIT_H** | **-0.3796** | **0.1308** | **2700** | **-2.902** | **0.003734** |
| **EffectMusic_2009_B** | **-0.451** | **0.1593** | **2971** | **-2.83** | **0.004681** |
| **EffectMusic_2009_I** | **-0.3863** | **0.1593** | **2971** | **-2.424** | **0.01539** |
| **EffectMusic_2019_ANIM** | **-0.5934** | **0.116** | **2503** | **-5.115** | **3.369e-07** |
| **EffectMusic_2019_ENV_A** | **-0.6324** | **0.116** | **2503** | **-5.452** | **5.469e-08** |
| **EffectMusic_2019_MUS_DR** | **-0.5671** | **0.116** | **2503** | **-4.889** | **1.08e-06** |
| **EffectMusic_2019_MUS_ORCH** | **-0.595** | **0.116** | **2503** | **-5.129** | **3.131e-07** |
| **EffectMusic_2019_MUS_SOLO** | **-0.703** | **0.116** | **2503** | **-6.06** | **1.567e-09** |
| **EffectPic_Perc** | **-0.1757** | **0.08248** | **2937** | **-2.13** | **0.03324** |
| EffectPic_Sem | 0.1499 | 0.08248 | 2937 | 1.817 | 0.06933 |
| EffectvMSIT_E | -0.1781 | 0.1318 | 2604 | -1.351 | 0.1769 |
| EffectvMSIT_H | -0.1332 | 0.1318 | 2604 | -1.01 | 0.3125 |
| **EffectvstroopBL_E** | **-0.3625** | **0.138** | **2633** | **-2.626** | **0.008682** |
| **EffectvstroopBL_H** | **-0.2748** | **0.138** | **2633** | **-1.991** | **0.0466** |

**Supplemental Table 15:** LangCereb2 (right posterior VI/Crus I)’s responses within non-linguistic categories. Linear mixed-effects (LME) models were used to evaluate within-category hypotheses in LangCereb2 (**Fig. 3c**). Each row corresponds to a term from an LME model testing a particular within-category hypothesis (as in **Suppl. Table 7**). All LME models were fit with maximum likelihood estimation and additionally included ‘Participant’ and ‘ROI’ (for the five ‘core’ left hemisphere language regions, <u>Methods</u>) as a random effect. See **Suppl. Table 7** and <u>Methods</u> for additional model details. Bolded terms are significant (p<0.05).

|  | *Effect* | *ROI* | *Estimate* | *Std..Error* | *df* | *t.value* | *P.uncorrected* |
| --- | --- | --- | --- | --- | --- | --- | --- |
| *EffectArticLoc_P* | Articulation_vs_Tapping | Right_posterior_VI_CrusI | -0.1875 | 0.1047 | 33 | -1.791 | 0.08 |
| ***ConditionHard*** | **Hard_vs_Easy_executive_function_tasks** | **Right_posterior_VI_CrusI** | **0.09255** | **0.01121** | **869.8** | **8.258** | **5.459e-16** |
| *ConditionStructured* | Structured_vs_Unstructured_music_perception | Right_posterior_VI_CrusI | 0.009038 | 0.0467 | 90.22 | 0.1935 | 0.847 |
| ***ConditionSocial*** | **Social_vs_Non-social_high_level_visual_perception** | **Right_posterior_VI_CrusI** | **-0.1513** | **0.05498** | **196** | **-2.752** | **0.006475** |
| *EffectCloudy_Mental* | Mental_vs_Physical_visual_event_semantics | Right_posterior_VI_CrusI | -0.0561 | 0.06661 | 46 | -0.8422 | 0.404 |
| *EffectCloudy_Social* | Social_vs_Pain_visual_event_semantics | Right_posterior_VI_CrusI | 0.07859 | 0.08369 | 46 | 0.939 | 0.3526 |
| ***EffectPic_Sem*** | **Semantic_vs_Perceptual_visual_task** | **Right_posterior_VI_CrusI** | **0.3256** | **0.07066** | **49.58** | **4.607** | **2.876e-05** |

**Supplemental Table 16:** LangCereb3 (right Crus I/II/VIIb)’s non-linguistic responses relative to baseline. A linear mixed-effects (LME) model was used to evaluate responses in LangCereb3 to the non-linguistic tasks (Expts. 2a-e, **Fig. 3d**) relative to a fixation baseline, as in **Suppl. Table 8**. Bolded terms are significant (p<0.05).

|  | *Estimate* | *Std..Error* | *df* | *t.value* | *Pr...t..* |
| --- | --- | --- | --- | --- | --- |
| **EffectArticLoc_P** | **0.158** | **0.06989** | **1295** | **2.26** | **0.02398** |
| EffectArticLoc_T | 0.1345 | 0.06989 | 1295 | 1.925 | 0.05446 |
| **EffectCloudy_Mental** | **0.222** | **0.04864** | **2137** | **4.566** | **5.265e-06** |
| **EffectCloudy_Pain** | **0.09986** | **0.04864** | **2137** | **2.053** | **0.04017** |
| **EffectCloudy_Physical** | **0.2397** | **0.04864** | **2137** | **4.929** | **8.886e-07** |
| **EffectCloudy_Social** | **0.3521** | **0.04864** | **2137** | **7.24** | **6.232e-13** |
| EffectDigSpan_E | 0.03086 | 0.08343 | 1985 | 0.3699 | 0.7115 |
| EffectDigSpan_H | -0.08876 | 0.08343 | 1985 | -1.064 | 0.2875 |
| **EffectDyLoc_Bodies** | **0.1411** | **0.04629** | **2085** | **3.049** | **0.002324** |
| **EffectDyLoc_Faces** | **0.1445** | **0.04629** | **2085** | **3.122** | **0.001819** |
| **EffectDyLoc_Objects** | **0.09327** | **0.04629** | **2085** | **2.015** | **0.04405** |
| **EffectDyLoc_Scenes** | **0.1033** | **0.04629** | **2085** | **2.231** | **0.02578** |
| **EffectDyLoc_ScrObjects** | **0.111** | **0.04629** | **2085** | **2.398** | **0.01657** |
| EffectMath_E | -0.0183 | 0.03631 | 2047 | -0.5042 | 0.6142 |
| EffectMath_H | -0.03263 | 0.03631 | 2047 | -0.8987 | 0.3689 |
| EffectMDloc_E | -0.01322 | 0.01557 | 1398 | -0.849 | 0.396 |
| **EffectMDloc_H** | **-0.08051** | **0.01557** | **1398** | **-5.171** | **2.667e-07** |
| EffectMSIT_E | 0.0168 | 0.08941 | 2193 | 0.1879 | 0.8509 |
| EffectMSIT_H | -0.00937 | 0.08941 | 2193 | -0.1048 | 0.9165 |
| EffectMusic_2009_B | 0.003233 | 0.1159 | 1295 | 0.02789 | 0.9778 |
| EffectMusic_2009_I | 0.000533 | 0.1159 | 1295 | 0.004599 | 0.9963 |
| EffectMusic_2019_ANIM | -0.008113 | 0.07323 | 1882 | -0.1108 | 0.9118 |
| EffectMusic_2019_ENV_A | 0.03552 | 0.07323 | 1882 | 0.485 | 0.6278 |
| EffectMusic_2019_MUS_DR | -0.07376 | 0.07323 | 1882 | -1.007 | 0.314 |
| EffectMusic_2019_MUS_ORCH | -0.01761 | 0.07323 | 1882 | -0.2404 | 0.81 |
| EffectMusic_2019_MUS_SOLO | -0.09522 | 0.07323 | 1882 | -1.3 | 0.1937 |
| EffectPic_Perc | 0.07566 | 0.05516 | 2137 | 1.372 | 0.1703 |
| **EffectPic_Sem** | **0.3334** | **0.05516** | **2137** | **6.044** | **1.771e-09** |
| EffectvMSIT_E | -0.0159 | 0.08794 | 2218 | -0.1808 | 0.8565 |
| **EffectvMSIT_H** | **-0.1946** | **0.08794** | **2218** | **-2.213** | **0.02698** |
| EffectvstroopBL_E | 0.08686 | 0.0937 | 2216 | 0.927 | 0.3541 |
| EffectvstroopBL_H | 0.08027 | 0.0937 | 2216 | 0.8566 | 0.3917 |

**Supplemental Table 17:** LangCereb3 (right Crus I/II/VIIb)’s non-linguistic responses relative to the control condition from the language localizer. A linear mixed-effects (LME) model was used to evaluate responses in LangCereb3 to the non-linguistic tasks (Expts. 2a-e, **Fig. 3d**) relative to the control condition from the language localizer, as in **Suppl. Table 9**. Bolded terms are significant (p<0.05).

|  | *Estimate* | *Std..Error* | *df* | *t.value* | *Pr...t..* |
| --- | --- | --- | --- | --- | --- |
| **(Intercept)** | **0.1067** | **0.01303** | **2142** | **8.19** | **4.427e-16** |
| *EffectArticLoc_P* | 0.05124 | 0.06501 | 1922 | 0.7883 | 0.4306 |
| *EffectArticLoc_T* | 0.02782 | 0.06501 | 1922 | 0.428 | 0.6687 |
| **EffectCloudy_Mental** | **0.1194** | **0.04767** | **2779** | **2.505** | **0.0123** |
| *EffectCloudy_Pain* | -0.002771 | 0.04767 | 2779 | -0.05813 | 0.9536 |
| **EffectCloudy_Physical** | **0.1371** | **0.04767** | **2779** | **2.876** | **0.004056** |
| **EffectCloudy_Social** | **0.2495** | **0.04767** | **2779** | **5.234** | **1.786e-07** |
| *EffectDigSpan_E* | -0.05636 | 0.07484 | 2935 | -0.7531 | 0.4515 |
| **EffectDigSpan_H** | **-0.176** | **0.07484** | **2935** | **-2.351** | **0.01877** |
| *EffectDyLoc_Bodies* | 0.02611 | 0.04486 | 2577 | 0.5821 | 0.5606 |
| *EffectDyLoc_Faces* | 0.0295 | 0.04486 | 2577 | 0.6576 | 0.5109 |
| *EffectDyLoc_Objects* | -0.02177 | 0.04486 | 2577 | -0.4853 | 0.6275 |
| *EffectDyLoc_Scenes* | -0.01176 | 0.04486 | 2577 | -0.2621 | 0.7932 |
| *EffectDyLoc_ScrObjects* | -0.004025 | 0.04486 | 2577 | -0.08974 | 0.9285 |
| **EffectMath_E** | **-0.1333** | **0.0357** | **2396** | **-3.735** | **0.0001918** |
| **EffectMath_H** | **-0.1477** | **0.0357** | **2396** | **-4.137** | **3.646e-05** |
| **EffectMDloc_E** | **-0.1205** | **0.015** | **2321** | **-8.028** | **1.561e-15** |
| **EffectMDloc_H** | **-0.1877** | **0.015** | **2321** | **-12.51** | **8.262e-35** |
| *EffectMSIT_E* | -0.06471 | 0.08153 | 2762 | -0.7937 | 0.4275 |
| *EffectMSIT_H* | -0.09088 | 0.08153 | 2762 | -1.115 | 0.2651 |
| *EffectMusic_2009_B* | -0.05946 | 0.09734 | 2956 | -0.6108 | 0.5414 |
| *EffectMusic_2009_I* | -0.06216 | 0.09734 | 2956 | -0.6386 | 0.5232 |
| *EffectMusic_2019_ANIM* | -0.1181 | 0.07281 | 2565 | -1.623 | 0.1048 |
| *EffectMusic_2019_ENV_A* | -0.07451 | 0.07281 | 2565 | -1.023 | 0.3062 |
| **EffectMusic_2019_MUS_DR** | **-0.1838** | **0.07281** | **2565** | **-2.524** | **0.01166** |
| *EffectMusic_2019_MUS_ORCH* | -0.1276 | 0.07281 | 2565 | -1.753 | 0.07973 |
| **EffectMusic_2019_MUS_SOLO** | **-0.2052** | **0.07281** | **2565** | **-2.819** | **0.004855** |
| *EffectPic_Perc* | -0.01236 | 0.05077 | 2964 | -0.2434 | 0.8077 |
| **EffectPic_Sem** | **0.2453** | **0.05077** | **2964** | **4.832** | **1.421e-06** |
| *EffectvMSIT_E* | -0.0882 | 0.08248 | 2648 | -1.069 | 0.285 |
| **EffectvMSIT_H** | **-0.2669** | **0.08248** | **2648** | **-3.236** | **0.001226** |
| *EffectvstroopBL_E* | 0.002134 | 0.08625 | 2687 | 0.02474 | 0.9803 |
| *EffectvstroopBL_H* | -0.004455 | 0.08625 | 2687 | -0.05165 | 0.9588 |

**Supplemental Table 18:**
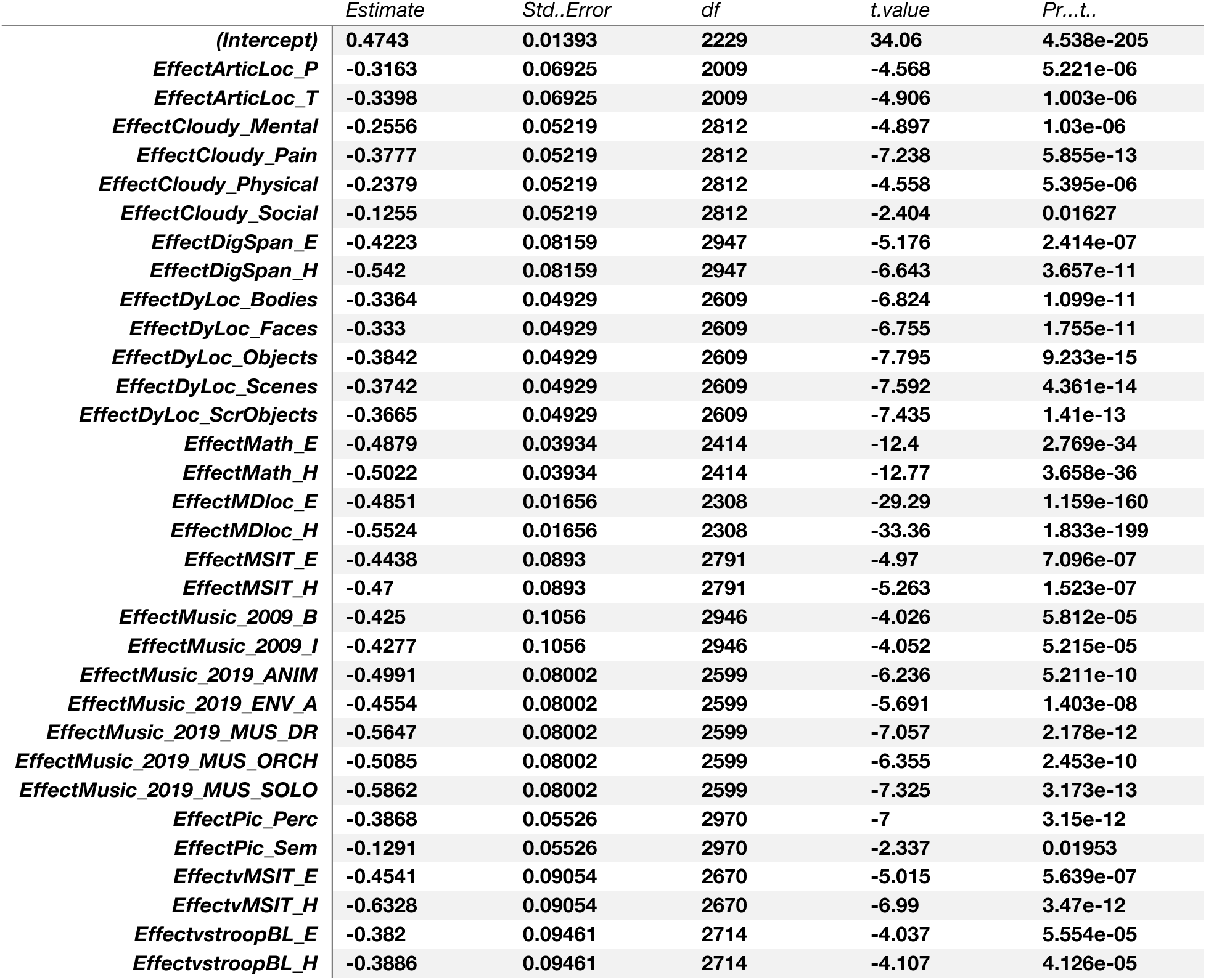
LangCereb3 (right Crus I/II/VIIb)’s non-linguistic responses relative to language. A linear mixed-effects (LME) model was used to evaluate responses in LangCereb3 to the non-linguistic tasks (Expts. 2a-e, **Fig. 3d**) relative to language, as in **Suppl. Table 9**. Bolded terms are significant (p<0.05).

**Supplemental Table 19:** LangCereb3 (right Crus I/II/VIIb)’s responses within non-linguistic categories. Linear mixed-effects (LME) models were used to evaluate within-category hypotheses in LangCereb3 (**Fig. 3d**). Each row corresponds to a term from an LME model testing a particular within-category hypothesis (as in **Suppl. Table 7**). All LME models were fit with maximum likelihood estimation and additionally included ‘Participant’ and ‘ROI’ (for the five ‘core’ left hemisphere language regions, <u>Methods</u>) as a random effect. See **Suppl. Table 7** and <u>Methods</u> for additional model details. Bolded terms are significant (p<0.05).

|  | *Effect* | *ROI* | *Estimate* | *Std..Error* | *df* | *t.value* | *P.uncorrected* |
| --- | --- | --- | --- | --- | --- | --- | --- |
| *EffectArticLoc_P* | Articulation_vs_Tapping | Right_CrusI_II_VIIb | 0.02342 | 0.1584 | 33 | 0.1479 | 0.8833 |
| ***ConditionHard*** | **Hard_vs_Easy_executive_function_tasks** | **Right_CrusI_II_VIIb** | **-0.06365** | **0.008819** | **857.6** | **-7.217** | **1.167e-12** |
| ***ConditionStructured*** | **Structured_vs_Unstructured_music_perception** | **Right_CrusI_II_VIIb** | **-0.05947** | **0.02964** | **89.01** | **-2.006** | **0.04787** |
| *ConditionSocial* | Social_vs_Non-social_high_level_visual_perception | Right_CrusI_II_VIIb | 0.04032 | 0.0292 | 196 | 1.381 | 0.1689 |
| *EffectCloudy_Mental* | Mental_vs_Physical_visual_event_semantics | Right_CrusI_II_VIIb | -0.01769 | 0.05208 | 46 | -0.3397 | 0.7356 |
| ***EffectCloudy_Social*** | **Social_vs_Pain_visual_event_semantics** | **Right_CrusI_II_VIIb** | **0.2523** | **0.0584** | **46** | **4.32** | **8.258e-05** |
| ***EffectPic_Sem*** | **Semantic_vs_Perceptual_visual_task** | **Right_CrusI_II_VIIb** | **0.2577** | **0.06164** | **49.8** | **4.18** | **0.0001175** |

**Supplemental Table 20:**
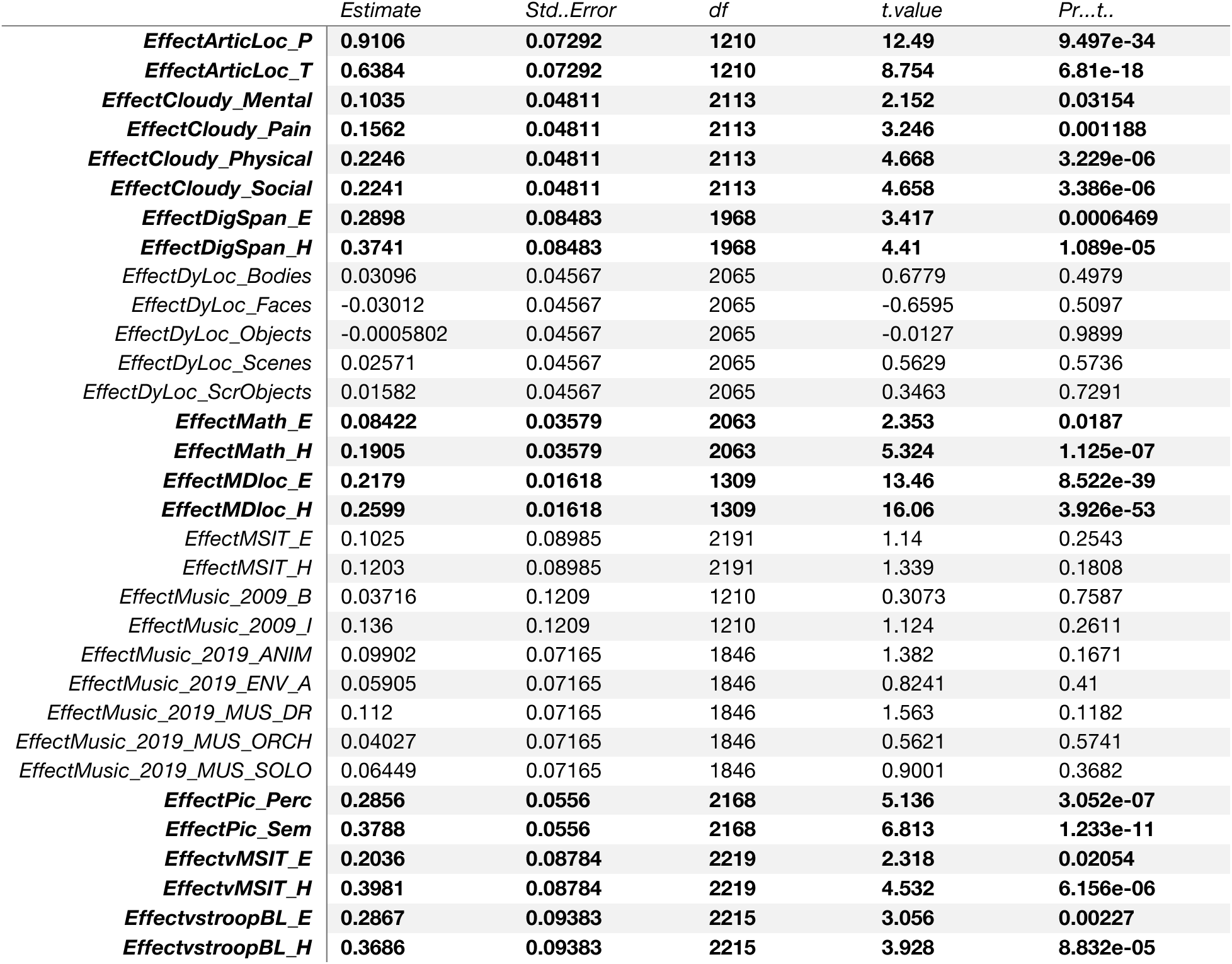
LangCereb4 (right VIIb/VIIIa)’s non-linguistic responses relative to baseline. A linear mixed-effects (LME) model was used to evaluate responses in LangCereb4 to the non-linguistic tasks (Expts. 2a-e, **Fig. 3e**) relative to a fixation baseline, as in **Suppl. Table 8**. Bolded terms are significant (p<0.05).

|  | *Estimate* | *Std..Error* | *df* | *t.value* | *Pr...t..* |
| --- | --- | --- | --- | --- | --- |
| ***EffectArticLoc_P*** | **0.9106** | **0.07292** | **1210** | **12.49** | **9.497e-34** |
| ***EffectArticLoc_T*** | **0.6384** | **0.07292** | **1210** | **8.754** | **6.81e-18** |
| ***EffectCloudy_Mental*** | **0.1035** | **0.04811** | **2113** | **2.152** | **0.03154** |
| ***EffectCloudy_Pain*** | **0.1562** | **0.04811** | **2113** | **3.246** | **0.001188** |
| ***EffectCloudy_Physical*** | **0.2246** | **0.04811** | **2113** | **4.668** | **3.229e-06** |
| ***EffectCloudy_Social*** | **0.2241** | **0.04811** | **2113** | **4.658** | **3.386e-06** |
| ***EffectDigSpan_E*** | **0.2898** | **0.08483** | **1968** | **3.417** | **0.0006469** |
| ***EffectDigSpan_H*** | **0.3741** | **0.08483** | **1968** | **4.41** | **1.089e-05** |
| *EffectDyLoc_Bodies* | 0.03096 | 0.04567 | 2065 | 0.6779 | 0.4979 |
| *EffectDyLoc_Faces* | -0.03012 | 0.04567 | 2065 | -0.6595 | 0.5097 |
| *EffectDyLoc_Objects* | -0.0005802 | 0.04567 | 2065 | -0.0127 | 0.9899 |
| *EffectDyLoc_Scenes* | 0.02571 | 0.04567 | 2065 | 0.5629 | 0.5736 |
| *EffectDyLoc_ScrObjects* | 0.01582 | 0.04567 | 2065 | 0.3463 | 0.7291 |
| ***EffectMath_E*** | **0.08422** | **0.03579** | **2063** | **2.353** | **0.0187** |
| ***EffectMath_H*** | **0.1905** | **0.03579** | **2063** | **5.324** | **1.125e-07** |
| ***EffectMDloc_E*** | **0.2179** | **0.01618** | **1309** | **13.46** | **8.522e-39** |
| ***EffectMDloc_H*** | **0.2599** | **0.01618** | **1309** | **16.06** | **3.926e-53** |
| *EffectMSIT_E* | 0.1025 | 0.08985 | 2191 | 1.14 | 0.2543 |
| *EffectMSIT_H* | 0.1203 | 0.08985 | 2191 | 1.339 | 0.1808 |
| *EffectMusic_2009_B* | 0.03716 | 0.1209 | 1210 | 0.3073 | 0.7587 |
| *EffectMusic_2009_I* | 0.136 | 0.1209 | 1210 | 1.124 | 0.2611 |
| *EffectMusic_2019_ANIM* | 0.09902 | 0.07165 | 1846 | 1.382 | 0.1671 |
| *EffectMusic_2019_ENV_A* | 0.05905 | 0.07165 | 1846 | 0.8241 | 0.41 |
| *EffectMusic_2019_MUS_DR* | 0.112 | 0.07165 | 1846 | 1.563 | 0.1182 |
| *EffectMusic_2019_MUS_ORCH* | 0.04027 | 0.07165 | 1846 | 0.5621 | 0.5741 |
| *EffectMusic_2019_MUS_SOLO* | 0.06449 | 0.07165 | 1846 | 0.9001 | 0.3682 |
| ***EffectPic_Perc*** | **0.2856** | **0.0556** | **2168** | **5.136** | **3.052e-07** |
| ***EffectPic_Sem*** | **0.3788** | **0.0556** | **2168** | **6.813** | **1.233e-11** |
| ***EffectvMSIT_E*** | **0.2036** | **0.08784** | **2219** | **2.318** | **0.02054** |
| ***EffectvMSIT_H*** | **0.3981** | **0.08784** | **2219** | **4.532** | **6.156e-06** |
| ***EffectvstroopBL_E*** | **0.2867** | **0.09383** | **2215** | **3.056** | **0.00227** |
| ***EffectvstroopBL_H*** | **0.3686** | **0.09383** | **2215** | **3.928** | **8.832e-05** |

**Supplemental Table 21:**
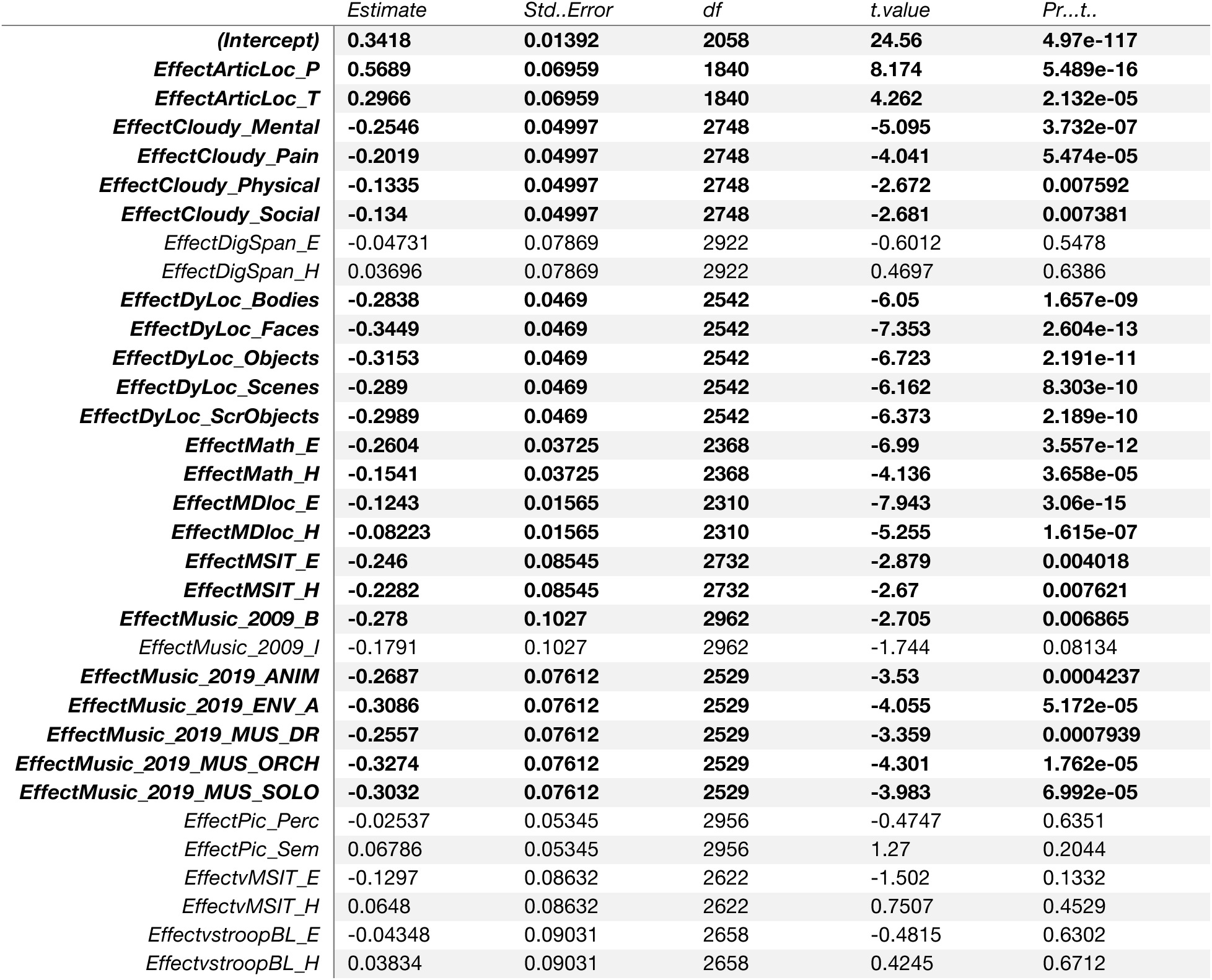
LangCereb4 (right VIIb/VIIIa)’s non-linguistic responses relative to the control condition from the language localizer. A linear mixed-effects (LME) model was used to evaluate responses in LangCereb4 to the non-linguistic tasks (Expts. 2a-e, **Fig. 3e**) relative to the control condition from the language localizer, as in **Suppl. Table 9**. Bolded terms are significant (p<0.05).

**Supplemental Table 22:** LangCereb4 (right VIIb/VIIIa)’s non-linguistic responses relative to language. A linear mixed-effects (LME) model was used to evaluate responses in LangCereb4 to the non-linguistic tasks (Expts. 2a-e, **Fig. 3e**) relative to language, as in **Suppl. Table 9**. Bolded terms are significant (p<0.05).

|  | Estimate | Std..Error | df | t.value | Pr...t.. |
| --- | --- | --- | --- | --- | --- |
| **(Intercept)** | **0.6368** | **0.01483** | **2150** | **42.94** | **1.719e-291** |
| **EffectArticLoc_P** | **0.2738** | **0.07393** | **1929** | **3.704** | **0.0002184** |
| EffectArticLoc_T | 0.001543 | 0.07393 | 1929 | 0.02086 | 0.9834 |
| **EffectCloudy_Mental** | **-0.5719** | **0.05456** | **2784** | **-10.48** | **3.066e-25** |
| **EffectCloudy_Pain** | **-0.5192** | **0.05456** | **2784** | **-9.517** | **3.735e-21** |
| **EffectCloudy_Physical** | **-0.4508** | **0.05456** | **2784** | **-8.263** | **2.176e-16** |
| **EffectCloudy_Social** | **-0.4513** | **0.05456** | **2784** | **-8.271** | **2.027e-16** |
| **EffectDigSpan_E** | **-0.2981** | **0.08557** | **2937** | **-3.484** | **0.0005011** |
| **EffectDigSpan_H** | **-0.2139** | **0.08557** | **2937** | **-2.499** | **0.0125** |
| **EffectDyLoc_Bodies** | **-0.5644** | **0.05138** | **2578** | **-10.98** | **1.822e-27** |
| **EffectDyLoc_Faces** | **-0.6255** | **0.05138** | **2578** | **-12.17** | **3.442e-33** |
| **EffectDyLoc_Objects** | **-0.5959** | **0.05138** | **2578** | **-11.6** | **2.344e-30** |
| **EffectDyLoc_Scenes** | **-0.5696** | **0.05138** | **2578** | **-11.09** | **6.15e-28** |
| **EffectDyLoc_ScrObjects** | **-0.5795** | **0.05138** | **2578** | **-11.28** | **7.769e-29** |
| **EffectMath_E** | **-0.5695** | **0.04091** | **2392** | **-13.92** | **2.1e-42** |
| **EffectMath_H** | **-0.4631** | **0.04091** | **2392** | **-11.32** | **5.626e-29** |
| **EffectMDloc_E** | **-0.4244** | **0.0172** | **2310** | **-24.67** | **1.715e-119** |
| **EffectMDloc_H** | **-0.3824** | **0.0172** | **2310** | **-22.23** | **2.354e-99** |
| **EffectMSIT_E** | **-0.4869** | **0.09332** | **2765** | **-5.218** | **1.944e-07** |
| **EffectMSIT_H** | **-0.4691** | **0.09332** | **2765** | **-5.027** | **5.31e-07** |
| **EffectMusic_2009_B** | **-0.5619** | **0.1112** | **2954** | **-5.054** | **4.585e-07** |
| **EffectMusic_2009_I** | **-0.4631** | **0.1112** | **2954** | **-4.165** | **3.197e-05** |
| **EffectMusic_2019_ANIM** | **-0.5693** | **0.0834** | **2566** | **-6.826** | **1.085e-11** |
| **EffectMusic_2019_ENV_A** | **-0.6093** | **0.0834** | **2566** | **-7.305** | **3.675e-13** |
| **EffectMusic_2019_MUS_DR** | **-0.5563** | **0.0834** | **2566** | **-6.67** | **3.114e-11** |
| **EffectMusic_2019_MUS_ORCH** | **-0.628** | **0.0834** | **2566** | **-7.53** | **6.964e-14** |
| **EffectMusic_2019_MUS_SOLO** | **-0.6038** | **0.0834** | **2566** | **-7.24** | **5.9e-13** |
| **EffectPic_Perc** | **-0.3194** | **0.05803** | **2965** | **-5.503** | **4.049e-08** |
| **EffectPic_Sem** | **-0.2261** | **0.05803** | **2965** | **-3.897** | **9.971e-05** |
| **EffectvMSIT_E** | **-0.3812** | **0.09445** | **2648** | **-4.036** | **5.592e-05** |
| **EffectvMSIT_H** | **-0.1867** | **0.09445** | **2648** | **-1.977** | **0.04814** |
| **EffectvstroopBL_E** | **-0.2879** | **0.09876** | **2689** | **-2.916** | **0.003579** |
| **EffectvstroopBL_H** | **-0.2061** | **0.09876** | **2689** | **-2.087** | **0.03697** |

**Supplemental Table 23:**
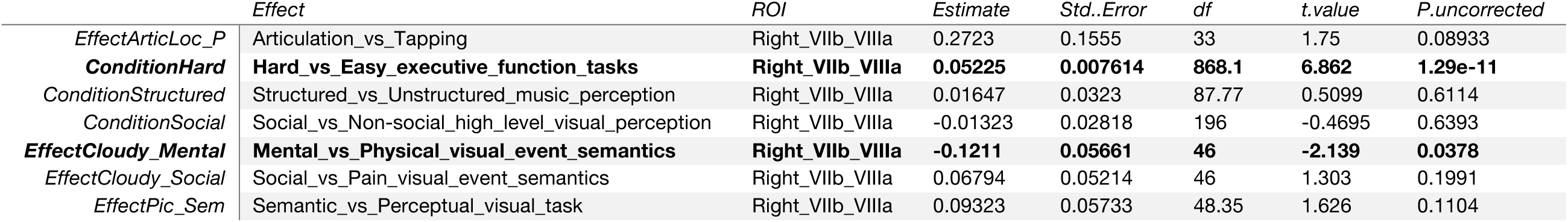
LangCereb4 (right VIIb/VIIIa)’s responses within non-linguistic categories. Linear mixed-effects (LME) models were used to evaluate within-category hypotheses in LangCereb4 (**Fig. 3e**). Each row corresponds to a term from an LME model testing a particular within-category hypothesis (as in **Suppl. Table 7**). All LME models were fit with maximum likelihood estimation and additionally included ‘Participant’ and ‘ROI’ (for the five ‘core’ left hemisphere language regions, <u>Methods</u>) as a random effect. See **Suppl. Table 7** and <u>Methods</u> for additional model details. Bolded terms are significant (p<0.05).

**Supplemental Table 24:**
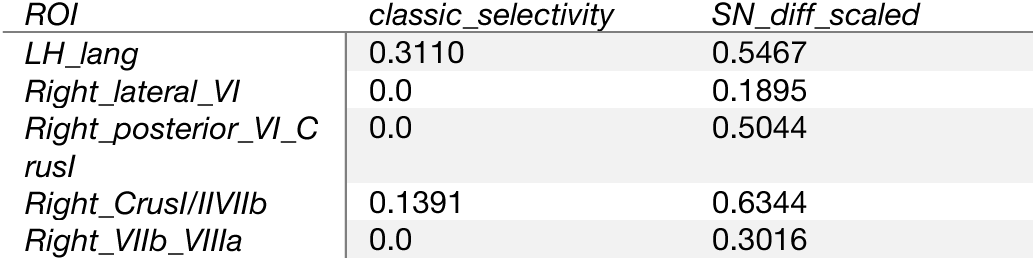
Selectivity for language in the neocortical and cerebellar language regions. Selectivity for language in the neocortical and cerebellar language regions. ‘classic_selectivity’ corresponds to the difference between language and the next highest non-linguistic condition, scaled by the magnitude of the two responses. We additionally report the difference to language and the control condition from the language localizer, scaled by the magnitude of the two responses, for each region.

**Supplemental Table 25:** Dissociation of cerebellar language and Theory of Mind (ToM) regions. Linear mixed-effects (LME) models were used to evaluate if the language and Theory of Mind (ToM) networks are dissociable in the cerebellum (Expt. 2f, **Suppl. Fig. 6e-g**). Three regions of the cerebellum were tested (right VI/Crus I/II/VIIb, left VI/Crus I/II/VIIb, right IX). Each model included three binary, sum-coded fixed effects: ‘Preference’ (0.5 the preferred conditions, language and false belief vignettes; −0.5 for non-preferred conditions, the control condition from the language localizer and false photo vignettes), ‘Response’ (0.5 for the responses to the language localizer, −0.5 for the responses to the ToM localizer), and ‘Localizer’ (0.5 for when the language localizer was used to identify voxels, −0.5 for when the ToM localizer was used). The model included each of these individual terms and their interactions. A separate model was fit per region. The LME models were fit with maximum likelihood estimation and included ‘Participant’ as a random effect. Bolded terms are significant (p < 0.05).

|  | ROI | Estimate | Std..Error | df | t.value | Pr...t.. |
| --- | --- | --- | --- | --- | --- | --- |
| **(Intercept)** | **Right_VI_CrusI_II_VIIb** | **0.4154** | **0.02062** | **156** | **20.15** | **2.869e-45** |
| **PreferencePref>NonPref** | **Right_VI_CrusI_II_VIIb** | **0.2662** | **0.01768** | **1092** | **15.05** | **1.101e-46** |
| **ExperimentLang>ToM** | **Right_VI_CrusI_II_VIIb** | **-0.2843** | **0.01768** | **1092** | **-16.08** | **2.323e-52** |
| **LocalizerLang>ToM** | **Right_VI_CrusI_II_VIIb** | **0.1217** | **0.01768** | **1092** | **6.88** | **1.006e-11** |
| PreferencePref>NonPref:ExperimentLang>ToM | Right_VI_CrusI_II_VIIb | -0.009497 | 0.03537 | 1092 | -0.2685 | 0.7884 |
| PreferencePref>NonPref:LocalizerLang>ToM | Right_VI_CrusI_II_VIIb | 0.05549 | 0.03537 | 1092 | 1.569 | 0.1169 |
| ExperimentLang>ToM:LocalizerLang>ToM | Right_VI_CrusI_II_VIIb | -0.03869 | 0.03537 | 1092 | -1.094 | 0.2742 |
| **PreferencePref>NonPref:ExperimentLang>ToM:LocalizerLang>ToM** | **Right_VI_CrusI_II_VIIb** | **0.2675** | **0.07074** | **1092** | **3.782** | **0.000164** |
| **(Intercept)** | **Left_VI_CrusI_II_VIIb** | **0.2968** | **0.02138** | **156** | **13.88** | **4.855e-29** |
| **PreferencePref>NonPref** | **Left_VI_CrusI_II_VIIb** | **0.2253** | **0.02343** | **1092** | **9.616** | **4.552e-21** |
| **ExperimentLang>ToM** | **Left_VI_CrusI_II_VIIb** | **-0.2127** | **0.02343** | **1092** | **-9.079** | **4.98e-19** |
| **LocalizerLang>ToM** | **Left_VI_CrusI_II_VIIb** | **0.07316** | **0.02343** | **1092** | **3.123** | **0.001837** |
| **PreferencePref>NonPref:ExperimentLang>ToM** | **Left_VI_CrusI_II_VIIb** | **-0.3319** | **0.04685** | **1092** | **-7.084** | **2.501e-12** |
| PreferencePref>NonPref:LocalizerLang>ToM | Left_VI_CrusI_II_VIIb | -0.04768 | 0.04685 | 1092 | -1.018 | 0.3091 |
| ExperimentLang>ToM:LocalizerLang>ToM | Left_VI_CrusI_II_VIIb | 0.06401 | 0.04685 | 1092 | 1.366 | 0.1721 |
| **PreferencePref>NonPref:ExperimentLang>ToM:LocalizerLang>ToM** | **Left_VI_CrusI_II_VIIb** | **0.3148** | **0.09371** | **1092** | **3.359** | **0.0008079** |
| **(Intercept)** | **Right_IX** | **0.2108** | **0.01343** | **156** | **15.7** | **6.53e-34** |
| **PreferencePref>NonPref** | **Right_IX** | **0.1512** | **0.0125** | **1092** | **12.09** | **1.131e-31** |
| **ExperimentLang>ToM** | **Right_IX** | **-0.2596** | **0.0125** | **1092** | **-20.76** | **5.737e-81** |
| **LocalizerLang>ToM** | **Right_IX** | **0.03138** | **0.0125** | **1092** | **2.51** | **0.01222** |
| **PreferencePref>NonPref:ExperimentLang>ToM** | **Right_IX** | **-0.0551** | **0.02501** | **1092** | **-2.204** | **0.02776** |
| PreferencePref>NonPref:LocalizerLang>ToM | Right_IX | 0.0248 | 0.02501 | 1092 | 0.9917 | 0.3216 |
| ExperimentLang>ToM:LocalizerLang>ToM | Right_IX | 0.009752 | 0.02501 | 1092 | 0.39 | 0.6966 |
| PreferencePref>NonPref:ExperimentLang>ToM:LocalizerLang>ToM | Right_IX | 0.03596 | 0.05001 | 1092 | 0.7191 | 0.4722 |

**Supplemental Table 26:** Dissociation of cerebellar language and Multiple Demand (MD) regions. Linear mixed-effects (LME) models were used to evaluate if the language and ‘Multiple Demand’ (MD) networks are dissociable in the cerebellum (Expt. 2b-ii, **Suppl. Fig. 7e-f**). Two regions of the cerebellum were tested (right VI/Crus I, right Crus II/VIIb/VIIIa). Each model included three binary, sum-coded fixed effects: ‘Preference’ (0.5 the preferred conditions, language and the Hard spatial working memory condition; −0.5 for non-preferred conditions, the control condition from the language localizer and the Easy spatial working memory condition), ‘Response’ (0.5 for the responses to the language localizer, −0.5 for the responses to the MD localizer), and ‘Localizer’ (0.5 for when the language localizer was used to identify voxels, −0.5 for when the MD localizer was used). The model included each of these individual terms and their interactions. A separate model was fit per region. The LME models were fit with maximum likelihood estimation and included ‘Participant’ as a random effect. Bolded terms are significant (p < 0.05).

|  | ROI | Estimate | Std..Error | df | t.value | Pr...t.. |
| --- | --- | --- | --- | --- | --- | --- |
| (Intercept) | Right_VI_CrusI | 0.8372 | 0.01927 | 574 | 43.46 | 1.135e-183 |
| PreferencePref>NonPref | Right_VI_CrusI | 0.2267 | 0.01375 | 4018 | 16.49 | 3.896e-59 |
| ExperimentLang>MD | Right_VI_CrusI | -0.3791 | 0.01375 | 4018 | -27.57 | 2.345e-153 |
| LocalizerLang>MD | Right_VI_CrusI | -0.4362 | 0.01375 | 4018 | -31.71 | 3.368e-197 |
| PreferencePref>NonPref:ExperimentLang>MD | Right_VI_CrusI | -0.2041 | 0.0275 | 4018 | -7.422 | 1.401e-13 |
| PreferencePref>NonPref:LocalizerLang>MD | Right_VI_CrusI | -0.01178 | 0.0275 | 4018 | -0.4284 | 0.6683 |
| ExperimentLang>MD:LocalizerLang>MD | Right_VI_CrusI | 0.338 | 0.0275 | 4018 | 12.29 | 4.249e-34 |
| PreferencePref>NonPref:ExperimentLang>MD:LocalizerLang>MD | Right_VI_CrusI | 0.5116 | 0.05501 | 4018 | 9.3 | 2.255e-20 |
| (Intercept) | Right_CrusII_VIIb_VIIIa | 0.3538 | 0.009427 | 574 | 37.53 | 1.219e-156 |
| PreferencePref>NonPref | Right_CrusII_VIIb_VIIIa | 0.1371 | 0.007189 | 4018 | 19.07 | 1.096e-77 |
| ExperimentLang>MD | Right_CrusII_VIIb_VIIIa | -0.08702 | 0.007189 | 4018 | -12.1 | 3.772e-33 |
| LocalizerLang>MD | Right_CrusII_VIIb_VIIIa | -0.07025 | 0.007189 | 4018 | -9.772 | 2.63e-22 |
| PreferencePref>NonPref:ExperimentLang>MD | Right_CrusII_VIIb_VIIIa | -0.03073 | 0.01438 | 4018 | -2.138 | 0.03261 |
| PreferencePref>NonPref:LocalizerLang>MD | Right_CrusII_VIIb_VIIIa | 0.04774 | 0.01438 | 4018 | 3.32 | 0.0009078 |
| ExperimentLang>MD:LocalizerLang>MD | Right_CrusII_VIIb_VIIIa | 0.547 | 0.01438 | 4018 | 38.05 | 8.611e-271 |
| PreferencePref>NonPref:ExperimentLang>MD:LocalizerLang>MD | Right_CrusII_VIIb_VIIIa | 0.5144 | 0.02876 | 4018 | 17.89 | 6.463e-69 |

**Supplemental Table 27:** Dissociation of cerebellar language and articulation regions. Linear mixed-effects (LME) models were used to evaluate if language and articulation are dissociable in the cerebellum (Expt. 2a, **Suppl. Fig. 9e**). One region of the cerebellum was tested (right VIIb/VIIIa). The model included three binary, sum-coded fixed effects: ‘Preference’ (0.5 the preferred conditions, language and articulation; −0.5 for non-preferred conditions, the control condition from the language localizer and finger tapping), ‘Response’ (0.5 for the responses to the language localizer, −0.5 for the responses to the articulation localizer), and ‘Localizer’ (0.5 for when the language localizer was used to identify voxels, −0.5 for when the articulation localizer was used). The model included each of these individual terms and their interactions. The LME model was fit with maximum likelihood estimation and included ‘Participant’ as a random effect. Bolded terms are significant (p < 0.05).

|  | *ROI* | *Estimate* | *Std..Error* | *df* | *t.value* | *Pr...t..* |
| --- | --- | --- | --- | --- | --- | --- |
| **(Intercept)** | **Right_VIIb/VIIIa** | **0.7844** | **0.07237** | **33** | **10.84** | **2.076e-12** |
| **PreferencePref>NonPref** | **Right_VIIb/VIIIa** | **0.4746** | **0.06161** | **231** | **7.703** | **3.879e-13** |
| **ExperimentLang>Artic** | **Right_VIIb/VIIIa** | **-0.1387** | **0.06161** | **231** | **-2.251** | **0.02535** |
| LocalizerLang>Artic | Right_VIIb/VIIIa | 0.06423 | 0.06161 | 231 | 1.042 | 0.2983 |
| PreferencePref>NonPref:ExperimentLang>Artic | Right_VIIb/VIIIa | -0.169 | 0.1232 | 231 | -1.371 | 0.1716 |
| PreferencePref>NonPref:LocalizerLang>Artic | Right_VIIb/VIIIa | -0.06245 | 0.1232 | 231 | -0.5068 | 0.6128 |
| **ExperimentLang>Artic:LocalizerLang>Artic** | **Right_VIIb/VIIIa** | **0.2946** | **0.1232** | **231** | **2.391** | **0.01762** |
| **PreferencePref>NonPref:ExperimentLang>Artic:LocalizerLang>Artic** | **Right_VIIb/VIIIa** | **0.8781** | **0.2465** | **231** | **3.563** | **0.0004455** |

**Supplemental Table 28:** The neocortical language network’s responses during language comprehension and production. Linear mixed-effects (LME) models were used to evaluate the responses in the neocortical language network during language comprehension and production (Expt. 3a-b, **Fig. 4c**). The responses to individual conditions were evaluated relative to a fixation baseline, the Nonword-list condition, and the Sentence condition (both in the respective modality). LME models were fit as in **Suppl. Tables 4-6**. Bolded terms are significant (p < 0.05).

|  | *Experiment* | *Reference* | *Estimate* | *Std..Error* | *df* | *t.value* | *Pr...t..* |
| --- | --- | --- | --- | --- | --- | --- | --- |
| **EffectJ** | **comprehension** | **baseline** | **1.029** | **0.2122** | **5.138** | **4.851** | **0.00434** |
| *EffectN* | comprehension | baseline | 0.5295 | 0.2122 | 5.138 | 2.496 | 0.05345 |
| **EffectS** | **comprehension** | **baseline** | **1.797** | **0.2122** | **5.138** | **8.471** | **0.0003289** |
| **EffectW** | **comprehension** | **baseline** | **0.9664** | **0.2122** | **5.138** | **4.555** | **0.00569** |
| **EffectJ** | **comprehension** | **N** | **0.4996** | **0.05936** | **1342** | **8.417** | **9.783e-17** |
| **EffectS** | **comprehension** | **N** | **1.268** | **0.05936** | **1342** | **21.36** | **2.414e-87** |
| **EffectW** | **comprehension** | **N** | **0.4368** | **0.05936** | **1342** | **7.359** | **3.215e-13** |
| **EffectJ** | **comprehension** | **S** | **-0.768** | **0.05936** | **1342** | **-12.94** | **3.592e-36** |
| **EffectN** | **comprehension** | **S** | **-1.268** | **0.05936** | **1342** | **-21.36** | **2.414e-87** |
| **EffectW** | **comprehension** | **S** | **-0.8308** | **0.05936** | **1342** | **-14** | **1.191e-41** |
| **EffectNProd** | **production_spoken** | **baseline** | **1.463** | **0.5156** | **6.659** | **2.837** | **0.02653** |
| **EffectSProd** | **production_spoken** | **baseline** | **2.346** | **0.5156** | **6.659** | **4.55** | **0.002994** |
| **EffectWProd** | **production_spoken** | **baseline** | **1.773** | **0.5156** | **6.659** | **3.438** | **0.01174** |
| **EffectSProd** | **production_spoken** | **NProd** | **0.883** | **0.1558** | **372** | **5.666** | **2.927e-08** |
| **EffectWProd** | **production_spoken** | **NProd** | **0.3101** | **0.1558** | **372** | **1.99** | **0.04735** |
| **EffectNProd** | **production_spoken** | **SProd** | **-0.883** | **0.1558** | **372** | **-5.666** | **2.927e-08** |
| **EffectWProd** | **production_spoken** | **SProd** | **-0.5729** | **0.1558** | **372** | **-3.677** | **0.0002712** |
| *EffectNProd* | production_typed | baseline | 0.3448 | 0.3315 | 16.75 | 1.04 | 0.313 |
| **EffectSProd** | **production_typed** | **baseline** | **1.078** | **0.3315** | **16.75** | **3.253** | **0.004759** |
| *EffectWProd* | production_typed | baseline | 0.2704 | 0.3315 | 16.75 | 0.8156 | 0.4262 |
| **EffectSProd** | **production_typed** | **NProd** | **0.7334** | **0.1745** | **176** | **4.202** | **4.2e-05** |
| *EffectWProd* | production_typed | NProd | -0.07446 | 0.1745 | 176 | -0.4266 | 0.6702 |
| **EffectNProd** | **production_typed** | **SProd** | **-0.7334** | **0.1745** | **176** | **-4.202** | **4.2e-05** |
| **EffectWProd** | **production_typed** | **SProd** | **-0.8079** | **0.1745** | **176** | **-4.629** | **7.123e-06** |

**Supplemental Table 29:** LangCereb1 (right lateral VI)’s responses during language comprehension and production. Linear mixed-effects (LME) models were used to evaluate the responses in LangCereb1 during language comprehension and production (Expt. 3a-b, **Suppl. Fig. 10a**). The responses to individual conditions were evaluated relative to a fixation baseline, the Nonword-list condition, and the Sentence condition (both in the respective modality). LME models were fit as in **Suppl. Tables 8-10**. Bolded terms are significant (p < 0.05).

|  | *Experiment* | *Reference* | *Estimate* | *Std..Error* | *df* | *t.value* | *Pr...t..* |
| --- | --- | --- | --- | --- | --- | --- | --- |
| **EffectJ** | **comprehension** | **baseline** | **0.645** | **0.07841** | **129.5** | **8.226** | **1.774e-13** |
| **EffectN** | **comprehension** | **baseline** | **0.5182** | **0.07841** | **129.5** | **6.608** | **9.191e-10** |
| **EffectS** | **comprehension** | **baseline** | **0.6506** | **0.07841** | **129.5** | **8.297** | **1.206e-13** |
| **EffectW** | **comprehension** | **baseline** | **0.4689** | **0.07841** | **129.5** | **5.98** | **2.042e-08** |
| *EffectJ* | comprehension | N | 0.1269 | 0.06814 | 210 | 1.862 | 0.06398 |
| *EffectS* | comprehension | N | 0.1324 | 0.06814 | 210 | 1.943 | 0.05334 |
| *EffectW* | comprehension | N | -0.04929 | 0.06814 | 210 | -0.7234 | 0.4702 |
| *EffectJ* | comprehension | S | -0.005517 | 0.06814 | 210 | -0.08097 | 0.9355 |
| *EffectN* | comprehension | S | -0.1324 | 0.06814 | 210 | -1.943 | 0.05334 |
| **EffectW** | **comprehension** | **S** | **-0.1817** | **0.06814** | **210** | **-2.667** | **0.00826** |
| **EffectNProd** | **production_spoken** | **baseline** | **1.466** | **0.2268** | **36.08** | **6.463** | **1.664e-07** |
| **EffectSProd** | **production_spoken** | **baseline** | **1.663** | **0.2268** | **36.08** | **7.331** | **1.196e-08** |
| **EffectWProd** | **production_spoken** | **baseline** | **1.685** | **0.2268** | **36.08** | **7.427** | **8.982e-09** |
| *EffectSProd* | production_spoken | NProd | 0.1971 | 0.1565 | 52 | 1.26 | 0.2135 |
| *EffectWProd* | production_spoken | NProd | 0.2188 | 0.1565 | 52 | 1.398 | 0.1679 |
| *EffectNProd* | production_spoken | SProd | -0.1971 | 0.1565 | 52 | -1.26 | 0.2135 |
| *EffectWProd* | production_spoken | SProd | 0.02172 | 0.1565 | 52 | 0.1388 | 0.8901 |
| **EffectNProd** | **production_typed** | **baseline** | **1.527** | **0.3883** | **14.14** | **3.933** | **0.001476** |
| **EffectSProd** | **production_typed** | **baseline** | **1.628** | **0.3883** | **14.14** | **4.193** | **0.0008845** |
| **EffectWProd** | **production_typed** | **baseline** | **1.629** | **0.3883** | **14.14** | **4.195** | **0.0008819** |
| *EffectSProd* | production_typed | NProd | 0.1011 | 0.1908 | 24 | 0.53 | 0.601 |
| *EffectWProd* | production_typed | NProd | 0.1017 | 0.1908 | 24 | 0.533 | 0.5989 |
| *EffectNProd* | production_typed | SProd | -0.1011 | 0.1908 | 24 | -0.53 | 0.601 |
| *EffectWProd* | production_typed | SProd | 0.0005857 | 0.1908 | 24 | 0.00307 | 0.9976 |

**Supplemental Table 30:** LangCereb2 (right posterior VI/Crus I)’s responses during language comprehension and production. Linear mixed-effects (LME) models were used to evaluate the responses in LangCereb2 during language comprehension and production (Expt. 3a-b, **Suppl. Fig. 10b**). The responses to individual conditions were evaluated relative to a fixation baseline, the Nonword-list condition, and the Sentence condition (both in the respective modality). LME models were fit as in **Suppl. Tables 8-10**. Bolded terms are significant (p < 0.05).

|  | *Experiment* | *Reference* | *Estimate* | *Std..Error* | *df* | *t.value* | *Pr...t..* |
| --- | --- | --- | --- | --- | --- | --- | --- |
| **EffectJ** | **comprehension** | **baseline** | **0.2008** | **0.04444** | **117** | **4.518** | **1.503e-05** |
| **EffectN** | **comprehension** | **baseline** | **0.1703** | **0.04444** | **117** | **3.832** | **0.0002062** |
| **EffectS** | **comprehension** | **baseline** | **0.38** | **0.04444** | **117** | **8.55** | **5.467e-14** |
| **EffectW** | **comprehension** | **baseline** | **0.1753** | **0.04444** | **117** | **3.945** | **0.0001367** |
| *EffectJ* | comprehension | N | 0.03048 | 0.03547 | 210 | 0.8595 | 0.391 |
| **EffectS** | **comprehension** | **N** | **0.2097** | **0.03547** | **210** | **5.912** | **1.354e-08** |
| *EffectW* | comprehension | N | 0.005015 | 0.03547 | 210 | 0.1414 | 0.8877 |
| **EffectJ** | **comprehension** | **S** | **-0.1792** | **0.03547** | **210** | **-5.052** | **9.471e-07** |
| **EffectN** | **comprehension** | **S** | **-0.2097** | **0.03547** | **210** | **-5.912** | **1.354e-08** |
| **EffectW** | **comprehension** | **S** | **-0.2046** | **0.03547** | **210** | **-5.77** | **2.81e-08** |
| *EffectNProd* | production_spoken | baseline | 0.1627 | 0.1279 | 62.54 | 1.272 | 0.208 |
| **EffectSProd** | **production_spoken** | **baseline** | **0.5596** | **0.1279** | **62.54** | **4.377** | **4.659e-05** |
| **EffectWProd** | **production_spoken** | **baseline** | **0.4391** | **0.1279** | **62.54** | **3.434** | **0.001061** |
| **EffectSProd** | **production_spoken** | **NProd** | **0.397** | **0.1456** | **52** | **2.726** | **0.008712** |
| *EffectWProd* | production_spoken | NProd | 0.2764 | 0.1456 | 52 | 1.898 | 0.06323 |
| **EffectNProd** | **production_spoken** | **SProd** | **-0.397** | **0.1456** | **52** | **-2.726** | **0.008712** |
| *EffectWProd* | production_spoken | SProd | -0.1206 | 0.1456 | 52 | -0.828 | 0.4114 |
| **EffectNProd** | **production_typed** | **baseline** | **0.6099** | **0.2454** | **13.52** | **2.486** | **0.02671** |
| **EffectSProd** | **production_typed** | **baseline** | **0.7901** | **0.2454** | **13.52** | **3.22** | **0.00642** |
| **EffectWProd** | **production_typed** | **baseline** | **0.6697** | **0.2454** | **13.52** | **2.729** | **0.01672** |
| *EffectSProd* | production_typed | NProd | 0.1801 | 0.1031 | 24 | 1.747 | 0.09338 |
| *EffectWProd* | production_typed | NProd | 0.05971 | 0.1031 | 24 | 0.5793 | 0.5678 |
| *EffectNProd* | production_typed | SProd | -0.1801 | 0.1031 | 24 | -1.747 | 0.09338 |
| *EffectWProd* | production_typed | SProd | -0.1204 | 0.1031 | 24 | -1.168 | 0.2543 |

**Supplemental Table 31:** LangCereb3 (right Crus I/II/VIIb)’s responses during language comprehension and production. Linear mixed-effects (LME) models were used to evaluate the responses in LangCereb3 during language comprehension and production (Expt. 3a-b, **Fig. 4d**). The responses to individual conditions were evaluated relative to a fixation baseline, the Nonword-list condition, and the Sentence condition (both in the respective modality). LME models were fit as in **Suppl. Tables 8-10**. Bolded terms are significant (p < 0.05).

|  | *Experiment* | *Reference* | *Estimate* | *Std..Error* | *df* | *t.value* | *Pr...t..* |
| --- | --- | --- | --- | --- | --- | --- | --- |
| **EffectJ** | **comprehension** | **baseline** | **0.2075** | **0.03955** | **131.1** | **5.248** | **6.028e-07** |
| **EffectN** | **comprehension** | **baseline** | **0.1442** | **0.03955** | **131.1** | **3.646** | **0.0003835** |
| **EffectS** | **comprehension** | **baseline** | **0.4866** | **0.03955** | **131.1** | **12.3** | **1.317e-23** |
| **EffectW** | **comprehension** | **baseline** | **0.1754** | **0.03955** | **131.1** | **4.435** | **1.931e-05** |
| *EffectJ* | *comprehension* | *N* | 0.06336 | 0.03469 | 210 | 1.827 | 0.06918 |
| **EffectS** | **comprehension** | **N** | **0.3424** | **0.03469** | **210** | **9.872** | **3.996e-19** |
| *EffectW* | *comprehension* | *N* | 0.03121 | 0.03469 | 210 | 0.8998 | 0.3692 |
| **EffectJ** | **comprehension** | **S** | **-0.2791** | **0.03469** | **210** | **-8.045** | **6.256e-14** |
| **EffectN** | **comprehension** | **S** | **-0.3424** | **0.03469** | **210** | **-9.872** | **3.996e-19** |
| **EffectW** | **comprehension** | **S** | **-0.3112** | **0.03469** | **210** | **-8.972** | **1.648e-16** |
| *EffectNProd* | *production_spoken* | *baseline* | 0.203 | 0.1368 | 32.14 | 1.484 | 0.1475 |
| **EffectSProd** | **production_spoken** | **baseline** | **0.4856** | **0.1368** | **32.14** | **3.551** | **0.001209** |
| *EffectWProd* | *production_spoken* | *baseline* | 0.215 | 0.1368 | 32.14 | 1.573 | 0.1256 |
| **EffectSProd** | **production_spoken** | **NProd** | **0.2826** | **0.0762** | **52** | **3.709** | **0.0005072** |
| *EffectWProd* | *production_spoken* | *NProd* | 0.01207 | 0.0762 | 52 | 0.1584 | 0.8748 |
| **EffectNProd** | **production_spoken** | **SProd** | **-0.2826** | **0.0762** | **52** | **-3.709** | **0.0005072** |
| **EffectWProd** | **production_spoken** | **SProd** | **-0.2705** | **0.0762** | **52** | **-3.55** | **0.0008268** |
| *EffectNProd* | *production_typed* | *baseline* | 0.2918 | 0.2227 | 13.01 | 1.31 | 0.2128 |
| *EffectSProd* | *production_typed* | *baseline* | 0.4759 | 0.2227 | 13.01 | 2.137 | 0.05216 |
| *EffectWProd* | *production_typed* | *baseline* | 0.2511 | 0.2227 | 13.01 | 1.128 | 0.2798 |
| **EffectSProd** | **production_typed** | **NProd** | **0.1841** | **0.07729** | **24** | **2.382** | **0.02547** |
| *EffectWProd* | *production_typed* | *NProd* | -0.04067 | 0.07729 | 24 | -0.5262 | 0.6036 |
| **EffectNProd** | **production_typed** | **SProd** | **-0.1841** | **0.07729** | **24** | **-2.382** | **0.02547** |
| **EffectWProd** | **production_typed** | **SProd** | **-0.2248** | **0.07729** | **24** | **-2.909** | **0.007703** |

**Supplemental Table 32:**
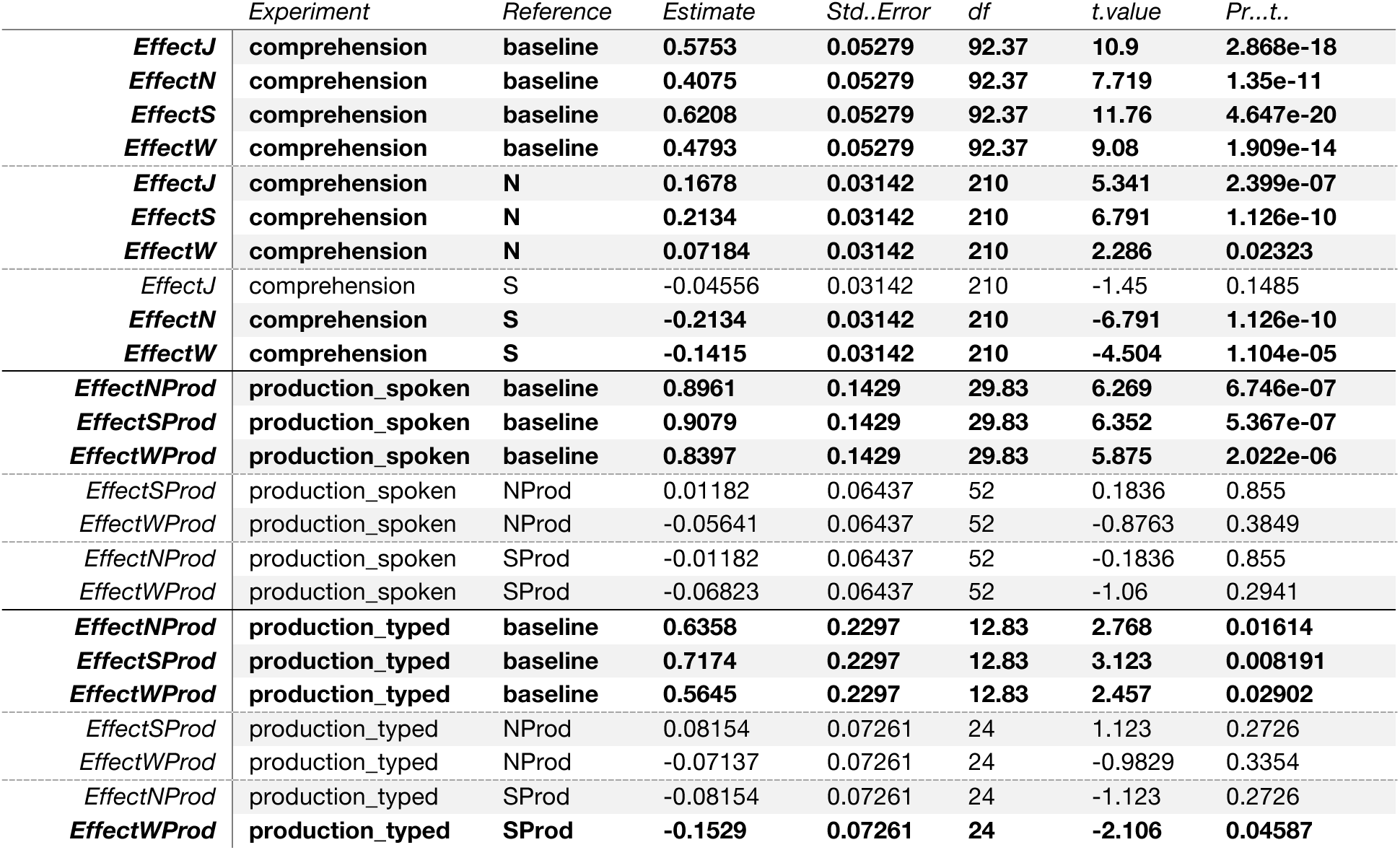
LangCereb4 (right VIIb/VIIIa)’s responses during language comprehension and production. Linear mixed-effects (LME) models were used to evaluate the responses in LangCereb4 during language comprehension and production (Expt. 3a-b, **Suppl. Fig. 10c**). The responses to individual conditions were evaluated relative to a fixation baseline, the Nonword-list condition, and the Sentence condition (both in the respective modality). LME models were fit as in **Suppl. Tables 8-10**. Bolded terms are significant (p < 0.05).

**Supplemental Table 33:** Sensitivity to compositional semantic processing in the neocortical and cerebellar language regions. A linear mixed-effects (LME) models and the likelihood ratio test were used to evaluate whether the neocortical and cerebellar language regions support computations related to semantic composition during language comprehension (Expt. 3a, **Fig. 4c-e**). All four cerebellar language regions reported in **Fig. 1d** were tested. Each model included two binary, treatment-coded fixed effects: ‘lexical’ (1 for conditions that required accessing real lexical items, sentences and word lists; 0 for conditions that did not, Jabberwocky sentences and nonword lists), and ‘syntax’ (1 for conditions that required building syntactic dependency structures, sentences and Jabberwocky sentences; 0 for conditions that did not, word and nonword lists). Two models were fit per region—one with only these fixed effects, and another with these fixed effects plus their interaction. The nested LME models both included ‘Participant’ and ‘Experiment’ (i.e., Expt. 3a-i, ii, or iii) as a random effect, and were fit with maximum likelihood estimation. The models for the neocortical language network additionally included ‘ROI’ as a random intercept (for the five ‘core’ left hemisphere language regions, <u>Methods</u>). Bolded terms indicate a significant improvement in the model performance with the addition of a ‘lexical:syntax’ interaction term via the likelihood ratio test (p < 0.05), suggesting that these regions (the neocortical language network, LangCereb2 (posterior VI/Crus I), and LangCereb3 (Crus I/II/VIIb)) support compositional semantic processing.

|  | ROI | npar | AIC | BIC | logLik | deviance | Chisq | Df | Pr(>Chisq) |
| --- | --- | --- | --- | --- | --- | --- | --- | --- | --- |
| *model_WITHOUT_lexical:syntax* | LH_Language | 7 | 3570 | 3607 | -1778 | 3556 | NA | NA | NA |
| ***model_WITH_lexical:syntax*** | **LH_Language** | **8** | **3556** | **3598** | **-1770** | **3540** | **15.51** | **1** | **8.203e-05** |
| *model_WITHOUT_lexical:syntax* | Right_lateral_VI | 6 | 446.1 | 468 | -217 | 434.1 | NA | NA | NA |
| *model_WITH_lexical:syntax* | Right_lateral_VI | 7 | 447.8 | 473.3 | -216.9 | 433.8 | 0.3279 | 1 | 0.5669 |
| *model_WITHOUT_lexical:syntax* | Right_posterior_VI_CrusI | 6 | 103.2 | 125.1 | -45.6 | 91.2 | NA | NA | NA |
| ***model_WITH_lexical:syntax*** | **Right_posterior_VI_CrusI** | **7** | **93.31** | **118.9** | **-39.65** | **79.31** | **11.89** | **1** | **0.0005636** |
| *model_WITHOUT_lexical:syntax* | Right_CrusI_II_VIIb | 6 | 84.96 | 106.9 | -36.48 | 72.96 | NA | NA | NA |
| ***model_WITH_lexical:syntax*** | **Right_CrusI_II_VIIb** | **7** | **62.53** | **88.07** | **-24.26** | **48.53** | **24.44** | **1** | **7.68e-07** |
| *model_WITHOUT_lexical:syntax* | Right_VIIb_VIIIa | 6 | 73.79 | 95.68 | -30.89 | 61.79 | NA | NA | NA |
| *model_WITH_lexical:syntax* | Right_VIIb_VIIIa | 7 | 75.43 | 101 | -30.72 | 61.43 | 0.3544 | 1 | 0.5516 |

**Supplemental Table 34:** Statistical comparison of linguistic responses in the neocortical language network vs. LangCereb3 (right Crus I/II/VIIb) A linear mixed-effects (LME) model was used to evaluate the sensitivity of the neocortical language regions and CerebLang3 (right Crus I/II/VIIb) to three linguistic processes: lexical semantic processing, syntactic structure building, and combinatorial semantic processing (Expt. 3a, **Fig. 4e**). The model included three binary, treatment-coded fixed effects: ‘Cortex’ (0 for LangCereb3, 1 for the neocortical language network), ‘LexicalSemantics’ (1 for the Word-list and Sentence conditions which requires lexical semantic processing, 0 for the Jabberwocky and Nonword-list conditions which do not), and ‘Syntax’ (1 for the Jabberwocky and Sentence conditions which requires syntactic processing, 0 for the Word-list and Nonword-list conditions which do not). The model included each of these individual terms and their interactions. Only responses during the comprehension experiment (i.e., Expt. 3c) were included in the model. The neocortical responses used in the model were the average over the five ‘core’ left hemisphere language regions. To account for the signal quality difference between the two regions, the BOLD responses were normalized by the average response to the “S” condition for a given region prior to model fitting (<u>Methods</u>). The LME model was fit with restricted maximum likelihood estimation and included ‘Participant’ as a random effect. Bolded terms are significant (p < 0.05).

|  | *Estimate* | *Std..Error* | *df* | *t.value* | *Pr...t..* |
| --- | --- | --- | --- | --- | --- |
| **(Intercept)** | **0.295** | **0.06525** | **276.3** | **4.521** | **9.126e-06** |
| *LexicalSemanticsTRUE* | 0.06935 | 0.07249 | 490 | 0.9566 | 0.3392 |
| *SyntaxTRUE* | 0.1298 | 0.07249 | 490 | 1.791 | 0.07393 |
| *CortexTRUE* | 0.008979 | 0.07249 | 490 | 0.1239 | 0.9015 |
| ***LexicalSemanticsTRUE:SyntaxTRUE*** | **0.5058** | **0.1025** | **490** | **4.934** | **1.106e-06** |
| *LexicalSemanticsTRUE:CortexTRUE* | 0.1807 | 0.1025 | 490 | 1.762 | 0.07863 |
| *SyntaxTRUE:CortexTRUE* | 0.1402 | 0.1025 | 490 | 1.368 | 0.172 |
| ***LexicalSemanticsTRUE:SyntaxTRUE:CortexTRUE*** | **-0.3141** | **0.145** | **490** | **-2.166** | **0.03078** |

**Supplemental Table 35:** The neocortical language network’s modulation by sentence-level linguistic features. Partial correlations of six linguistic features with responses in the neocortical language network to 1,000 diverse sentences (Expt. 3c, n=5, data from Tuckute et al., 2024), controlling for surprisal (extracted from GPT2-xl, Methods). These sentence-level features all measure linguistic processing difficulty (e.g., how frequent the words in the sentence were, whether it was grammatically well-formed, or perceived as frequently occurring, etc.).

|  | *n* | *r* | *CI95%* | *p-val* |
| --- | --- | --- | --- | --- |
| *surprisal_5gram_mean* | 1000 | 0.1272 | [0.07 0.19] | 0.0001 |
| *surprisal_pcfg_mean* | 1000 | 0.0945 | [0.03 0.16] | 0.0028 |
| *rating_gram_mean* | 1000 | -0.2566 | [-0.31 -0.2 ] | 0.0 |
| *rating_sense_mean* | 1000 | -0.2506 | [-0.31 -0.19] | 0.0 |
| *rating_frequency_mean* | 1000 | -0.2767 | [-0.33 -0.22] | 0.0 |
| *rating_conversational_mean* | 1000 | -0.181 | [-0.24 -0.12] | 0.0 |

**Supplemental Table 36:** LangCereb1 (right lateral VI)’s modulation by sentence-level linguistic features. Partial correlations of six linguistic features with responses in LangCereb1 to 1,000 diverse sentences (Expt. 3c, n=5, data from Tuckute et al., 2024), controlling for surprisal (extracted from GPT2-xl, Methods). These sentence-level features all measure linguistic processing difficulty (e.g., how frequent the words in the sentence were, whether it was grammatically well-formed, or perceived as frequently occurring, etc.).

|  | *n* | *r* | *CI*95% | *p-val* |
| --- | --- | --- | --- | --- |
| *surprisal_5gram_mean* | 1000 | 0.034 | [-0.03 0.1 ] | 0.2832 |
| *surprisal_pcfg_mean* | 1000 | 0.0884 | [0.03 0.15] | 0.0052 |
| *rating_gram_mean* | 1000 | -0.1811 | [-0.24 -0.12] | 0.0 |
| *rating_sense_mean* | 1000 | -0.1605 | [-0.22 -0.1 ] | 0.0 |
| *rating_frequency_mean* | 1000 | -0.1327 | [-0.19 -0.07] | 0.0 |
| *rating_conversational_mean* | 1000 | -0.1134 | [-0.17 -0.05] | 0.0003 |

**Supplemental Table 37:** LangCereb2 (right posterior VI/Crus I)’s modulation by sentence-level linguistic features. Partial correlations of six linguistic features with responses in LangCereb2 to 1,000 diverse sentences (Expt. 3c, n=5, data from Tuckute et al., 2024), controlling for surprisal (extracted from GPT2-xl, Methods). These sentence-level features all measure linguistic processing difficulty (e.g., how frequent the words in the sentence were, whether it was grammatically well-formed, or perceived as frequently occurring, etc.).

|  | *n* | *r* | *CI*95% | *p-val* |
| --- | --- | --- | --- | --- |
| *surprisal_5gram_mean* | 1000 | 0.0166 | [-0.05 0.08] | 0.5997 |
| *surprisal_pcfg_mean* | 1000 | 0.054 | [-0.01 0.12] | 0.0881 |
| *rating_gram_mean* | 1000 | -0.191 | [-0.25 -0.13] | 0.0 |
| *rating_sense_mean* | 1000 | -0.1588 | [-0.22 -0.1 ] | 0.0 |
| *rating_frequency_mean* | 1000 | -0.0941 | [-0.16 -0.03] | 0.0029 |
| *rating_conversational_mean* | 1000 | -0.1057 | [-0.17 -0.04] | 0.0008 |

**Supplemental Table 38:** LangCereb3 (right Crus I/II/VIIb)’s modulation by sentence-level linguistic features. Partial correlations of six linguistic features with responses in LangCereb3 to 1,000 diverse sentences (Expt. 3c, n=5, data from Tuckute et al., 2024), controlling for surprisal (extracted from GPT2-xl, Methods). These sentence-level features all measure linguistic processing difficulty (e.g., how frequent the words in the sentence were, whether it was grammatically well-formed, or perceived as frequently occurring, etc.).

|  | *n* | *r* | *CI*95% | *p-val* |
| --- | --- | --- | --- | --- |
| *surprisal_5gram_mean* | 1000 | 0.0439 | [-0.02 0.11] | 0.166 |
| *surprisal_pcfg_mean* | 1000 | 0.0514 | [-0.01 0.11] | 0.1045 |
| ***rating_gram_mean*** | **1000** | **-0.1936** | **[-0.25 -0.13]** | **0.0** |
| ***rating_sense_mean*** | **1000** | **-0.2065** | **[-0.27 -0.15]** | **0.0** |
| ***rating_frequency_mean*** | **1000** | **-0.1792** | **[-0.24 -0.12]** | **0.0** |
| ***rating_conversational_mean*** | **1000** | **-0.1372** | **[-0.2 -0.08]** | **0.0** |

**Supplemental Table 39:** LangCereb4 (right VIIb/VIIIa)’s modulation by sentence-level linguistic features. Partial correlations of six linguistic features with responses in LangCereb4 to 1,000 diverse sentences (Expt. 3c, n=5, data from Tuckute et al., 2024), controlling for surprisal (extracted from GPT2-xl, Methods). These sentence-level features all measure linguistic processing difficulty (e.g., how frequent the words in the sentence were, whether it was grammatically well-formed, or perceived as frequently occurring, etc.).

|  | *n* | *r* | *CI*95% | *p_val* |
| --- | --- | --- | --- | --- |
| *surprisal_5gram_mean* | 1000 | 0.0744 | [0.01 0.14] | 0.0187 |
| *surprisal_pcfg_mean* | 1000 | 0.097 | [0.04 0.16] | 0.0021 |
| ***rating_gram_mean*** | **1000** | **-0.2235** | **[-0.28 -0.16]** | **0.0** |
| ***rating_sense_mean*** | **1000** | **-0.1864** | **[-0.25 -0.13]** | **0.0** |
| ***rating_frequency_mean*** | **1000** | **-0.1769** | **[-0.24 -0.12]** | **0.0** |
| ***rating_conversational_mean*** | **1000** | **-0.1727** | **[-0.23 -0.11]** | **0.0** |

**Supplemental Table 40:** Comparison of functional correlations during resting state vs. story comprehension and between pairs of cerebellar language regions. A linear mixed-effects (LME) model was used to evaluate functional correlations in the cerebellar language regions with the left neocortical language network (**Fig. 6h**). The model included two, treatment-coded fixed effects: ‘cerebROI’ (with four levels corresponding to each of the cerebellar language regions), and ‘dataset’ (1 for functional correlations during a naturalistic story, 0 for functional correlations during resting state). The model included random intercepts for ‘Participant’ and ‘ROI’ (for the five ‘core’ left hemisphere language regions, <u>Methods</u>). The model additionally included random slopes for ‘Participant’ and ‘ROI’ for the ‘dataset’ fixed effect. This random-effects structure was selected by iteratively removing terms from a maximal random-effects structure until the fit was no longer singular. The model was fit with restricted maximum likelihood estimation. The difference in the functional correlations between the two datasets (top), and pairwise comparisons between individual cerebellar regions by dataset (bottom) were estimated using the *‘emmeans’* package. Bolded terms are significant (p<0.05). P-values for the pairwise comparisons (bottom) were adjusted using the Holm correction.

| *contrast* | *dataset* | *estimate* | *SE* | *df* | *t.ratio* | *p.value* |
| --- | --- | --- | --- | --- | --- | --- |
| **resting - story** | **NA** | **-0.09553** | **0.02005** | **10.12** | **-4.763** | **0.0007405** |
| *LangCereb1-LangCereb3* | resting | -0.08333 | 0.008389 | 3216 | -9.933 | 1.909e-22 |
| *LangCereb2-LangCereb3* | resting | -0.06216 | 0.008389 | 3216 | -7.409 | 3.225e-13 |
| *LangCereb4-LangCereb3* | resting | -0.04284 | 0.008389 | 3216 | -5.106 | 3.481e-07 |
| *LangCereb1-LangCereb3* | story | -0.2238 | 0.008389 | 3216 | -26.68 | 2.554e-141 |
| *LangCereb2-LangCereb3* | story | -0.1082 | 0.008389 | 3216 | -12.9 | 3.772e-37 |
| *LangCereb4-LangCereb3* | story | -0.1195 | 0.008389 | 3216 | -14.25 | 1.995e-44 |

**Supplemental Table 41:** Comparison of functional correlations between the right cerebellar language regions and left vs. right hemisphere neocortical language regions. A linear mixed-effects (LME) model was used to evaluate functional correlations in right cerebellar language regions vs. the left and right neocortical language regions (**Fig. 6g**). The model included one binary, treatment-coded fixed effect: ‘hemisphere’ (0 for functional correlations with the left neocortical language network, 1 for functional correlations with the right neocortical language regions). The model included random intercepts for ‘Participant’, ‘cerebROI’ (for the four cerebellar language regions), ‘dataset’ (naturalistic story or resting state), and ‘ROI’ (for the five ‘core’ left hemisphere language regions, <u>Methods</u>). This random-effects structure was selected by iteratively removing terms from a maximal random-effects structure until the fit was no longer singular. The model was fit with restricted maximum likelihood estimation. Bolded terms are significant (p<0.05).

|  | *Estimate* | *Std..Error* | *df* | *t.value* | *Pr...t..* |
| --- | --- | --- | --- | --- | --- |
| (Intercept) | 0.2326 | 0.05786 | 1.938 | 4.02 | 0.05972 |
| **hemisphereRH** | **-0.09661** | **0.02573** | **8** | **-3.755** | **0.005581** |

**Supplemental Table 42:** Comparison of functional correlations between LangCereb3 (Crus I/II/VIIb) and frontal vs. temporal left-hemisphere neocortical language regions. A linear mixed-effects (LME) model was used to evaluate functional correlations between LangCereb3 (Crus I/II/VIIb) and the frontal vs. temporal regions of the ‘core’ neocortical language network (**Fig. 6f**). The model included two binary, treatment-coded fixed effects: ‘lobe’ (0 for frontal language regions, 1 for temporal language regions) and ‘dataset’ (0 for resting state, 1 for the naturalistic story). The model included random intercepts for ‘Participant’ and ‘cortical_region’ (for the five ‘core’ left hemisphere language regions, <u>Methods</u>). This random-effects structure was selected by iteratively removing terms from a maximal random-effects structure until the fit was no longer singular. The model was fit with restricted maximum likelihood estimation. Bolded terms are significant (p<0.05).

|  | *Estimate* | *Std..Error* | *df* | *t.value* | *Pr...t..* |
| --- | --- | --- | --- | --- | --- |
| **(Intercept)** | **0.2212** | **0.01968** | **8.497** | **11.24** | **2.165e-06** |
| *lobeTemporal* | 0.02686 | 0.02543 | 3.846 | 1.056 | 0.3525 |
| **datasetstory** | **0.1349** | **0.01099** | **759** | **12.28** | **9.976e-32** |
| ***lobeTemporal:datasetstory*** | **0.06596** | **0.01738** | **759** | **3.795** | **0.0001595** |

**Supplemental Table 43:**
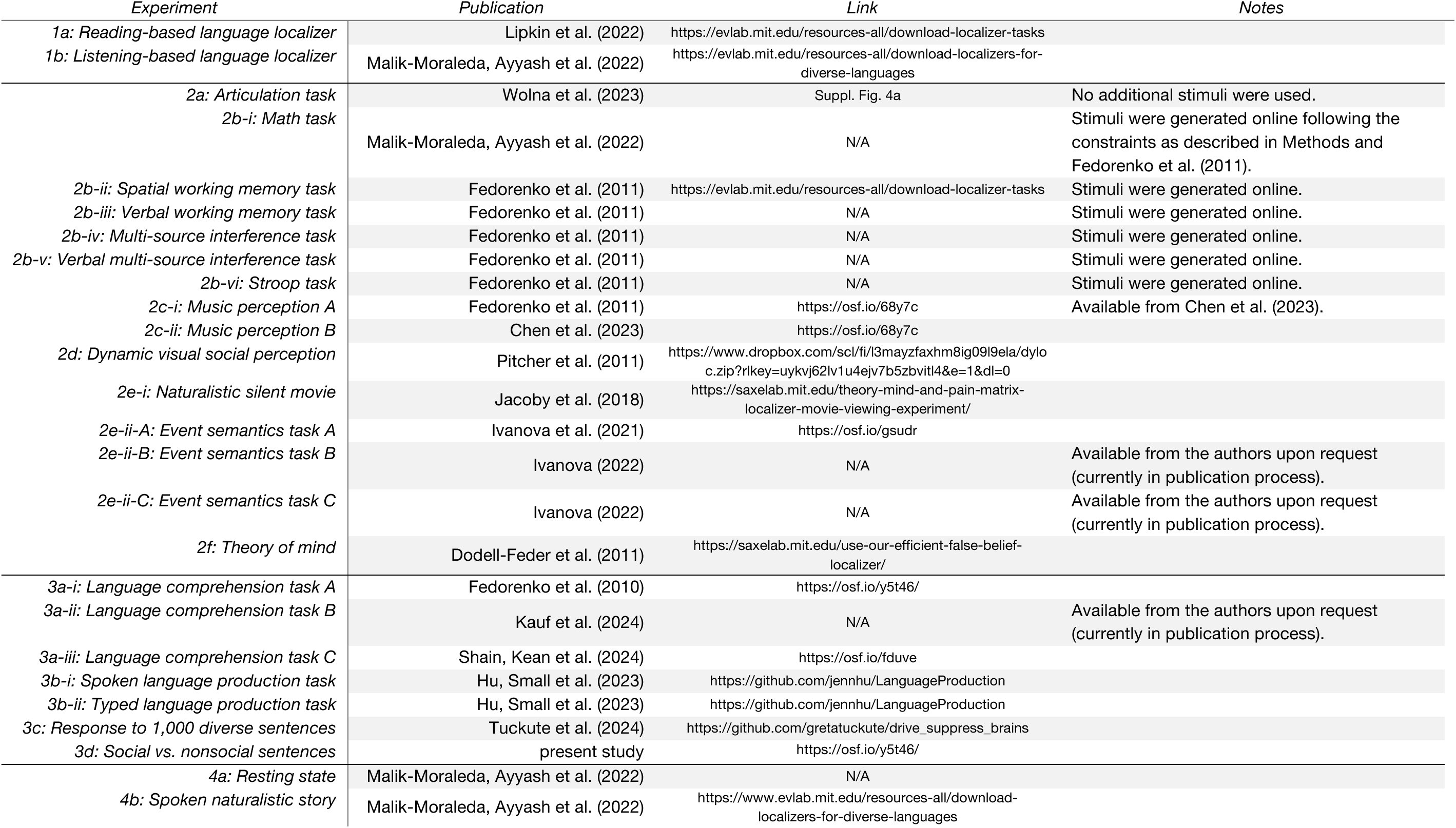
Experimental Stimuli. Links to all experimental stimuli.

